# A nested shell structure coordinates enzyme communication in pyruvate oxidation

**DOI:** 10.64898/2026.05.12.724543

**Authors:** Toni K. Träger, Sourav Maity, Fotis L. Kyrilis, Christian Tüting, Farzad Hamdi, Greg Kafetzopoulos, Z Faidon Brotzakis, Alexander Neuhaus, Anja Blanque, Christos Gatsogiannis, Georgios Skretas, Wouter H. Roos, Panagiotis L. Kastritis

## Abstract

The pyruvate dehydrogenase complex (PDHc)^1^ links glycolysis to the Krebs cycle by catalyzing the oxidative decarboxylation of pyruvate to acetyl-CoA and CO₂, a process essential for life^2,3^. PDHc is formed by structural proteins (E3-binding protein, E3BP)^4–7^, enzymatic subunits (E1, E2, E3)^4,6^, and mobile lipoyl domains (LDs), the latter shuttling intermediates across active sites^6,8^. Although numerous details regarding pyruvate oxidation steps have been elucidated^9^, the precise organization of the entire PDHc remains unknown due to its large size and dynamic heterogeneity. Here, we employ *in silico*, *in vitro*, and *in situ* methods to propose a multi-scale model of PDHc that includes approximately one million atoms and to visualize multiple conformational states. This model reveals a ∼40-50 nm nested shell structure, formed by flexible linkers that spatially coordinate the E1 and E3 enzyme complexes around the E2-E3BP core scaffold. This structure acts as a molecular sieve, selectively guiding lipoyl arms while maintaining enzyme positioning with sub-nm precision. During catalysis, the nested shell structure expands and adopts a mechanically reinforced state comparable in magnitude to viral assemblies^10^. Our findings provide structural context for the textbook “link reaction”^11^, building on decades of biochemical knowledge, are transferable to functional aspects of other α-ketoacid dehydrogenase complexes, and, ultimately, expand our understanding of primary metabolism as a whole.

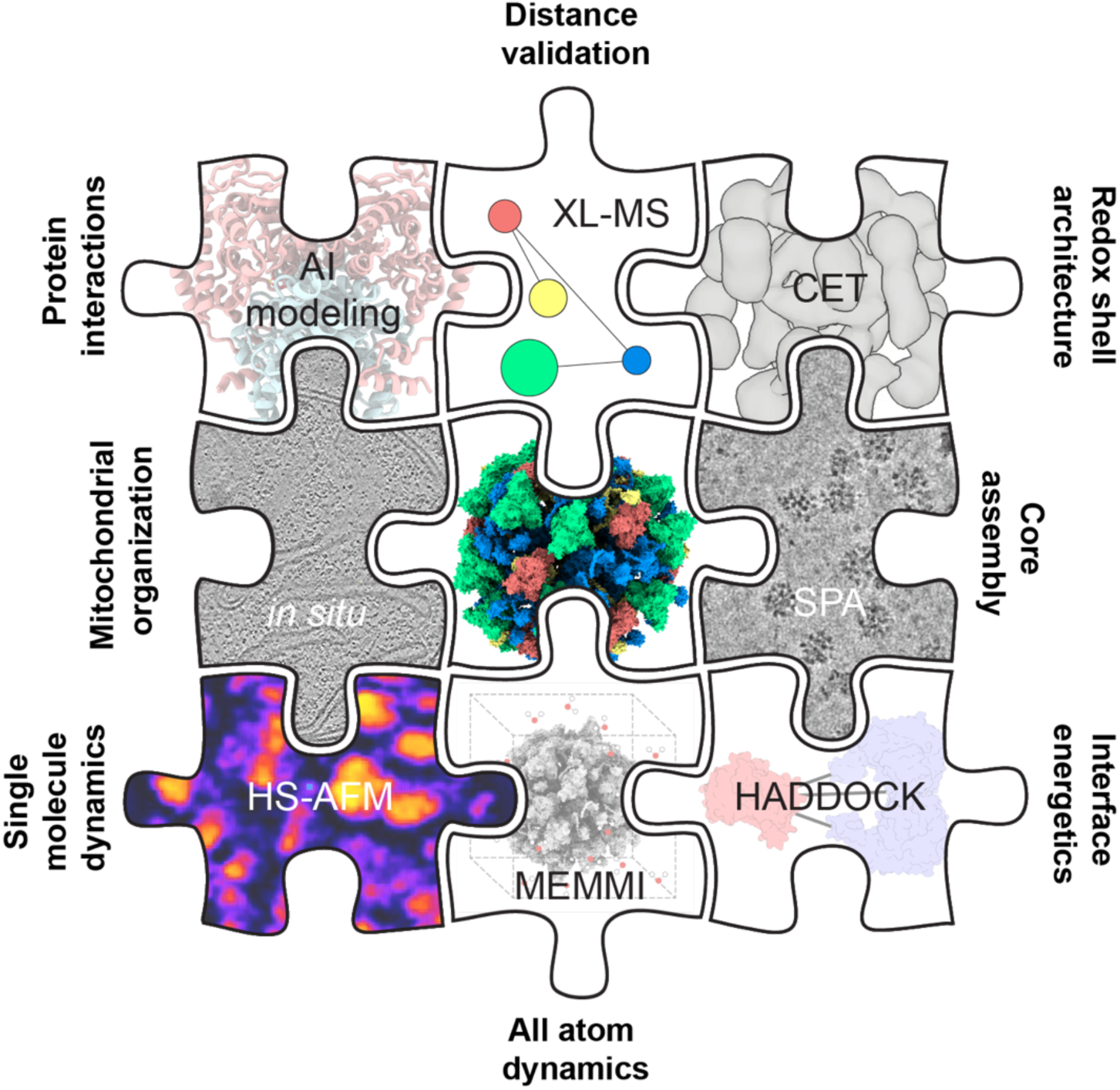

## Main

The pyruvate dehydrogenase complex (PDHc) is the central metabolic checkpoint in all domains of life^3^, linking glycolysis to the tricarboxylic acid (TCA) cycle^1,2^. PDHc catalyzes the oxidative decarboxylation of pyruvate to acetyl-CoA and CO₂, coupled to the reduction of NAD⁺ to NADH, a reaction referred to as the essential “link reaction”^11^. In eukaryotes, PDHc assembles in the mitochondrial matrix as one of the largest soluble enzyme complexes of the cell, supporting substrate channeling across numerous reaction sites^6^. In essence, the complex incorporates three core enzymatic activities: thiamine diphosphate-dependent pyruvate decarboxylation by pyruvate dehydrogenase (E1)^12^, acetyl group transfer to coenzyme A by dihydrolipoamide acetyltransferase (E2), carried by a flexible lipoyl domain (LD), and FAD-dependent reduction by dihydrolipoamide dehydrogenase (E3)^2^, coupled to the reduction of NAD⁺. Each enzyme subunit adopts a multimeric structure with specific stoichiometry. These subunits are brought in proximity through flexible E2 regions and, in eukaryotes, a fourth protein (E3BP)^4,5,7^, which specifically binds E3, stabilizing the complex and influencing regulatory dynamics^13^. Both E3BP and E2 contain flexible, intrinsically disordered regions (IDRs) that are interspersed with peripheral subunit-binding domains (PSBDs), which coordinate the association of E1 and E3, respectively. They also contain subunit-specific LDs that facilitate efficient intermediate transfer^2,9^ and regulate lipoic acid metabolism^14^. This organization allows sequential intermediate shuttling while minimizing substrate loss, ensuring efficient metabolic flux from glycolysis into the TCA cycle. Such elaborate composition and higher-order organization are shared across the larger family of α-ketoacid dehydrogenase complexes^15^. PDHc-mediated pyruvate conversion is tuned by cellular energy and redox state through post-translational modifications (PTMs) and allosteric control^16^. In eukaryotes, PDHc is reversibly phosphorylated by low-abundant E1-specific pyruvate dehydrogenase phosphatases (PDPs) and kinases (PDKs), activating and inactivating the complex, respectively^17^. Additionally, PDHc subunits undergo multiple lysine-directed modifications, including acetylation, succinylation, ubiquitination, and SUMOylation, which modulate enzymatic turnover and availability^18–21^. High ATP/ADP and NADH/NAD⁺ ratios promote PDK-mediated inhibition, whereas PDP activation under low-energy conditions restores flux^16^. In addition, product inhibition by acetyl-CoA and NADH provides rapid feedback at the E2 and E3 steps^15^. These regulatory inputs act on a structurally heterogeneous assembly that contains extensive flexible linkers, raising the question of how enzymatic communication and regulatory accessibility are coordinated across the metabolon. PDHc regulatory mechanisms and structural adaptations within the entire 10-megadalton metabolon are connected. Recent works showed that PDHc stoichiometric variations^4,7,13,22^ and changes in the localization of its embedded biomolecules^4,9,23,24^ may modulate the complex’s conformation^5,6,24^ and its membrane interactions^25^. PDHc’s intricate regulatory architecture balances cellular energy production with resource conservation, sustaining metabolic homeostasis across diverse physiological contexts^26^. PDHc component enzymes further facilitate extensive moonlighting functions^27^ across cellular compartments and regulate organelle-specific pathways, such as amino acid catabolism via the glycine cleavage system or transcription regulation via direct histone acetylation by alternate nuclear entry^28^, possibly sharing similar organizational principles^29^. Ultimately, defects in PDHc are connected to metabolic diseases^30^, the development of aging processes^31^, memory impairment^32^, neurodegeneration^33^, and cancer^34^. However, a comprehensive structural understanding of this highly intricate metabolic supercomplex remains limited by the lack of high-resolution information on the full, endogenous, and conformationally flexible metabolon.

### Architecture of the pyruvate dehydrogenase metabolon

To better understand the molecular basis of pyruvate oxidation, we enriched the native, endogenous PDHc directly from *Chaetomium thermophilum (Thermochaetoides thermophila, T. thermophila)* cultures (Methods). The complex eluted at the expected megadalton size and retained full catalytic activity, as monitored by NADH production (**Extended Data Fig. 1a-b**). Cryo-electron microscopy (cryo-EM) revealed intact PDHc particles with a diameter of 44 ± 2 nm (**N = 100; Fig. 1a-b, Extended Data Fig. 2a-b**), indicating preservation of native architecture under applied biochemical and vitrification conditions^6^. Owing to its size and intrinsic conformational heterogeneity, the PDHc spans structural scales that cannot be captured at uniform resolution by a single reconstruction. Therefore, we first implemented a divide-and-conquer approach (Methods, **Extended Data Fig. 3**), where local masking and subsequent refinement of component subunits are performed.

**Fig. 1:**
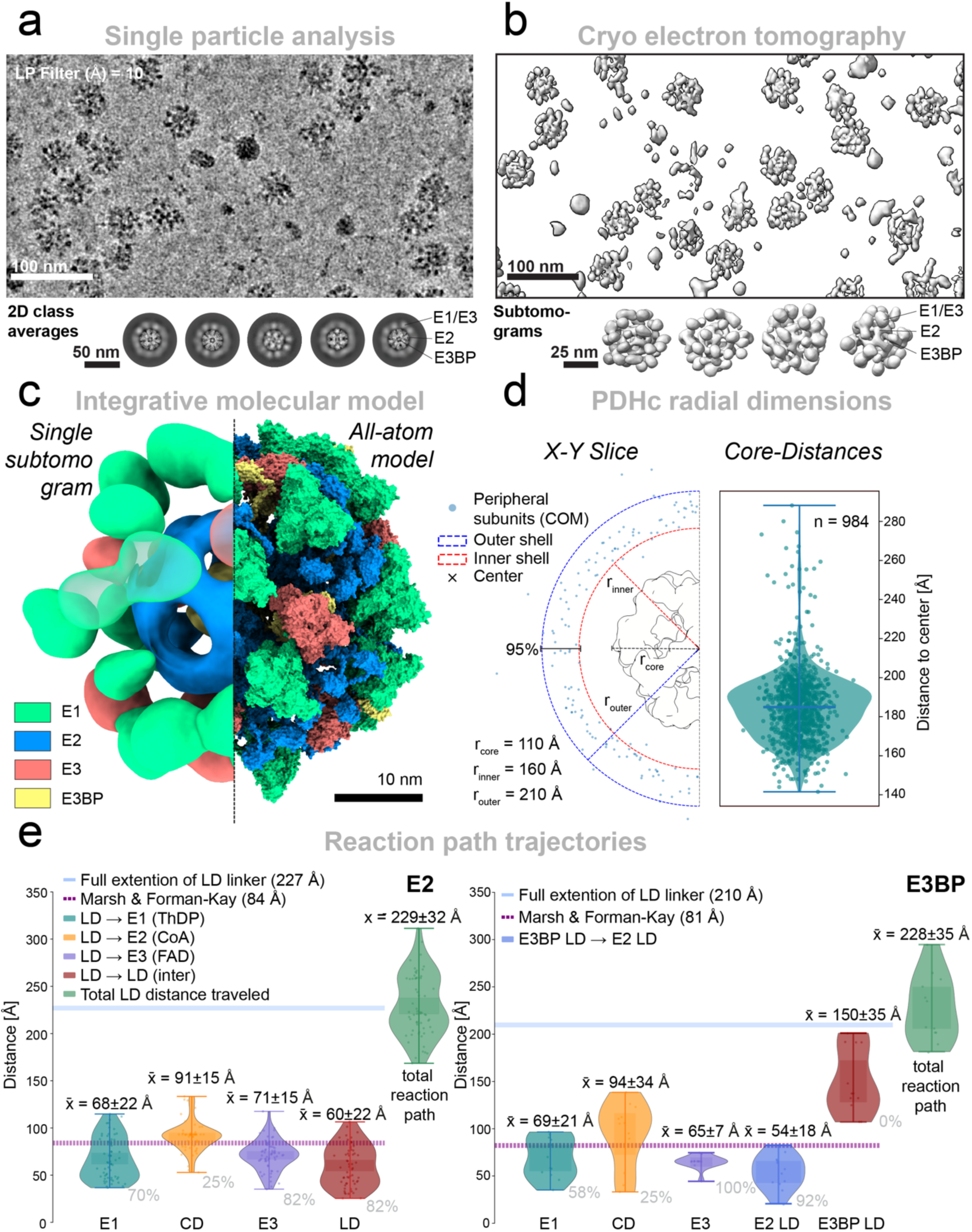
Spatial organization of the pyruvate dehydrogenase complex. **a,** Representative micrograph from the SPA (top) and selected 2D class averages (bottom) display the overall architecture of the PDHc. The core domain containing E2 and E3BP, as well as the peripheral E1 and E3, is annotated. **b,** The reconstructed tomogram (top) of the enriched sample provides additional validation for the spatial confinement and placement of the peripheral subunits. Non-averaged subtomograms show the overall macromolecular assembly of individual PDHc particles (bottom). **c**, Non-averaged subtomogram reflects macromolecular assembly of the native PDHc (left). The all-atom model (right) combines structural information from different experimental and computational levels to create an integrative model for pyruvate oxidative decarboxylation (E1, green; E2, blue; E3, orange; E3BP, yellow). **d**, X–Y slice showing the spatial distribution of the peripheral subunit via the center of mass (COM) points. Dashed circles indicate the inner and outer shell boundaries in which 95% of all peripheral subunits (n = 984) are confined, with a core radius (rcore) of 110 Å, an inner shell radius (rinner) of 160 Å, and an outer shell radius (router) of 210 Å. **e,** To map the reaction distances covered by the LD within the model of the PDHc, distances between reactive LD residues and active site cofactors (E1 ThDP, E2 CoA, and E3 FAD) and inter-LD were calculated. The total LD distance traveled during the complete catalytic cycle is shown in green for both the E2 LD (left) and the E3BP LD (right). Dashed lines indicate the expected mean end-to-end distance of an intrinsically disordered linker (Marsh & Forman-Kay) ^36^, while the solid blue line represents the maximum possible extension of the linker based on a per-residue distance of 3.5 Å. The percentage of feasible active-site distances, determined using the Marsh & Forman-Kay calculated distance, is shown below each distribution.

This approach yielded a 2.84 Å (FSC = 0.143) structure of the native E2 core (**Extended Data Fig. 4-5**), representing the highest-resolution structure of any endogenous or eukaryotic dihydrolipoamide acetyltransferase reported to date (**Extended Data Tab. 1&2**). Focused refinement of the E3BP further resolved two tetrahedral assembly states, each composed of four trimeric E3BP building blocks arranged within the E2 core (**Extended Data Fig. 6-7**). In both configurations, E3BP engages the E2 catalytic domains through conserved electrostatic interactions, forming a rigid internal scaffold while projecting flexible linker regions toward the periphery (**Extended Data Fig. 8-9**). Compared with previously reported fungal PDHc structures, the *T. thermophila* E3BP exhibits an expanded E2-E3BP interface that includes a secondary α-helical contact absent in other systems^7^, strengthening electrostatic coupling and stabilizing the inner core assembly (**Extended Data Fig. 10**).

To define the organization of the intact PDHc, we combined asymmetric single particle analysis (SPA) reconstructions **(Fig. 1a**, **Extended Data Fig. 11**) with the analysis of non-averaged subtomograms from the enriched preparation (**Extended Data Fig. 12**). Specifically, stochastic particle subsampling approximated the posterior distribution of reconstructed cryo-EM maps to identify common density features. Reconstructions without symmetry from randomly subdivided particle sets converged on an architecture comprising a rigid E2-E3BP core surrounded by persistent peripheral densities, demonstrating that the observed organisation is not imposed by symmetry averaging (**Extended Data Fig. 13a**). In all reconstructions, these peripheral densities occupied reproducible spatial zones with E1 preferentially localising near the vertices and E3 near the faces of the dodecahedral scaffold; direct reconstruction of the endogenous E1 heterotetramer from the intact complex validated the predicted architecture and provided an experimental anchor for model integration (**Extended Data Fig. 13b, 14, 15**). The same organisational principles were observed at the level of individual complexes derived from denoised tilt series^35^, where individual subtomograms resolved intact PDHc particles with an icosahedral E2/E3BP core surrounded by clustered peripheral subunits (**Fig. 1b-c, Extended Data Fig. 16a**). Classification of these subtomograms revealed 31-36 peripheral densities per particle, with ∼24 at vertex positions and ∼9 at face positions, consistent with partial but reproducible occupancy of E1 and E3 binding sites (**Extended Data Fig. 16b**). Across both modalities, analysis of peripheral subunit centre-of-mass positions (COM) (n = 987) showed that ∼95% of all E1 and E3 subunits reside within a confined hollow shell with a radius of ∼160 Å and ∼210 Å (**Fig. 1d, Extended Data Fig. 17**), defining a geometrically constrained redox-active compartment surrounding the catalytic core.

Combining the experimentally defined core architecture, peripheral enzyme localisation, and stoichiometric constraints consistent with previous data on PDHc stoichiometry^6^, we modelled the complete metabolon within the reconstructed densities, yielding a composite model for the entire PDHc (**Fig. 1c**). The resulting assembly comprises 60 E2 subunits, 12 E3BP subunits, 20 E1 heterotetramers, and 12 E3 dimers, yielding a total molecular mass of 7.6 MDa and encompassing ∼1.1 million atoms (**Extended Data Tab. 3**). Within this geometry, the majority of productive LD–active-site interactions are spatially feasible (**Fig. 1e**), indicating that catalysis can occur throughout the assembly (**Extended Data Fig. 18**). The peripheral enzymes are arranged in a near-uniform manner, positioning catalytic sites at comparable radial distances and allowing complex-wide turnover. Recurrent peripheral clusters containing both E1 and E3 shorten competent LD trajectories. Consistent with the higher copy number of E2-linked LDs, E3BP-linked LDs are more sparsely distributed and therefore predicted to contribute less frequently to catalytic cycling (**Fig. 1e**).

### Operating principles of pyruvate processing

To understand how the structural features of the PDHc give rise to coordinated enzyme communication, we next analyzed the physical and structural principles that govern organization in the complex’s peripheral regions and its coupling to the E2-E3BP core scaffold. The architectural features of the PDHc that we report imply that coordinated enzyme communication must arise from physical constraints that act across the metabolon (**Fig. 1c-d**), an observation supported by the stoichiometry and preferential placement of peripheral subunits in the mammalian PDHc as well^22,37^. Following the observation that E1 and E3 occupy confined spatial zones surrounding the core (**Fig. 1b**), the lateral organisation of peripheral subunits within the redox shell was quantified (**Fig. 2a**). Statistical analysis of peripheral subunit positions revealed an average nearest-neighbour distance (NND) of 87 Å (**Fig. 2a, Extended Data Fig. 17**). The spacing distribution closely matches the expected hydrodynamic radii of the free E1 (*R*_t_ = 48 Å) and E3 (*R*_t_= 42 Å) models, explaining a dense but non-overlapping packing within this outer shell at the observed stoichiometries (**Extended Data Fig. 16b** and **Extended Data Fig. 17d**). Although E1 and E3 do not form stable catalytic interactions, their proximity is sufficient to support transient encounters, rationalizing previously published cross-linking data^38^. Random placement under geometric confinement of the redox shell (**Fig. 1d**, Methods) (2.7×10^3^ nm^3^) failed to reproduce the observed spacing with statistical significance, demonstrating that peripheral enzyme positioning is non-random and instead reflects intrinsic constraints imposed by tethering and steric packing. The functional consequence of this organization is that most catalytic sites are within linker-accessible distance of the tethered lipoyl domains for efficient substrate cycling within a densely packed yet dynamic redox shell (**Fig. 1e, Extended Data Fig. 18**).

**Fig. 2:**
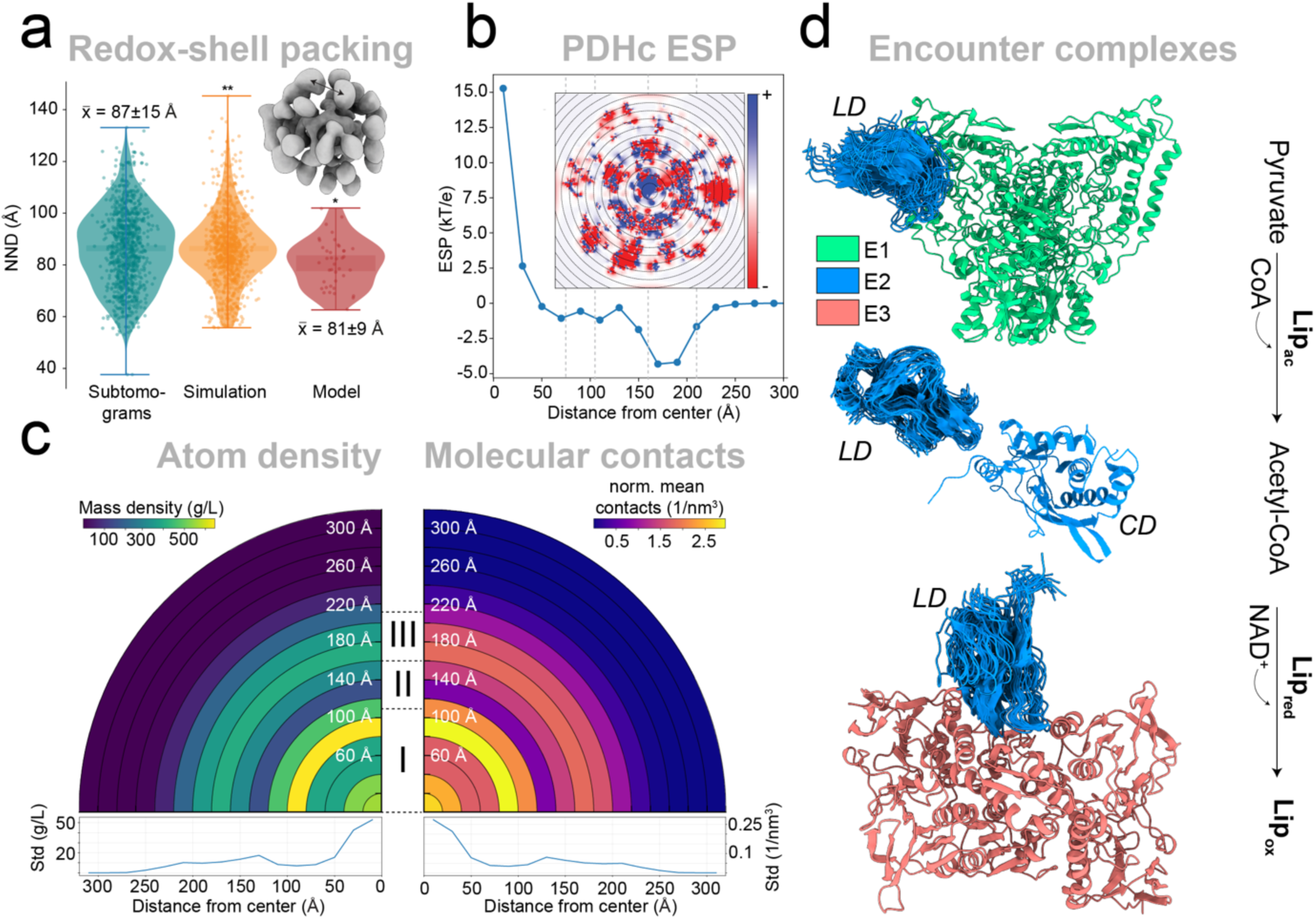
Hierarchical logic of the PDHc shell complex. **a,** Statistical analysis of peripheral subunit spacing revealed an average nearest-neighbor distance (NND) of 87 Å. Poisson disk sampling of COM NND within the redox shell, matched to the experimental mean, yielded a distribution different from the experimental results (KS test, p ≪ 0.01), indicating non-random organization of E1 and E3, also supported by the integrative model**. b,** Electrostatic potential (ESP) calculated in concentric hollow spheres (inset), revealed distinct electrostatic zones that coincide with the geometrical boundaries of the complex, delineating compartments within the metabolon. **c,** Global radial analysis of the simulated PDHc reveals higher-order organization. The average mass density was calculated in concentric shells and highlights distinct, compartment-specific crowding environments across the complex (left). Normalized molecular contacts (Cα–Cα <10 Å) in radial shells to identify local interaction hotspots and changes in local packing across the PDHc architecture. **d,** Encounter complexes for LD insertion into the active sites of E1, E2, and E3 were extracted from the simulated ensemble. For each target complex, the 20 top-scoring insertion models, ranked by active-site residue distance, are shown.

Analysis of electrostatic potential and mass density as a function of radial distance from the complex center, tracking global trends across the entire PDHc, revealed pronounced compartmentalization that coincides with geometrical boundaries across the assembly (**Fig. 2b, c**). The core-localized E3BP assembly generates a dense, positively charged electrostatic environment, primarily arising from basic residues within intrinsically disordered regions (residues 360-400), which engage complementarily charged surface patches on the E2 catalytic domains to form a rigid scaffold. The IDR-rich nested shell between the core and the peripheral region exhibits an overall neutral electrostatic character, consistent with its permissive and flexible nature, whereas the E1/E3-occupied redox shell is characterized by a net negative electrostatic environment, which is likely to reduce the residence time of the negatively charged LD (**Fig. 2b**).

To probe the dynamic organization of the full PDHc beyond the static model, we performed guided all-atom metadynamics electron microscopy metainference (MEMMI) simulations of the complete assembly using the asymmetric reconstruction (**Extended Data Fig. 13, Extended Movie 1**) as simulation restraints (Methods). We first examined global properties by quantifying radial mass density (**Fig. 2c**) and normalized molecular contacts (**Fig. 2c**, **Fig. 2d**) from the center of the complex outward, revealing a physically stratified architecture with distinct core, intermediate, and peripheral regions. The same ensemble further enabled sampling of linker dynamics and transient encounter states that are inaccessible in static SPA or CET models (**Extended Data Figs. 19-26**). To validate the molecular interactions underlying this experimentally-guided molecular simulation, we mapped uncharacterized cross-links from mass spectrometry (XL-MS) data^38^ and provided orthogonal experimental restraints reporting on proximity relationships across the native, dynamic complex (**Extended Data Fig. 27-35**).

Analysis of the integrative ensemble shows close agreement with the experimentally observed crosslinks (**Extended Data Fig. 28**), with 93% of intra- and 80% of inter-subunit restraints satisfied (28 Å cut-off). Inspection of the underlying interaction patterns reveals that the flexible E2 and E3BP chains form frequent contacts between flexible segments flanking the PSBD and LD (**Extended Data Fig. 19-20, 29-31**), demonstrating that local disorder acts as an active architectural element that aids PSBD coordination and complex stability within the crowded nested shell. Furthermore, E2/E3BP PSBDs were found to interact frequently with elements beyond the known E1/E3 interaction site (**Extended Data Fig. 19-20**), with enrichment among PSBDs from different E2 trimers, suggesting a mechanism reminiscent of PSBD dimerization observed *in vitro* for the heterologously expressed bacterial counterpart^24^ (**Extended Data Fig. 23**). A second frequent interaction site was the flexible linker that resides within the core binding domain CBD, previously highlighted for a highly conserved stretch of amino acids of unknown function^13^ (residues 377-382, **Extended Data Fig. 36**). Our model shows that alternate charges within this element underlies frequent interactions with the trimeric interface of the E3BP core domain (**Extended Data Fig. 19** and **Extended Data Fig. 30**), as well as with the E2 core domain CD (**Extended Data Fig. 20** and **Extended Data Fig. 31**). These interactions indicate that the conserved core loop stabilizes the E3BP trimer, securing it firmly within the central assembly. Consequently, the adjacent flexible regions appear to emanate from a shared, stably anchored origin.

The MEMMI ensemble and crosslinks further capture catalytically relevant encounter states for all steps of the reaction cycle. LDs repeatedly sample geometries proximal to the E1 active site (**Fig. 2d**, **Extended Data Fig. 21c, 22, 32**), forming transient encounter complexes which structurally rationalize the suggested pre-insertion states required for pyruvate decarboxylation^37^ (**Extended Data Fig. 37**). Similar, though less frequent, insertion-compatible configurations were observed for E2 LD-CD interactions (**Fig. 2d, Extended Data Fig. 23** and **Extended Data Fig. 29**), agreeing with the transient nature of transacetylation^5^. For LD reoxidation, E2 LDs engage E3 through a defined α-helical docking interface (**Fig. 2d, Extended Data Fig. 24** and **Extended Data Fig. 33b**), whereas E3BP LDs display markedly fewer productive contacts (**Extended Data Fig. 22, 25c, 34**), reflecting related geometric constraints imposed by E3BP organization. In agreement with cryogenic electron tomography (CET) and subtomogram analyses, direct E1-E3 interactions were rare (**Extended Data Fig. 26** and **Extended Data Fig. 35**), consistent with electrostatic repulsion within the outer shell structure of PDHc that enforces their spatial segregation and functional coupling exclusively through E2/E3BP-mediated lipoyl transfer.

Together, these results define a biophysical framework in which PDHc organization and catalysis emerge from the interplay of spatial confinement, electrostatics, and extensive conformational variation driven by flexible and unstructured, spatially confined regions. Importantly, the MEMMI ensemble reveals that catalytically relevant encounter states are already partially populated in the ground-state architecture, and are expected to be selectively stabilized or disfavoured depending on the chemical state of the lipoyl group, consistent with reported differences in subunit affinities for oxidized, reduced, and acetylated lipoyl domains^2^. Long-range, directional electrostatic gradients and PDHc-embedded, spatially distinct macromolecular crowding conditions further bias lipoyl domain trafficking, disfavoring residence within the negatively charged redox shell while shuttling to the electrostatically stabilized core.

### Substrate-induced remodeling of the conformational landscape

After delineating the principles and architectural features of the ground state, we analyzed how substrate binding affects the conformational landscape of the PDHc. Previous structural analyses suggested a disorder-to-order transition within the E2 active site in the fungal PDHc, which may selectively regulate LD interaction^5,6^, coupling catalysis with substrate-induced shifts in conformational equilibria.

To address this, we resolved the endogenous PDHc core under substrate- and cofactor-saturated conditions, reaching a resolution of 3.1 Å (FSC = 0.143) (**Extended Data Fig. 38** and **Extended Data Fig. 39**). Direct comparison with the apo state revealed several substrate-dependent changes: Firstly, a measurable increase in particle diameter was observed (**Extended Data Fig. 2b**), indicating a subtle expansion of the assembly under catalytic conditions. Secondly, the binding mode of CoA was resolved within the active site, expanding on previous results^5,39,40^, and representing the first experimentally determined structure of a CoA-bound eukaryotic PDHc (**Fig. 3a, Extended Data Tab. 1&2**). Here, the adenosine and phosphate moieties are anchored via a network of basic residues, including Lys281, Arg269, and Arg357, positioning the pantetheine arm towards the LD-binding tunnel and pre-organizing the thiol for nucleophilic attack. Lastly, substrate binding induced an observable shift in the folding equilibrium of the N-terminal element that forms the entrance to the LD insertion tunnel (**Fig. 3b-c**). In the ground state, this segment is resolved in approximately 52% of active sites, reflecting a dynamic equilibrium between ordered and disordered conformations. Upon substrate saturation, the equilibrium shifts towards a fully ordered α-helical configuration at all 60 observable sites. This folding transition generates an extended, positively charged interface that stabilizes docking of the negatively charged LD and reinforces the catalytic tunnel architecture^5^. Notably, this mechanism differs from the gating-loop displacement described for the homologous branched-chain keto acid dehydrogenase complex^39^ (**Extended Data Fig. 40**). Rather than relying primarily on local loop movements to transiently open the LD insertion pore, the fungal PDHc appears to exploit a cooperative, domain-level stabilization mechanism, where ground-state conformational ensembles that sample catalytic state configurations go beyond those reported for isomerases^41^.

**Fig. 3:**
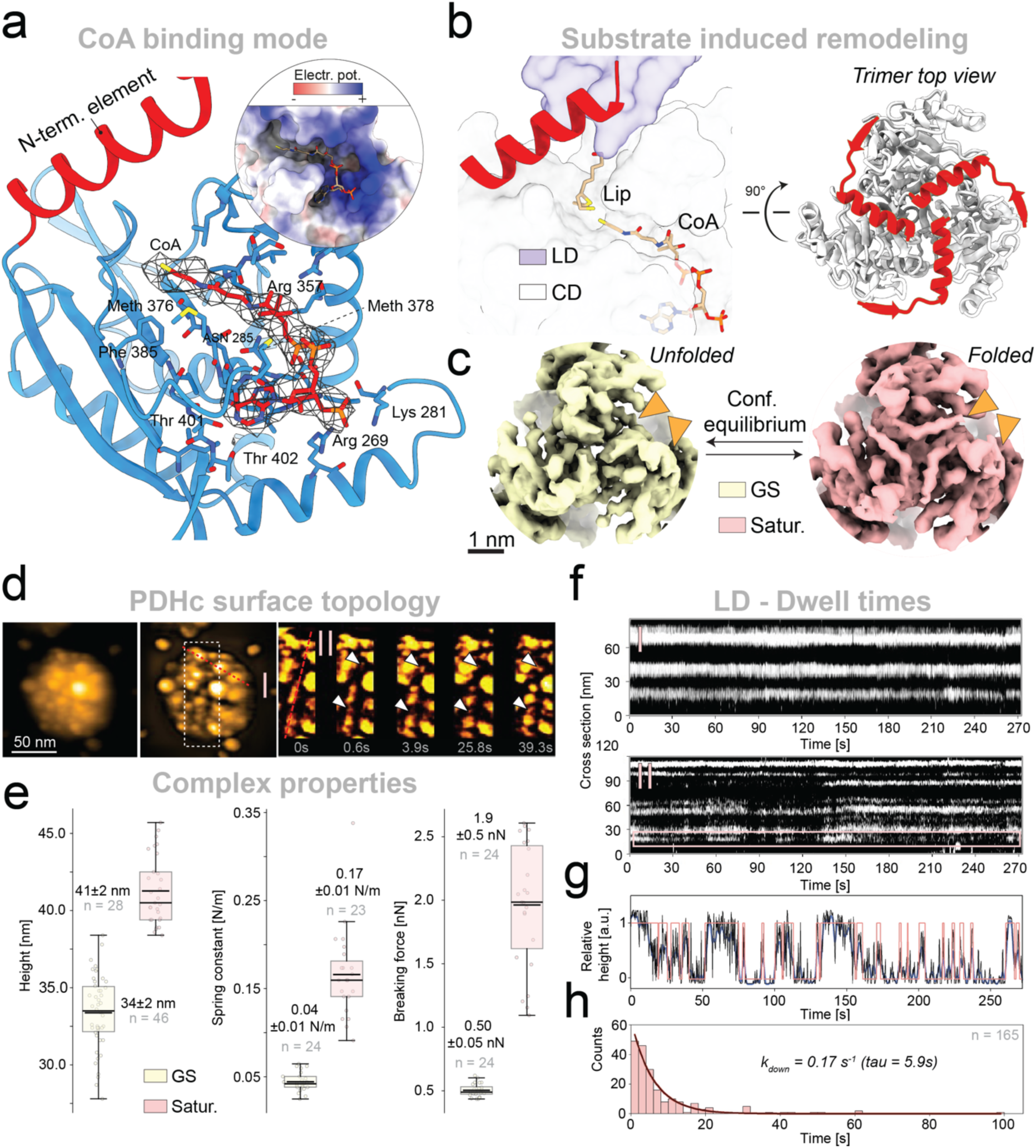
Conformational plasticity drives catalytic coordination. **a,** CoA binding within the active site is stabilized by charge complementarity between its negatively charged phosphate groups and basic residues of the E2 catalytic domain. Interacting side chains and density are indicated. **b,** The N-terminal segment (red), guides LD insertion, taking part in the formation of the transacetylation tunnel. **c,** A substrate-dependent equilibrium shift transitions a partially folded active site to a fully ordered one under substrate saturation conditions. **d,** HS-AFM image of a PDHc molecule adsorbed on a mica surface (left), bandpass filtered image revealing sub-domain organization (middle), and (right) time-lapse zoomed in image from the white dotted rectangle box showing the height fluctuations of eight LD occupancy over time with translocation events highlighted (white arrows). Dashed red lines indicate line-scan trajectories used to extract temporal height profiles. **e,** Substrate saturation increases the observed particle height, consistent with the effects observed in SPA (left). To probe the mechanical properties of endogenous PDHc, nanoindentation experiments were performed, revealing an increased spring constant (middle) and breaking force (right), consistent with increased mechanical stability. **f,** Kymographs showing the height fluctuations along each scan path (**I**, **II** in panel a) reveal localised domain rearrangements (arrowheads in a). (**I**), While the larger domains (top) show rare or no significant rearrangements, the smaller subunits exhibited more frequent cycling behaviour (bottom, kymograph **II**). **g,** Representative relative height profiles over time of one smaller domain marked by a magenta rectangle in f. Black line representing direct cross-section from the kymograph; blue line shows smooth cross-section over two frames; the red line represents the assigned up (1) and down (0) states of the domain of interest. **h,** The axial translocation of LD-signatures was quantified across molecules, yielding a dwell-time reaction rate constant (*kdown*) of 0.17 s^-1^, considering that the down states of LDs can be assigned to the reaction step at the core.

To assess how these local structural changes propagate to the mesoscale architecture during catalysis, we visualized intact PDHc particles by high-speed atomic force microscopy (HS-AFM) (**Fig. 3d, Extended Data Fig. 41-44, Extended Movie 2**)^42,43^. Under saturation conditions, individual complexes displayed a stable globular architecture with consistent dimensions and stoichiometric ratios in line with SPA and CET reconstructions, therefore clearly differentiating between core and peripheral regions (**Fig. 3d, Extended Data Fig. 41-42**). In the presence of all substrates, larger surface signatures corresponding to E1 and E3 assemblies remained largely immobile over the observation period, indicating that the redox shell forms a mechanically stable scaffold, in line with previous observations of preferential angular coverage^22,37^ and our systematic localization of external subunits in the multiple cryo-EM reconstructions (Methods, **Extended Data Fig. 3**). The addition of PDHc substrates caused observable changes in the size and mechanical behaviour of the intact complex (**Fig. 3e, Extended Data Fig. 43a-b**). Particles increased in apparent height from 34 ± 2 nm in the ground state (GS) to 41 ± 2 nm under substrate-saturating conditions, mirroring the substrate-dependent size increase observed by cryo-EM (**Extended Data Fig. 2b**). This size change was accompanied by increased mechanical stability as observed using particle-fatigue assay by HS-AFM imaging (**Extended Movie 3**). Saturated complexes remained structurally persistent over substantially longer recording times (**Extended Data Fig. 43a**) and mechanical properties could be quantified via nanoindentation experiments (**Fig. 3e, Extended Data Fig. 43b**). Incubation increased both the particle spring constant and the breaking force by 4-fold, indicating that catalysis drives PDHc into a mechanically reinforced state comparable to large rigid viral assemblies^10^. The individual force-distance curves further recapitulated the hierarchical organization of the assembly, with compliant redox/nested-shell regions behaving as deformable outer layers and the central core remaining mechanically more resistant.

Localized translocation events became observable, appearing in kymographs as discrete, multi-second transitions between height states. Movements of the larger assemblies in the metabolons’ periphery were a rare event, whereas the smaller domains cycled on timescales of several seconds (**Fig. 3f, Extended Data Fig. 44a**). Quantification of LD-related transitions yielded an apparent rate constant of 0.17 ± 0.01 s^-1^, with a dwell time of ∼6 s (**Fig. 3g&h, Extended Data Fig. 44b, Extended Movie 4&5**). Comparing these observations with catalytic rates observed for the intact PDHc (28.9 - 69.9 s^-1^)^44,45^ places the HS-AFM dynamics into a functional context. Given overall reaction rates on the order of tens of turnovers per second and the presence of 72 lipoyl domains per complex, the effective catalytic demand on an individual LD corresponds to one productive cycle every ∼1–7 seconds. The HS-AFM-resolved transitions, therefore, fall within the expected temporal window for catalysis-coupled relocalization of single lipoyl domains.

These observations extend the biophysical model established for the ground state PDHc (**Fig. 1**, **Fig. 2**), connecting function to structural organization. The compartmentalization of PDHc catalytic steps, electrostatic gradients, and crowding effects related to those define a pre-organized landscape of partially populated encounter states, which is selectively stabilized upon substrate addition. At the atomic level, CoA-induced ordering of the N-terminal active-site element enhances LD docking, which has been shown to significantly decrease K_d_ in dehydrogenase complexes of the same family^39^. Similar observations have been reported for the human PDHc, leading to increased frequency of LD-CD interactions upon CoA addition^8^. At the mesoscale, this shift manifests as substrate-dependent, localized LD translocation events (**Fig. 3d**), while the global architecture of the complex is reorganized towards a mechanically more stable and extended configuration, likely to allow pathways for LD translocation.

### Cellular organization of the PDHc

Recent studies focusing on the cellular organization of the PDHc and related complexes have analyzed isolated mammalian mitochondria, either by collecting tilt series directly from the organelle or by thinning it with an ion beam^25,46,47^. However, mitochondrial isolation disrupts ultrastructural integrity, membrane tension, ionic gradients, and macromolecular crowding, raising the question of how PDHc is organised within its native cellular environment^48^. Such perturbations are well known to induce partial swelling and substantial alterations in cristae morphology and overall mitochondrial architecture, depending on the specific purification procedure ^49^. These methodological issues challenge reported results, requiring a true *in situ* approach to establish the native mitochondrial organization of PDHc and to evaluate how earlier *in vitro* or *in organello* analyses may reflect these.

To overcome these limitations, resolving the PDHc directly in its native environment, *T. themophila* was directly cultivated on TEM grids, vitrified, and processed by cryo-focused ion beam (cryo-FIB) milling to generate electron-transparent lamellae (**Extended Data Fig. 45a-f**). The reconstructed, denoised tomograms showed mitochondria with intact ultrastructure: tightly packed matrices, abundant respiratory chain assemblies, and membrane-bound signatures that closely match the expected morphology of ATP synthase complexes (**Extended Data Fig. 45g**)^50^. Within these exceptionally crowded mitochondrial volumes, individual PDHc particles were visually identified based on their characteristic size and morphology (**Fig. 4a**). Subtomogram averaging yielded a reference-free reconstruction of the rigid E2 catalytic core, resolving its dodecahedral architecture (**Extended Data Fig. 46**). The *in situ* core dimensions closely matched those obtained from the endogenous, biochemically extracted particles, demonstrating that the structural integrity of the E2-E3BP scaffold is preserved within the native mitochondrial matrix (**Fig. 4b**). Statistical analysis of the manually curated PDHc assemblies highlighted that the particle diameters (**Fig. 4c**) closely match the *in vitro* analysis (**Fig. 1a-b, Extended Data Fig. 2b, 47**), confirming that the dimensions *in situ* are similar, and validating the structural integrity of the integrative model. In addition, we found PDHc molecules to be localized in close proximity to cristae and frequently adjacent to regions enriched in respiratory complexes (**Fig. 4c**). This preferential positioning supports a model in which PDHc occupies mitochondrial spaces that facilitate metabolic coupling between pyruvate oxidation and oxidative phosphorylation, previously shown in isolated mitochondria^25^. In addition to membrane-associated populations, a subset of PDHc particles appeared in closely spaced clusters within the matrix (**Fig. 4c**). The observed distances in these regions were lower than expected for random placement and more in line with higher-order metabolic organization, potentially enhancing substrate flux through coordinated pyruvate conversion and NADH production in regions of elevated respiratory demand. Our simulation model resolves transiently protruding LDs beyond the outer oxidation shell surface (**Fig. 2c**), as well as HS-AFM data confirms their surface motions upon substrate addition (**Fig. 3d-h**), therefore suggesting that lipoyl cross-shuttling between two distinct, closely packed metabolons is reasonable. These observations demonstrate that the integrative model of PDHc is applicable within the mitochondrial matrix, extending it to the physiological context of the living cell. The preservation of particle dimensions, core architecture, and steric compatibility within the crowded matrix confirms that the higher-order organization defined from endogenous cell extracts is maintained under native conditions. Moreover, the preferential localization of PDHc indicates that its spatial distribution is functionally tuned to the metabolic landscape of the mitochondrion.

**Fig. 4:**
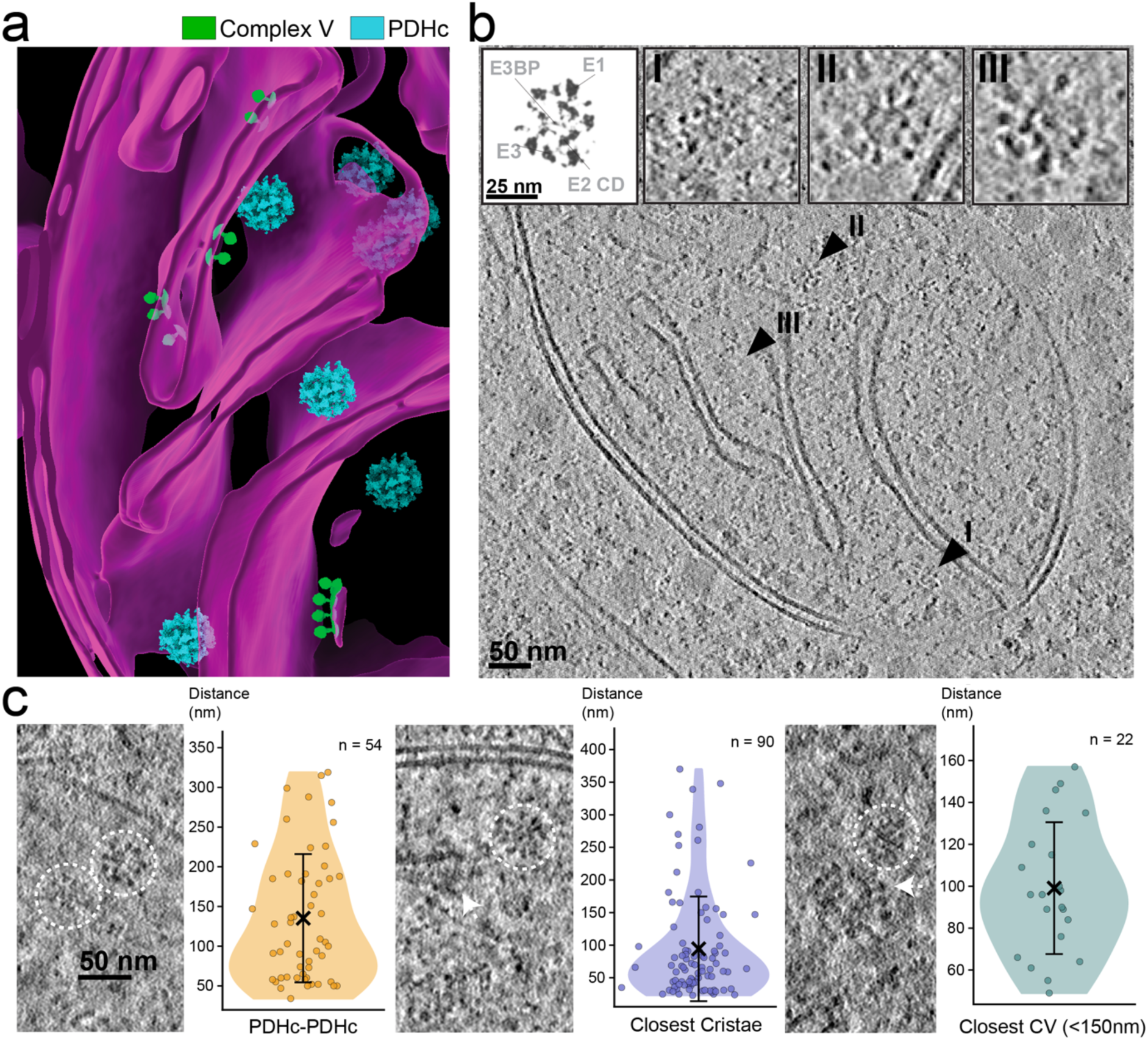
Mitochondrial accommodation of PDHc. **a,** Segmented mitochondrial membranes are shown in magenta, with PDHc particles in cyan and Complex V (CV) in green. **b,** Representative cryo-electron tomogram slice of mitochondrial cristae and spatially associated PDHc (**I-III**). The simulated slice of the integrative model closely matches the experimentally determined signatures. Positions of the PDHc molecules (**I-III**) in the original tomogram are marked. **c,** Spatial analysis of PDHc positions demonstrates higher-order PDHc-PDHc association (left), preferential localization near mitochondrial cristae (middle), and colocalization with Complex V (CV) of the respiration chain (right).

### The nested shell, an architectural model for pyruvate oxidative decarboxylation

The delineated principles of PDHc function are based on the compartmentalization of specific reaction steps within a spatially organized multienzyme metabolon. Rather than operating as a freely diffusing collection of enzymes, the PDHc establishes a hierarchical reaction environment in which geometry (**Fig. 1, 2a**), electrostatics/molecular crowding (**Fig. 2b**), and tethered flexibility (**Fig. 2 c**) collectively govern substrate flow. Our integrative analyses reveal that pyruvate oxidative decarboxylation occurs within concentric functional shells that coordinate catalysis across molecular, supramolecular, and cellular scales (**Fig. 5a**). Based on the presented results, PDHc can be structured into three concentric, distinct regions:

**Fig. 5.**
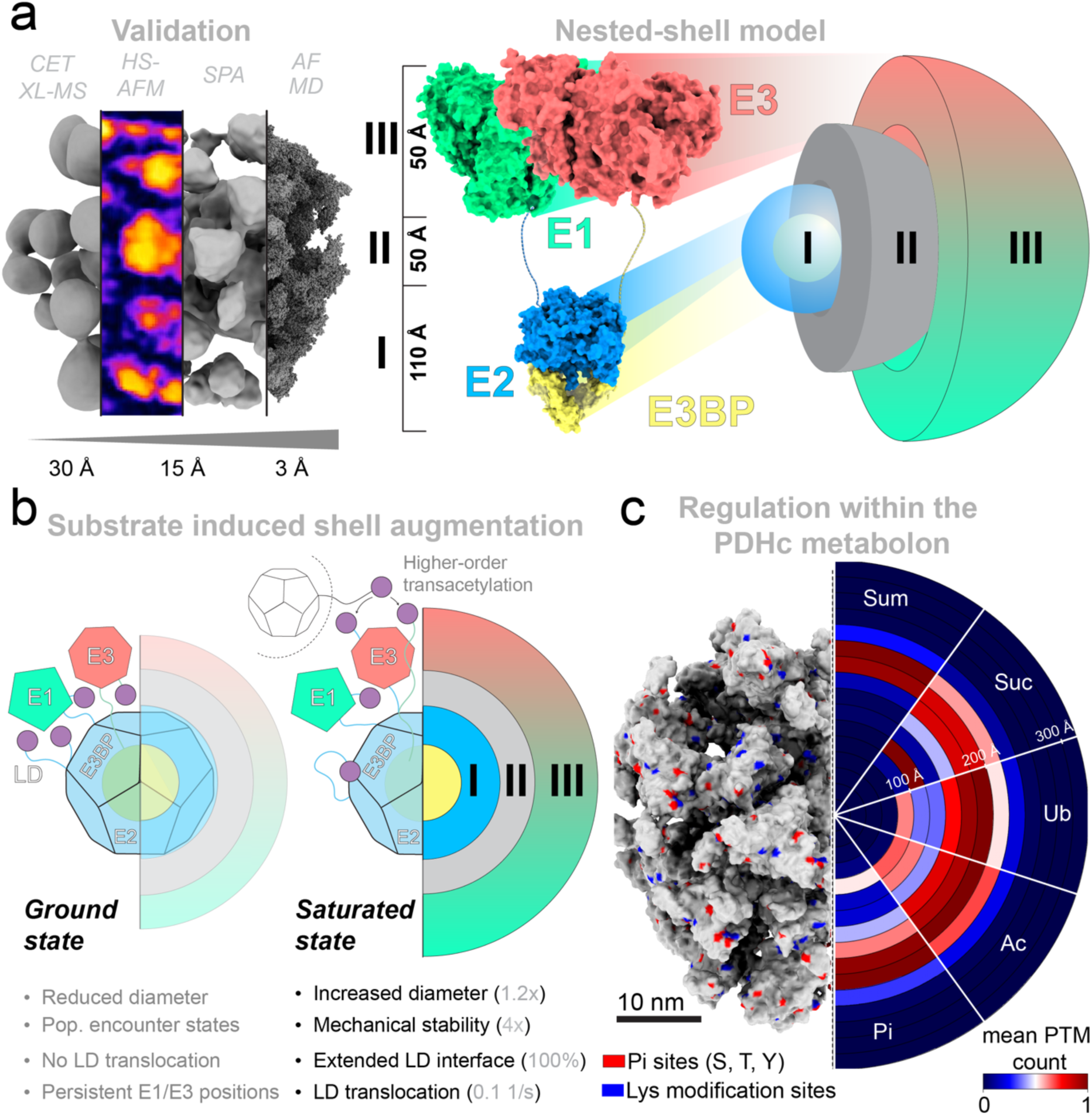
The nested shell model. **a,** The nested shell model links the overall PDHc morphology to biophysical principles. The E2 CD (blue) and E3BP (yellow) form the inner scaffold. Flexible linker regions of E2 and E3BP, containing the lipoyl (LD) and peripheral subunit-binding (PSBD) domains, give rise to a heterogeneous exclusion zone that mechanically and spatially couples the core scaffold to the outer redox shell formed by the E1 and E3 complexes (green and orange)**. b,** The PDHc was resolved in two conformational states, a ground state and a substrate-saturated state. In both, the inner scaffold formed by the E2 CD and E3BP CBD remains largely structurally conserved. Substrate saturation induces expansion and mechanical stiffening of the complex and is accompanied by observable LD translocation. Together, these findings support a model in which the giant PDHc assembly adapts its structural properties to substrate availability, thereby modulating LD trafficking across the metabolon. **c,** The integrative model of the PDHc metabolon expands not only on the structure-function relationship of pyruvate oxidative decarboxylation but can be expanded to provide structural explanations for mitochondrial regulation via PTMs within the PDHc. Putative regulatory PTM sites were mapped onto each frame of the simulated PDHc ensemble and radially averaged to estimate their potential distribution across the structural shells of the metabolon. Lysine modification sites (blue) and phosphorylation positions (red) are displayed on the structural model (left). Phosphorylation (red) sites and lysine modifications (lysine acetylation, ubiquitination, succinylation, and sumoylation), mapped from yeast and human orthologs, are marked in the integrated model (left) and annotated for each protein sequence (right).

I) At the center of the 7.6 MDa assembly, the E2 CD and E3BP CBD form a rigid scaffold to direct the anchoring of the peripheral domains and a distinct region that gates CoA production. This inner hub, built from a dodecahedrally enclosed tetrahedron, extends beyond a purely structural role and defines a chemically specialized transacetylation compartment. While the role of the E2 in this reaction has been characterized at atomic detail previously^5,9^, we extend the functional interpretation of the entire PDHc core, resolving both its unbound and CoA-bound states. Our results suggest that E3BP comprises an active architectural element that reinforces and stabilizes the core (**Fig. 2b**). Beyond its canonical binding interface (**Extended Data Fig. 10**), E3BP engages in transient electrostatic interactions mediated by highly conserved, intrinsically disordered regions that contact neighboring E2 catalytic domains and adjacent E3BP trimers (**Fig. 2c, Extended Data Fig. 19, 20**). These interactions reinforce the scaffold and contribute to core rigidity through a self-stabilizing network of flexible contacts.

While models such as the random sampling of equivalent LDs^51^, and partial reaction zones with distinct catalytic functions^6,8^ have been proposed, it remained unclear how a system-wide communication could be carried out. We show that CoA binding directly couples core chemistry to mesoscale organization by shifting the folding equilibrium of the CoA-binding tunnel. Under substrate-saturated conditions, CoA binding stabilizes the N-terminal element of the E2 catalytic domain, generating an ordered and positively charged LD docking interface (**Fig. 3a-c**). This transition not only enhances transacetylation efficiency but also shifts the conformational ensemble of the tethered linker regions, effectively shortening the functional reach of E2 linkers and biasing lipoyl-domain trajectories towards the catalytic core. In this way, substrate-induced ordering at the active site propagates outward, expanding and rigidifying the entire PDHc, linking chemical state to LD positioning and promoting productive encounter states within the nested-shell architecture.

II) The core is enveloped by an intermediate “nested” shell formed by flexible regions, PSBDs, and lipoyl domains of LDs of E2 and E3BP. The nested shell exhibits a generally neutral electrostatic potential, compared to the positively charged core and negatively charged redox shell (**Fig. 2b**). This molecule-wide gradient creates a macromolecular polarity that allows LDs to sample a wide conformational space while preventing non-productive trapping in either neighboring compartment. Within this zone, frequent contacts among the disordered segments flanking the PSBDs and LDs are observed, demonstrating an active architectural element that stabilizes peripheral subunit coordination and maintains shell integrity despite those regions participating in interactions of low complexity (**Fig. 2c, Extended Data Fig. 19, 20, 29-31**). Beyond canonical E1/E3 recruitment, PSBDs form transient contacts across neighboring E2 trimers and E3BP subunits, reminiscent of PSBD dimerization in bacterial PDHc^24^. These interactions promote peripheral subunit clustering (**Extended Data Fig. 11**) and position lipoyl domains near E1 and E3 active sites for the formation of insertion complexes (**Fig. 2d**, **Extended Data Fig. 21, 23, 24**). Functionally, this IDR-rich zone of PDHc acts as a selective molecular sieve analogous to the FG-repeat-containing nucleoporins that form the inner channel of the nuclear pore complex^52^, controlling rapid carrier sampling while biasing productive trajectories and excluding non-functional states. Recent work also shows that IDR-based condensates can mediate nonenzymatic C-N bond formation, including pyruvate-derived reactions, suggesting that the PDHc IDR-rich shell could influence downstream chemistry beyond substrate simple subtrate channeling^53^.

III) The outermost region constitutes the redox shell, formed by spatially confined E1 and E3 that enclose the nested shell. E1 and E3 occupy segregated but LD-accessible positions, creating a discrete decarboxylation-reoxidation zone that is functionally coupled exclusively through lipoyl-mediated transfer, akin to a combined mechanism of the “Division of labor”^8^ and the “Pyruvate dehydrogenase factory organization”^6^ modes. The dense yet non-overlapping packing suppresses non-productive E1-E3 contacts through electrostatic segregation (**Fig. 2a-b**) and maximizes productive encounter complexes, even under ground-state conditions (**Fig. 2c-d**), enforcing catalytic competence, despite the absence of direct physical connection. HS-AFM shows that this peripheral layer is mechanically stable at the global level while supporting local, catalytic-timescale rearrangements, indicating that redox cycling occurs within a structurally persistent framework (**Fig. 3d-h**). These findings establish a unified structural framework that explains how the PDHc mesoscale architecture encodes catalytic function (**Fig. 5a**). The nested-shell model reconciles decades of biochemical and kinetic observations by showing how spatial confinement, electrostatic compartmentalization, and LD dynamics collectively enforce a sequential, ordered intermediate transfer without the need for stable protein-protein interaction (**Fig. 2**). Catalysis, therefore, emerges from a mesoscale reaction environment that pre-organizes encounter states and biases lipoyl-domain trafficking along productive pathways. Importantly, the same organizational principles are preserved *in situ*, showing agreement between cell-extract-derived and *in situ* PDHc in dimensions, core geometry, and peripheral zoning (**Fig. 4**).

The availability of an experimentally restrained, all-atom mesoscale model further allows quantitative analysis of higher-order behaviors such as localization of PTMs within the entire shell structure of PDHc (**Fig. 5b, Extended Data Fig. 48, Extended Data Tab. 4-8**). Relevant encounter geometries are already populated in the ground state (**Fig. 2d, Extended Data Fig. 21-25**), indicating that the complex continuously samples transient interactions without large-scale architectural rearrangement. In this context, access to modifying enzymes, including PDKs/PDPs and factors responsible for lysine-directed modifications, emerges from LD cycling: LDs revisit defined docking interfaces and traverse the nested shell on catalytic timescales, rendering modification events feasible. These modifications are expected to act at the level of the entire metabolon by perturbing an already crowded and electrostatically structured reaction environment. Consistent with this view, radial PTM analysis resolves a non-uniform regulatory landscape across the nested shells: phosphorylation sites are enriched toward the peripheral reaction layers, whereas lysine-based modifications are distributed more broadly across the metabolon and extend into core-proximal linker regions. Because mitochondrial acyl-CoA intermediates, including acetyl-CoA and succinyl-CoA, can nonenzymatically modify lysine residues ^54^, lysine acylation is expected to accumulate broadly across PDHc, including core-proximal regions, whereas enzyme-directed PTMs remain more strongly confined to the peripheral shells. This shell-dependent partitioning supports the notion that PTMs regulate PDHc not by uniformly tuning isolated subunits, but by selectively remodeling electrostatic and steric properties along the preferred LD trafficking routes. These PTMs introduce local charge and steric changes within regions that govern encounter probabilities, IDR corridors, LD docking surfaces, and peripheral enzyme interfaces, restructuring and biasing the lipoyl-domain trajectories.

## Methods

### Cultivation, harvest, and cell lysis of *T. thermophila*

Cultivation and extract preparation of *Thermochaetoides thermophila* were performed as previously described^6,55^, with minor adaptations. Cells were grown at 52 °C under 10 % CO₂ in complete culture medium (CCM) containing 3 g/l sucrose, 0.5 g/l NaCl, 0.65 g/l K_2_HPO_4_ x 3 H_2_0, 0.5 g/l MgSO_4_ × 7 H_2_0, 0.01 g/l Fe(III)sulfate-hydrate, 5 g/l tryptone, 1 g/l peptone, 1 g/l yeast cell extract, 15 g/l dextrin, and 20 g/l agar for solid culture media. Mycelium, grown for 3 days on solid CCM plates, was used to inoculate 200 ml of CCM. The precultures were then incubated for 20 h at 110 rpm (52 °C, 10 %CO_2_). The main cultures were inoculated with 25 ml of preculture, and mycelial globules were transferred into 800 ml of CCM media. Main cultures were grown for 16 h, until the mycelium formed even globules of 5 to 10 mm. Mycelium was harvested using a 180 µm sieve, washed three times with ice-cold PBS (pH 7.4), and centrifuged at 3,000 *g* for 5 min at 4 °C. Pellets were flash-frozen in liquid nitrogen, cryo-ground, and stored at −80 °C. For lysis, frozen mycelium was homogenized with zirconia beads in lysis buffer (100 mM HEPES pH 7.4, 95 mM NaCl, 5 mM KCl, 1 mM MgCl_2_, 0.5 mM EDTA, 5 % glycerol, 1 mM DTT, 10 μg/ml DNAse, 2.5 mM Pefabloc, 40 μM E-64, 130 μM Bestatin, 0.5 μM Aprotinin, 1 μM Leupeptin, and 60 μM Pepstatin A). Samples were disrupted using a FastPrep homogenizer (3 × 20 s, 6.5 mps, 4 °C), cooling the sample on ice between cycles for 3 min. Cell debris was removed by centrifugation (4,000 *g*, 5 min, 4 °C), and the soluble fraction was clarified by ultracentrifugation (100,000 *g*, 45 min, 4 °C). The lipid layer was removed, and the extract was filtered (0.22 µm) prior to size-exclusion chromatography.

### Enrichment of the endogenous PDHc

The clarified cell lysate was concentrated to ∼30 g/l (measured via Bradford assay) using pre-equilibrated Amicon^®^-cellulose spin filtration columns (100 kDa cutoff) at 3000 *g*. Chromatography was performed on an ÄKTA pure 25 M FPLC system. Size exclusion chromatography (SEC) was carried out on a Biosep SEC-S4000 column equilibrated in running buffer (100 mM HEPES pH 7.4, 95 mM NaCl, 5 mM KCl, 1 mM MgCl₂, 5 % glycerol) at 0.15 ml/min (**Extended Data Fig. 1a**). 500 µl Lysate was injected via an external loop, and fractions of 250 µl were collected. High-molecular-weight SEC fractions (fractions 1-10; 30 runs), previously shown to contain intact PDHc^6,55^, were subjected to anion exchange chromatography (AIEX) on a Capto HiRes Q 10/100 column. The equilibration buffer (Buffer A) contained 100 mM HEPES, pH 7.4, 50 mM NaCl, 5 mM KCl, 1 mM MgCl₂, and 5 % glycerol, and the elution buffer (Buffer B) contained 100 mM HEPES, pH 7.4, 50 mM NaCl, 5 mM KCl, 1 M NaCl, 1 mM MgCl₂, and 5 % glycerol. Samples were loaded at 1 ml min⁻¹, washed with 10 % buffer B for 10 column volumes (CV), and eluted using a stepwise linear gradient (10–25 % B over 10 CV, 25–35 % B over 10 CV, and 35–100 % B over 1 CV). PDHc-containing fractions were identified by activity assay, and composition was analyzed via SDS-PAGE, pooled, and concentrated to ∼1 g/l using Amicon^®^-cellulose spin filtration columns (100 kDa cutoff) at 3000 *g*.

### PDHc activity assay

PDHc activity was measured using a WST-8–based coupled assay adapted from^17^. Each 100 µl reaction mixture contained 100 mM NaCl, 30 mM K₂HPO₄, pH 7.5, 2 mM MgCl₂, 2 mM ThDP, 3 mM NAD⁺, 0.4 mM CoA, 4 mM pyruvate, and 4 µl Cell Counting Kit-8. 2 µl of the respective fraction were pre-incubated without pyruvate for 5 min at 37 °C, and reactions were initiated by pyruvate addition. Formazan formation was monitored at 460 nm for 60 min, and initial rates were obtained from the linear steady-state phase.

### Cryo-EM sample preparation

A detailed overview of vitrification parameters for all datasets is provided in **Extended Data Table 9**. Grids were glow-discharged using a PELCO easiGlow™ (0.4 mbar, 15 mA, 25 s). Standard filter paper (Grade 595, Ø55/20 mm) was equilibrated for 10 min prior to vitrification. A 3.5 µl aliquot of the sample was applied to TEM grids and plunge frozen using a Vitrobot^®^ Mark IV (TFS). All samples were frozen at 4 °C and 100% humidity. Vitrified grids were clipped and loaded under cryogenic, low-humidity conditions according to the manufacturer’s instructions.

### Cryo-FIB milling

Vitrified grids containing *T. thermophila* hyphae were milled to create electron-transparent lamellae using an Aquilos 2 cryo-FIB/SEM (TFS) under high-vacuum cryogenic conditions with AutoTEM (**Extended Data Fig. 45**). Grids were transferred in a cryo-shuttle, brought to eucentric height, and regions of interest were identified in the SEM at 5 kV and 12.5 pA (dwell time 3 µs; working distance 7.1 mm). Samples were sputter-coated (15 s, 30 mA, 10 Pa) followed by organometallic Pt deposition using the gas injection system. Stress-relief trenches were milled on both sides of the target area using a 30 kV Ga⁺ beam at 1 nA, leaving a ∼10 µm-wide lamella window. Lamellae were thinned symmetrically at 30 kV using a stepwise current reduction (1.0 nA, 0.5 nA, 0.3 nA) at a stage tilt of 10° with the ion beam normal to the lamella surface. Final polishing was performed at 30 kV with 30 pA followed by 10 pA to a final thickness of 120–200 nm. Lamellae were transferred under liquid nitrogen to the TEM for imaging.

### Cryo-EM screening and data acquisition

Grid screening, data collection, and automated alignment were performed using TFS EPU (v3.7.1) for single-particle analysis (SPA) and Tomo (v5.17.0) for cryo-electron tomography (cryo-ET). Atlas maps were acquired, and grids were evaluated from low to high magnification to assess particle distribution, ice thickness, and contamination. Data were collected under low-dose conditions, with autofocus and tracking performed on adjacent carbon regions. Detailed acquisition parameters are provided in **Extended Data Table 10.**

SPA datasets were collected on a 200 kV Glacios (TFS) equipped with a Falcon 4i detector (Datasets 1–2) and on a 300 kV Titan Krios G3 (TFS) equipped with a Gatan K3 camera and BioQuantum energy filter (Dataset 3). For the PDHc ground state, 45,780 micrographs were acquired on the Glacios at 92,000× magnification (1.53 Å pixel size) with a total dose of 30 e⁻/Å² and a defocus range of −3.0 to −0.8 µm (Dataset 1). An additional ground-state dataset was collected on the Titan Krios at 64,000× magnification (1.36 Å pixel size), yielding 19,049 movies at a total dose of 30 e⁻/Å² and a defocus range of −3.0 to −1.0 µm (Dataset 3). A substrate-saturated PDHc dataset was acquired on the Glacios under the same optical conditions (92,000× magnification, 1.53 Å pixel size, 30 e⁻/Å² dose, −3.0 to −0.8 µm defocus), comprising 35,668 movies (Dataset 2).

Tilt series were acquired both on a 300 kV Titan Krios G3, equipped with a Gatan K3 camera and BioQuantum energy filter (Dataset 4) and a 300 kV Titan Krios G4 with a Falcon 4i camera and a Selectris energy filter (Dataset 5). Tilt series were acquired in a dose-symmetric scheme with a tilt range of -60° to +60° and 3° increments. The PDHc ground-state tomographic dataset (Dataset 4) comprised 181 tilt series collected at 42,000× magnification (2.13 Å pixel size), with a total dose of 144 e⁻/Å² and a defocus range of −5.0 to −1.0 µm. The tilt series of *T. thermophila* lamellae were acquired at 42,000× magnification (2.38 Å pixel size), consisting of 229 tilt series with a total dose of 140 e⁻/Å² and a defocus range of −5.0 to −1.0 µm (Dataset 5).

### Image analysis from single-particle data

Datasets 1–3 were processed in CryoSPARC (version 4.1.1–4.7.1^56^). A divide-and-conquer workflow (**Extended Data Fig. 3**) was applied to reconstruct the E2 CD (**Extended Data Fig. 4-5**), the E3BP CBD (**Extended Data Fig. 6-7**), the asymmetric full complex (**Extended Data Fig. 13**), peripheral subunits (**Extended Data Fig. 14-15**), and substrate-dependent states (**Extended Data Fig. 38-39**). Datasets 1-3 were imported and subjected to patch motion, followed by patch CTF estimation.

Template-based particle picking was initiated using 15 projections derived from the previously reported fungal PDHc core (PDB ID: 7OTT^5^). An initial picking round on a subset of 100 micrographs yielded a high-quality particle set used for template refinement and subsequent picking across the full dataset. Templates and micrographs were low-pass filtered to 10 Å and processed with a particle diameter of 600 Å. After iterative 2D classification, 683000 particles were retained and extracted with a box size of 512 px. Using the mask_create function in RELION 3.0 ^57^, a mock volume approximating the dimensions of the PDHc was generated (**Extended Data Fig. 4**). Heterogeneous refinement of the selected particles and the generated mock volume was performed, resulting in three volumes being reconstructed at 10.6, 8.6, and 8.8 Å (FSC = 0.143), respectively. The highest-resolution map served to generate 50 new templates for picking on Dataset 4. Following further 2D classification, 421,139 particles were refined with icosahedral symmetry to 2.98 Å. Reference-based motion correction was carried out, and a final homogenous refinement resulted in a final resolution of 2.84 Å (FSC = 0.143). To analyze the folding state of the E2 CD LD entry site, previously observed in^5^, 3D classification was carried out (6 Å filter resolution, 10 classes), identifying the subset of particles corresponding to the folded/unfolded state.

To reconstruct the E3BP CBD, the particles initially picked using the E2 CD reference were re-extracted with a box size of 300 px, and a heterogeneous refinement with tetrahedral symmetry was performed, yielding a 3.52 Å (FSC = 0.143) reconstruction of the E3BP core domain. Subsequent asymmetric non-uniform refinement and 3D classification resolved two tetrahedral configurations, comprising 69.8% (state A) and 30.2% (state B) of particles. Independent refinement with enforced tetrahedral symmetry of both states resulted in maps at 3.55 Å and 3.74 Å (FSC = 0.143), respectively. Following super sampling (0.76 Å/px), reference-based motion correction and symmetry-enforced refinement improved the resolutions to 3.28 Å (state A) and 3.52 Å (state B) (FSC = 0.143). To enhance local detail for model building, particles from Dataset 3 were picked using templates derived from asymmetric PDHc reconstructions. After motion correction and tetrahedral refinement, symmetry expansion and focused local refinement on an E3BP–E2 interface produced a 3.11 Å map of the E3BP CBD trimer in complex with neighboring E2 CDs.

Reconstruction of the E1 domain was initiated via an initial round of blob picking on the super-sampled Dataset 1, defining a particle diameter between 80 and 120 Å, a lowpass filter for the templates and micrographs of 5 Å, and a minimal separation diameter of 160 Å. Particle picks were extracted with a box size of 312, and Fourier cropped to a box size of 78 pixels. Three rounds of 2D classification were carried out, using a circular mask diameter of 150 Å, an increased number of 50 online-EM iterations, a class batch size of 400, and 300 classes. Particles resembling E1 projections were chosen, reextracted without binning, and used for ab initio reconstruction. Ab initio reconstruction was carried out with an increased number of initial and final iterations (400, 600 iterations), an increased initial and final minibatch size (300, 1000 images), and running the initial minibatch size for 600 iterations. The resulting volume was used to generate 20 equally spaced templates, which were applied for template picking (particle diameter: 120 Å; low-pass filter: 10 Å). Following three rounds of 2D classification, homogeneous refinement with C2 symmetry produced a final reconstruction at a resolution of 5.8 Å (FSC = 0.143).

To analyze peripheral subunit clustering, asymmetric reconstructions of PDHc were used for template-based picking (particle diameter, 600 Å; low-pass filter, 7 Å; minimal separation distance, 360 Å). A total of 1020732 particles were then randomly split into a subset of 100000 particles and subjected to non-uniform refinements, utilizing a spherical approximation for the peripheral subunits (maximum alignment resolution was limited to 15 Å, disabling auto batch size). The resulting reconstructions were aligned based on their icosahedral core, the core densities (E2/E3BP) were subtracted using ChimeraX, and the peripheral densities were segmented ^58^. COM for the peripheral densities was calculated using the ChimeraX function measure center.

To resolve the substrate- and cofactor-saturated state of the PDHc, template picking was performed using projections from asymmetric reconstructions of the full complex (particle diameter 450 Å; low-pass filter 4 Å; minimum separation 270 Å). Extracted particles (box size 600 px) were cleaned by iterative 2D classification, yielding 130633 particles for refinement. Initial icosahedral homogeneous refinement produced a 3.14 Å map of the E2 core. Following 3D classification, the selected particles were subjected to reference-based motion correction and a final icosahedral refinement, resulting in a 3.09 Å reconstruction of the substrate-saturated E2 core (Supp. Fig. 16). Focused 3D classification (10 classes, 6 Å) was used to quantify substrate-dependent ordering of the LD entry element relative to the ground state.

### Image analysis from acquired tilt series and tomogram reconstruction

Dataset 4, containing the tilt series of enriched PDHc, was preprocessed in Warp (v1.0.9)^59^ with motion correction and CTF estimation, excluding tilts with fits worse than 20 Å. After manual alignment in Etomo^60^, tomograms were reimported into Warp, and full tomograms were reconstructed at a binning factor of 3 (ps = 8.53 Å), a z height of 280 nm, normalizing input images and reconstructing tilt-based half maps for denoising. A manually curated subset of the reconstructed tomograms (n = 5) was then used to train the dewedging and denoising algorithm DeepDeWedge ^35^, with a sub-tomogram size of 96 and a missing wedge angle of 60°. 20% of the sub-tomograms were used for validation. 1000 epochs of training were performed, showing fitting and validation stagnation around 500 epochs. The full set of tomograms was then refined based on the trained parameters (**Extended Data Fig. 12a-b**).

The *in situ* analysis of fungal PDHc from intact hyphae was preprocessed in Warp, performing CTF estimation and motion correction. The motion-corrected tilt series were then aligned using AreTomo3^61^, and subsequently reimported into Warp for per-tilt CTF estimation and tomogram reconstruction. Tomograms were reconstructed with a binning factor of 4 (9.52 Å/pix). Manual picking was performed using Cube on Warp-deconvoluted tomograms ^62^, defining PDHc particle positions. The corresponding subtomograms were reconstructed in Warp at a binning factor of 4 (9.52 Å/pix) and 2 (4.76 Å/pix) and imported and aligned in Relion 4^63^. A reference-free model of the PDHc E2 core was reconstructed with Relions 3D initial model and 3D auto-refine functions. The reconstruction was then used as a reference for subsequent subtomogram refinement and postprocessing at a binning factor of 2. Both the initial model and 3D auto-refine processes were performed in all cases with Icosahedral symmetry.

### Subtomogram analysis of PDHc

A curated subset of denoised subtomograms (n = 30) was extracted for per-particle analysis and aligned to a 30 Å low-pass–filtered E2 CD reference (**Extended Data Fig. 17a**). Peripheral densities were assigned based on spatial position, with vertex-proximal densities classified as E1 and face-proximal densities as E3. To analyze peripheral organization, the core signal was removed using a spherical mask, and the remaining densities were segmented after excluding low-occupancy voxels. COM positions were calculated for each peripheral density, and the redox shell was defined as the hollow sphere enclosing 90% of COM positions (**Extended Data Fig. 17b-c**). Randomized control datasets were generated within this shell using 3D Poisson disk sampling with varying minimum distance constraints (n = 100) to probe maximal packing (**Extended Data Fig. 17d**). Nearest-neighbor distances (NNDs) were calculated for each subtomogram, and matched Poisson controls were generated by adjusting particle number and exclusion distance to reproduce the experimental mean NND on a per subtomogram basis (**Extended Data Fig. 17e**). Experimental and simulated distributions were compared using a Kolmogorov–Smirnov test.

### AI-based structure prediction and model building

The asymmetric SPA reconstruction provided the framework for building an all-atom PDHc model. High-resolution maps of the E2 CD and E3BP CBD were rigid-body fitted into the density, while peripheral assemblies were based on AF-Multimer Prediction of the PSBD-bound E1 and E3 complexes (**Extended Data Fig. 49**)^64^. The predicted models were placed into the peripheral densities based on cross-correlation at spatial position relative to the dodecahedral core (E1 at vertices, E3 at faces), and an assumed stoichiometry of 30:60:12:12 (E1:E2:E3:E3BP). AF-predicted linker regions and lipoyl domains were added to connect peripheral modules to the core, and steric clashes were manually resolved in COOT. The model was supplemented with all relevant cofactors, positioned according to their placement in orthologous reference structures: E1 (PDB ID: 2OZL [ThDP] ^65^), E2 (PDB ID: 6ZZK [CoA] ^66^), and E3 (PDB ID: 1EBD [FAD] ^67^).

### Calculation of the electrostatic surface potential across the PDHc metabolon

The ESP of the integrative all-atom model was calculated using the APBS plugin in PyMOL^68^. The resulting OpenDX maps were parsed to extract grid parameters and ESP values, which were reshaped into a three-dimensional array, assigning each voxel a defined position and potential. Radial distances from the grid center were calculated, and ESP values were averaged within concentric spherical shells of 20 Å thickness to generate a radial electrostatic profile across the PDHc.

### Calculations of molecular energetics

High ambiguity-driven protein-protein docking (HADDOCK) refinements are based on the previously described methodology^69^, which has successfully been applied to probe the energetics of metabolic enzymes^5^. The HADDOCK pipeline incorporates a Berendsen thermostat ^70^ to maintain constant temperature, the Crystallography and NMR Software package as the computational engine^71^, and the OPLS force field^72^ for integration of the equations of motion and calculation of energetic components. The trimeric E3BP and E3BP-E2 interface was analyzed via HADDOCK 2.4^73^. Models from *T. thermophila* were compared with the previously reconstructed structure from *N. crassa*^13^. The interface included one E3BP subunit interacting with two E2 subunits originating from adjacent E2 trimers, thereby representing the physiologically relevant inter-trimer contact.

### XL-MS validation of the integrative PDHc model

For the analysis of the molecular interaction sites within PDHc, the XL-MS dataset previously published for the endogenous *T. thermophila* OGDH complex was reanalyzed^38^, serving as an orthogonal validation step of the integrative model. All crosslink positions used in the structural validation were taken exactly as reported in the primary publication and visualized with xiVIEW^74^. Crosslinks corresponding to PDHc components (E1α, E1β, E2, E3BP, and E3) were filtered for unique pairs and mapped onto the integrative model to assess inter- and intra-subunit contacts. To account for conformational heterogeneity, crosslink distances were additionally evaluated across the final MEMMI simulation ensemble. Crosslink positions were projected onto the structural models in ChimeraX^75^ to visualize domain connectivity and validate the assembly against both the static model and the dynamic ensemble.

### MEMMI simulation of the full PDHc

The all-atom, integrative model (Section 2.2.18), was used for the guided simulations. The structure was protonated, resulting in a total of 1073398 protein atoms. A triclinic simulation box of 48.10 × 49.28 × 48.85 nm was constructed with box faces at least 1.5 nm from the nearest protein atom. The system was then solvated with 3459821 water molecules, and the net charge was neutralized with 1244 Na⁺ ions. The AMBER99SB-ILDN ^76^ force field was used for the protein, and TIP3P ^77^ for water. After energy minimization, a 300 ps position-restrained constant NPT annealing protocol at 1 atm was applied, stepping the temperature from 250 K to 270 K to 290 K in 60 ps increments, followed by 120 ps at 310 K. This was followed by an additional 150 ps NVT equilibration at 310 K with restraints on protein heavy atoms. All remaining molecular dynamics parameters were as described previously ^78–80^.

The all-atom model was rigid-body fitted into the asymmetric reconstruction of the PDHc using ChimeraX ^75^. The experimental map was subsequently segmented to retain only voxels within 6 Å of the model, as density decays and becomes information-poor at increasing distance from the specimen ^80^. The trimmed map was then converted into a Gaussian mixture model (GMM), using 10000 components via gmmconvert ^81^, achieving a correlation of 0.9962 with the trimmed density (**Extended Data Fig. 50**). From the preceding 150 ps NVT equilibration, six equally spaced snapshots were extracted to seed a MEMMI simulation with six replicas (one per snapshot), for a total of 84 ns (14 ns per replica), run with PLUMED 2.6.0-dev ^82^ and GROMACS 2020.5 ^83^. MEMMI was performed in the NVT ensemble at 310 K using the equilibration parameters but without heavy-atom restraints; coordinates were saved every 10 ps. The cryo-EM restraint was updated every two MD steps (4 fs) using neighbor lists to evaluate overlaps between model and data GMMs (neighbor cutoff 0.01; list updated every 100 steps). Collective variables used for biasing were biased with the PBMetad ^84^ scheme with a well-tempered, multiple-walkers protocol ^85^. The hill height was set to 0.3 kJ/mol, with a deposition frequency of 200 steps, a bias-factor of 84 and an adaptive Gaussians diffusion scheme ^86^. The 72 biased collective variables (CVs) were the radii of gyration of the disordered linker regions in E2 (res. 215-250) and E3BP (res. 216-270). Final ensembles were obtained by resampling configurations according to converged Torrie–Valleau unbiasing weights, following established procedures ^84,87,88^. Convergence was assessed from time-dependent free-energy profiles and block-averaged error estimates for each CV (**Extended Data Fig. 51** and **Extended Data Fig. 52**). Global and local correlations between the ensemble-derived map and the GMM representation of the 6 Å-trimmed restraint map were computed with gmmconvert^88^.

Simulation results were processed by loading the topology and trajectory files and converting them into a unified data frame to index every atom and residue across all structures of the final ensemble. This step ensured a one-to-one mapping between PDB residue numbering and trajectory indices, and enabled the reconstruction of chain identities. The resulting data frame contained all atom coordinates for each ensemble structure together with explicit chain and protein assignments (E1α, E1β, E2, E3BP, and E3), providing the basis for subsequent domain annotation and ensemble-level structural analyses.

To derive the principal intra- and inter-chain interactions of all PDHc components, a systematic analysis of all proteins across all chains and MEMMI ensemble structures was performed. For each protein, coordinates from all MEMMI snapshots were filtered, retaining only Cα atoms. All ensemble structures were processed to extract per-frame coordinates, which were grouped by chain identity to enable residue distance calculations. The pairwise Cα–Cα distances were computed, and a true interaction was defined as a distance below 10 Å. Contacts satisfying these conditions were accumulated into a residue-residue contact matrix for each protein combination. To eliminate overrepresentation introduced by symmetry, based on the presence of multiple equivalent chains and from the inherent equivalence of residue pairs (A–B = B–A), interaction counts were normalized for unique residue–residue pairs and copy number prior to normalization for the number of ensemble structures. These pair-chain, per-frame contact probabilities were then visualized as intra-/inter-chain contact heatmaps with the protein-specific domain annotation. Intra- and inter-chain contact propensities were directly visualized on the respective atomic models in ChimeraX ^75^. To quantify radial properties of the simulated PDHc, atomic coordinates from the ensemble were assigned to concentric radial shells of 20 Å thickness relative to the frame-wise geometric center of the complex. Shell-wise mass density was calculated from the summed atomic masses normalized by shell volume and averaged across the ensemble. To assess shell-dependent molecular packing, residue-level Cα-Cα contact analyses were performed on the ensemble, identifying the closest valid partner for each residue, excluding self-matches and immediate sequence neighbors within the same chain. Contacts were scored using a 10 Å cutoff and normalized by shell volume to obtain radial contact densities. Mean values and standard deviations across the ensemble were used for visualization. For binary LD-enzyme interaction analyses, nearest-neighbor Cα-Cα distances were calculated in the ensemble structure between the lipoamide-bearing lysine of the LD and the selected catalytic-site residue. Potential encounter complexes were scored using a 25 Å cutoff (chosen to account for the bridging lipoamide group and to account for flexibility), and the number of possible satisfied contacts per complex was quantified across the ensemble and benchmarked against thresholds of 15-30 Å. Potential PTM sites were mapped onto the corresponding residues in the model and, using Cα coordinates as positional proxies, their radial distributions were quantified across frames and expressed as shell-wise averaged occupancies. These values represent the potential spatial distribution of candidate PTM sites within the assembly.

### HS-AFM of the PDHc

AFM was performed using an SS-NEX High-Speed Atomic Force Microscope (RIBM, Japan) in amplitude modulation tapping mode in liquid. Ultrashort cantilevers (7 μm) with high resonance frequency (1200 kHz) (USC-F1.2-k0.15, Nanoworld, Switzerland) were used. AIEX-enriched PDHs samples in 100 mM HEPES pH 7.4, 50 mM NaCl, 5 mM KCl, 1 mM MgCl_2_, 5% glycerol were incubated on a freshly cleaved mica surface for 10 min. Substrate-dependent PDHc movements were imaged after the addition of 2 mM ThDP, 4 mM Pyr, 3 mM NAD^+^, and 0.4 mM CoA. All images were processed using ImageJ software^89^. The images were primarily drift-corrected and background-levelled. After the initial corrections, a bandpass filter (2-20 pixels) was used for a better identification of the domains^90^. Kymographs were created using time-resolved HS-AFM movies. For the calculation of the up and down states, two-step boundary conditions were applied; (1) Downward movement of >2 nm was considered as the down state. (2) The relative height data acquired from a kymograph were smoothed over two pixels, in order to impose a minimum of two consecutive frames for the domains to spend on a state (up or down) to be eligible for the state (**Extended Data Figure 44)**. For example, if a domain moves up or down by just one frame, it would not be considered a change in state. The distribution of all time frames of the down states (reaction state) was plotted to extract the dwell time and the rate constant by fitting the following: *y* = *ae* − *kt* (where *k (k_down_)* is the obtained rate constant, and 1/*k* is the dwell time (*Ƭ*) of the reaction).

HS-AFM-induced fatigue experiments were performed by continuously imaging the particle with minimal imaging force, using a free amplitude of 3 nm and a setpoint amplitude of ∼2.4 nm. All fatigue studies were performed using a similar force range and pixel size (1.3 nm/pixel).

### AFM Nanoindentation

AFM-based nanoindentation experiments were performed using a NanoWizard Ultra-speed AFM (JPK, Berlin). The qp-BioAC2 (NanoAndMore) cantilevers were used with a nominal spring constant of 0.1 N/m. Similar to HS-AFM experiments, AIEX-enriched PDHs samples in 100 mM HEPES pH 7.4, 50 mM NaCl, 5 mM KCl, 1 mM MgCl_2_, 5% glycerol were incubated on a freshly cleaved mica surface for 10 min. Substrate-dependent PDHc movements were imaged after the addition of 2 mM ThDP, 4 mM Pyr, 3 mM NAD^+^, and 0.4 mM CoA. The particles were visualized under incubation buffer conditions using QI imaging mode. The nanoindentation experiments were done in closed-loop contact mode with a tip approaching speed of 500 nm/s and an indentation force of 3 nN. Each indented particle was imaged before and after indentation. The acquired images and force-distance (F-D) curves were processed using JPK Data Processing Software. Hooke’s law with two springs in series (particle and cantilever) was used in order to extract the particle spring constant^90^.

## Data availability

Cryo-EM maps and their corresponding models were deposited in EMDB and PDB under the following entries: EMD-57265, 29OF (E2 core ground state); EMD-57264, 29OE (E2 core saturated); EMD-57266, 29OG (E3BP); EMD-57263, 29OD (E1). The integrative model of the PDHc is deposited in the PDB-IHM under the accession number 5V82. The following publicly available PDB accessions were used within this study: 2OZL, 6ZZK, and 1EBD.

## Acknowledgements

We thank Ioannis Skalidis for assistance in data analysis, Noah Binay for experimental support, and our laboratory members for useful discussions. Work in the W.H.R. laboratory was supported by the ERC through the Advanced Grant “LabFree” and S.M. was supported by NWO through an ENW-XS grant. This work was supported by the European Research Council (ERC) under the European Union’s Horizon 2020 research and innovation programme through the project “ProMiDis” (grant agreement No. 819934; support for GS), as well as by the Horizon Europe Programme under the “Widening Participation & Spreading Excellence” component, call “HORIZON-WIDERA-2022-TALENTS-01-01–ERA Chairs”, through the project “Boost4Bio” (grant agreement No. 101087471; support for ZFB and GS). Computational resources were provided by AWS-GRNET under the project “Cryo-EM Modelling” and were used in MEMMI. The authors acknowledge the Life Science Electron Microscopy facility at the Ernst-Ruska Centre, Forschungszentrum Jülich, Germany for providing imaging time (proposal number 230712_TK_01). We especially thank Dr. Saba Shahzad for her assistance with SPA analysis and CET imaging of the samples. Work in the C.G. laboratory was supported by the Interdisciplinary Center for Clinical Research (IZKF) Münster; project number Z/005/24. This project has received funding from the European Union’s Horizon 2020 research and innovation programme under grant agreement No 101004806, profiting from two rounds of MOSBRI TNA (MOSBRI-2023-192, MOSBRI-2024-325) This work was supported by the European Union through funding of the Horizon Europe ERA Chair “hot4cryo” project number 101086665 (to PLK), the Federal Ministry for Education and Research (BMBF, ZIK program; Grant nos. 03Z22HN23 and 03COV04 to PLK), the European Regional Development Funds for Saxony-Anhalt (grant no. EFRE: ZS/2016/04/78115 and ZS/2024/05/187255 to PLK), the Federal State Saxony-Anhalt (to PLK), funding by DFG (project number 391498659, RTG 2467 to PLK; project number 421152132, SFB 1664 to PLK; project number SA 494 11/1 to RGS) the region of Saxony-Anhalt, and the Martin-Luther University of Halle-Wittenberg (to PLK).

## Author information

### Contributions

T.K.T. and P.L.K. conceptualized the project. T.K.T. and F.L.K. performed protein purification and grid preparation. T.K.T. and P.L.K. model building and SPA image analysis. T.K.T. and G.K. performed the CET image analysis. T.K.T., P.L.K., and C.T performed integrative analysis. T.K.T, F.H., and S.S. screened and acquired data collection of the purified PDHc. A.N. and A.B. performed cryo-FIB milling and tilt series acquisition of cellular samples. S.M. performed and analyzed HS-AFM measurements. Z.F.B. performed MEMMI simulations. C.S., C.G., G.S., W.R., and P.L.K. supervised the study.

