## Supplementary Figures, Tables and Movies for "A nested shell structure coordinates enzyme communication in pyruvate oxidation"

**This PDF file includes:**

Extended Data Figures 1 to 52
Extended Data Tables 1 to 10
Extended Movie 1 to 5
Supplementary References

#### Extended Data Figures

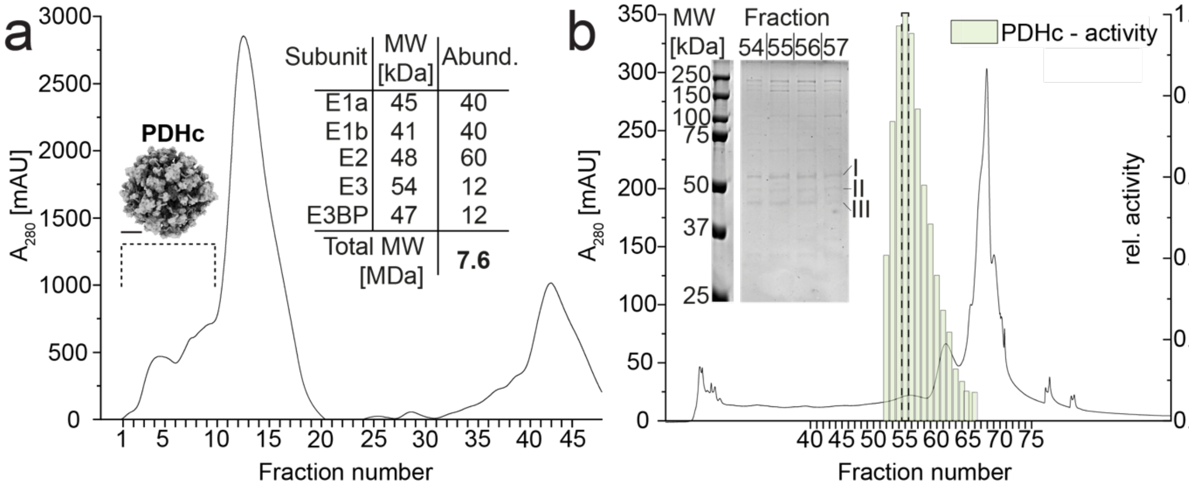


**Extended Data Fig. 1. Tandem purification of the endogenous PDHc.**

**a**, SEC elution profile, displaying absorption at 280 nm against elution volume, of *T. thermophila* cell extract is displayed. The estimated molecular mass of the complex (~7.6 MDa) was calculated based on the known subunit composition and stoichiometry of E1α, E1β, E2, E3, and E3BP[1], as summarized in the inset table. Fractions corresponding to the high-molecular-weight shoulder (fractions 1-10, indicated by dashed lines) were pooled for subsequent affinity capture (scale bar is 10 nm). **b,** AIEX of the pooled SEC fractions was performed, and protein complexes were eluted using an increasing NaCl gradient. The absorbance at 280 nm is plotted as a function of the elution volume, while the normalized enzymatic activity is shown for each corresponding fraction. SDS-PAGE of the most catalytically active fractions displays multiple bands in the molecular ranges of E1α/β (I), E2/E3BP (II), and E3 (III).

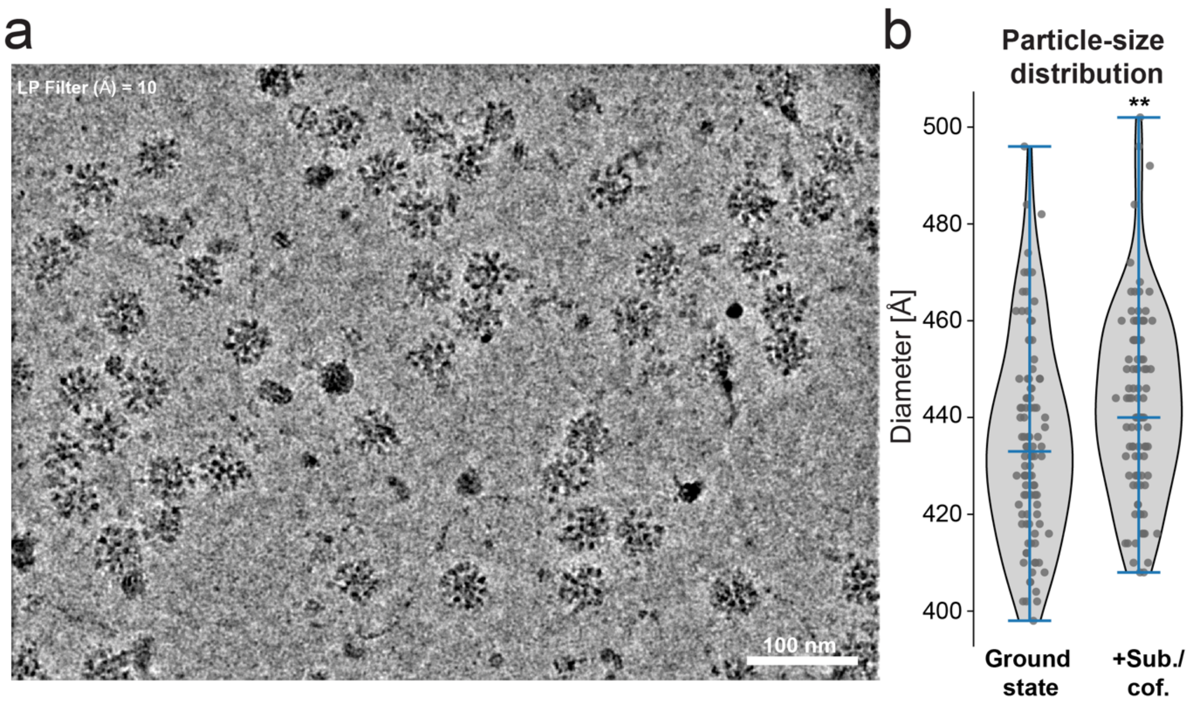


**Extended Data Fig. 2. Characterization of the enriched PDHc sample.**
**a,** A representative low-pass filtered micrograph of the enriched PDHc fraction, displaying the homogeneity of the enriched sample, is shown. **b,** Particle-size distribution of the PDHc was measured from cryo-EM micrographs for the ground- and saturated state. The average particle diameter was 435 ± 21 Å (N = 100) for the ground state and 442 ± 19 Å (N = 100) for the saturated state.

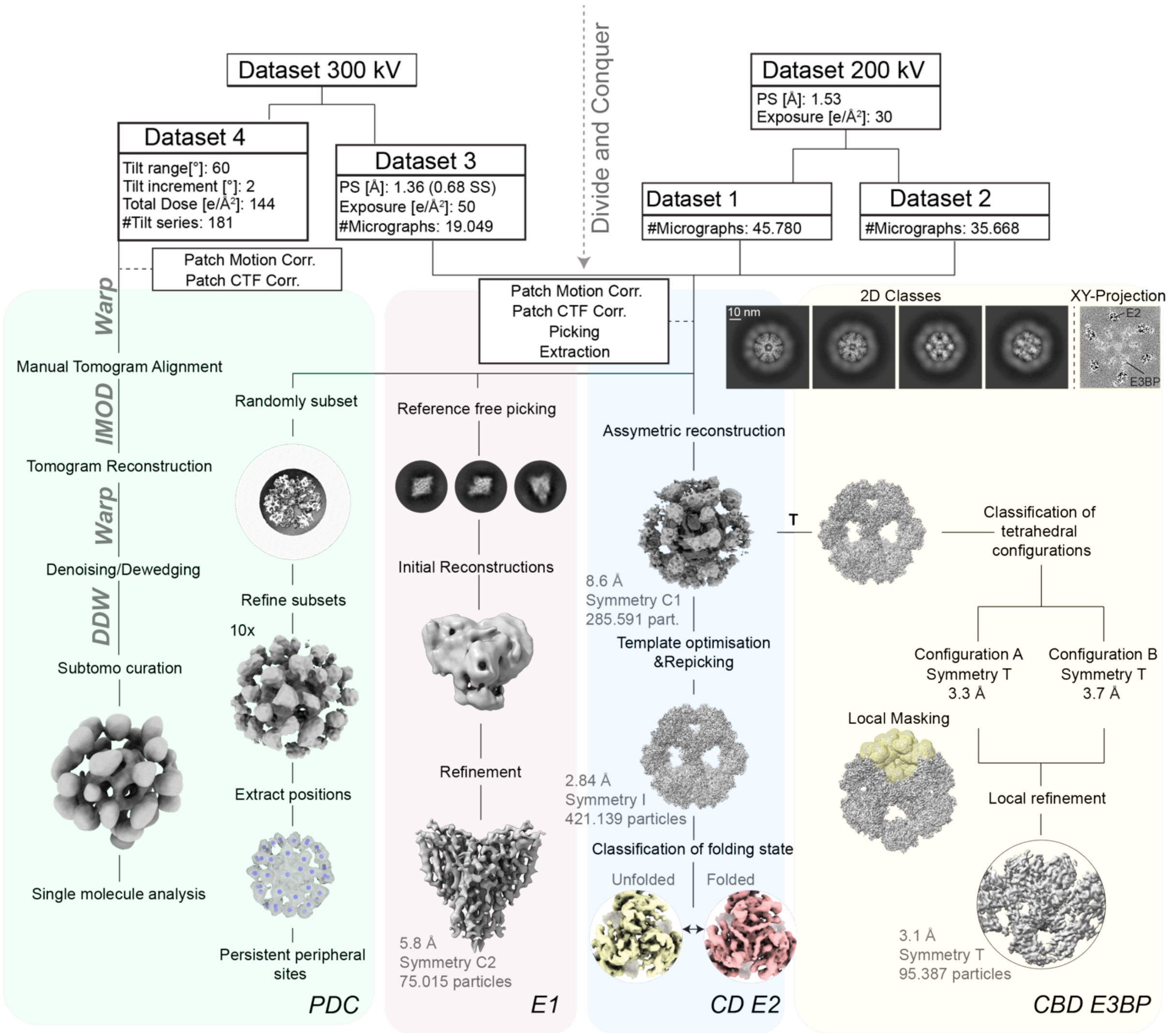


**Extended Data Fig. 3. Resolving the endogenous PDHc.**

A schematic representation of the image analysis pipeline for the reconstruction of the full PDHc is shown

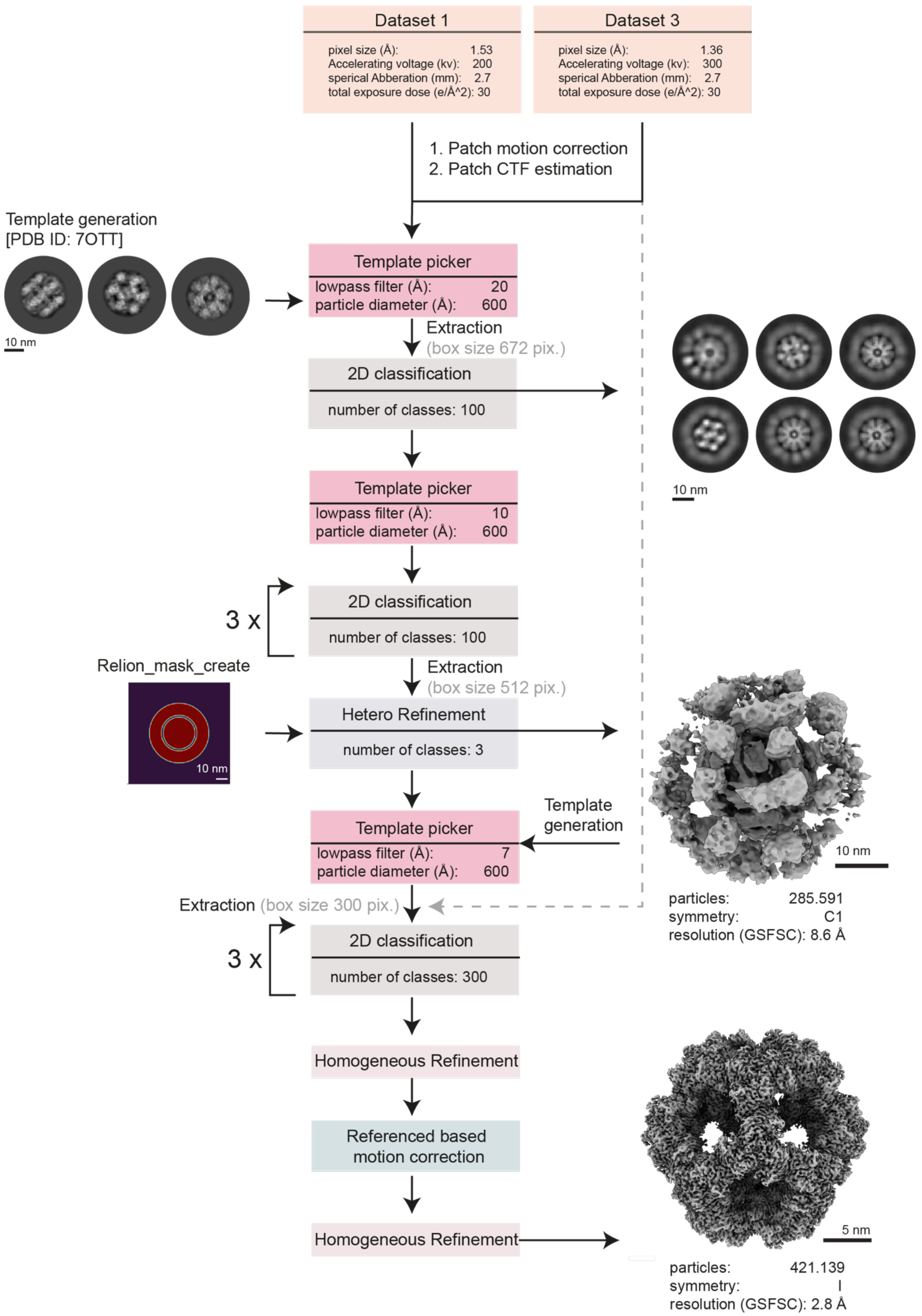


**Extended Data Fig. 4. Reconstruction of the E2 CD.**
Schematic representation of the image analysis pipeline for the E2 CD.

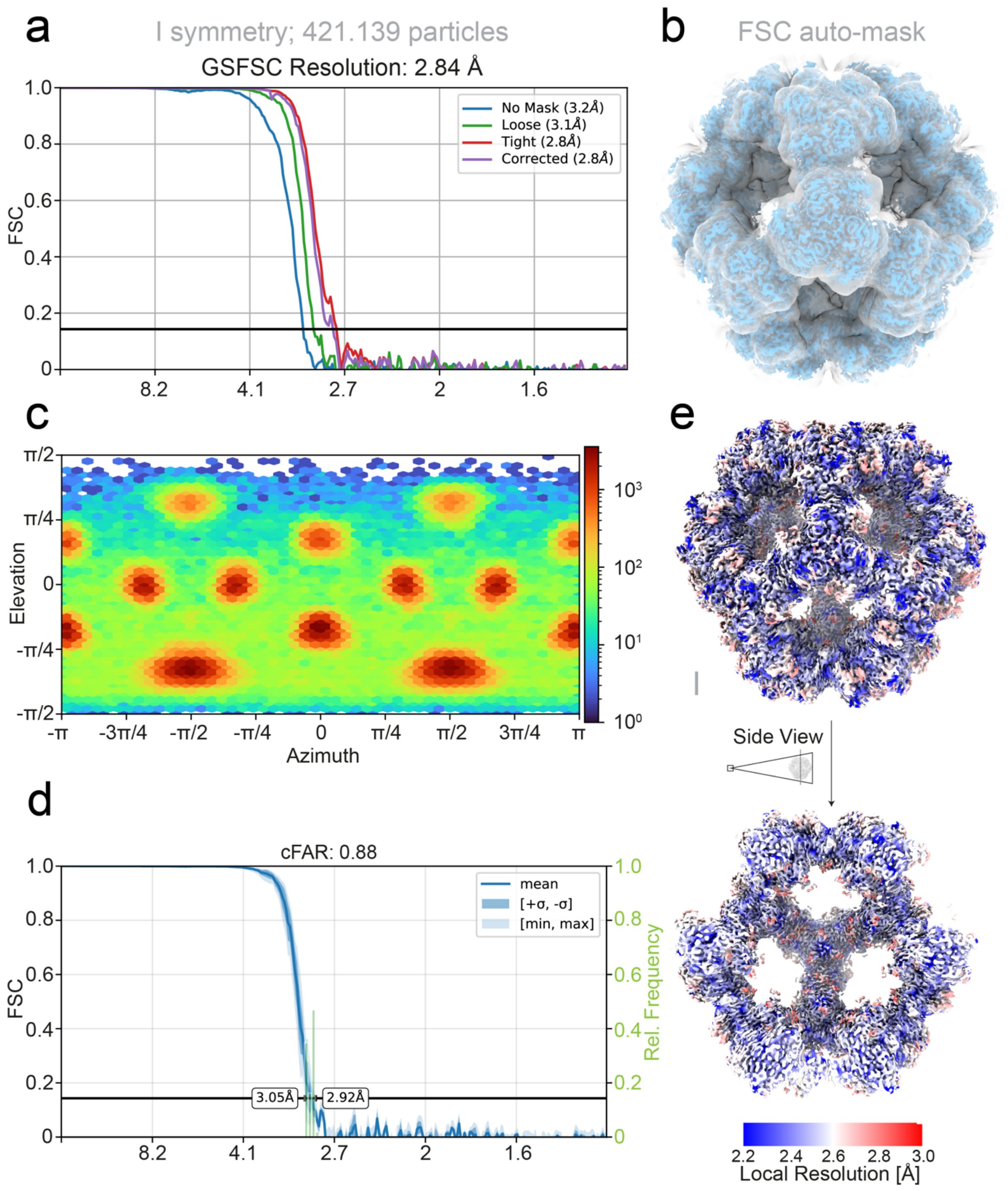


**Extended Data Fig. 5. Quality metrics of the E2 CD reconstruction.**
**a,** The E2 CD of the PDHc was reconstructed at 2.84 Å (FSC = 0.143, I symmetry) using 421,139 particles. **b,** The mask used for structure refinement is displayed. **c-d,** The overall view coverage and the directional FSC (3D FSC) [2] are shown. **e,** The calculated local resolution estimation shows a distribution between 2.2 and 3 Å (contour level σ 0.2).

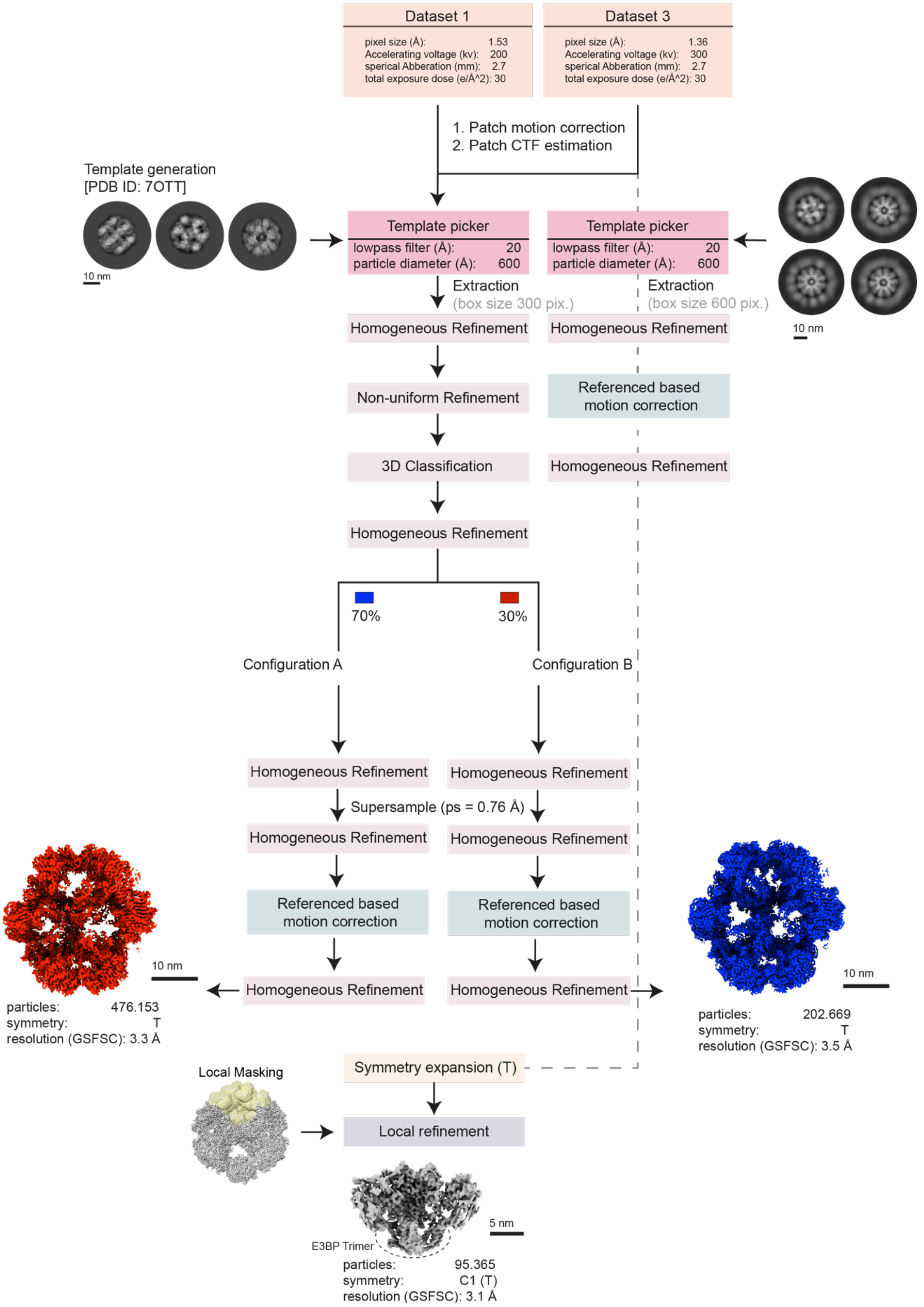


**Extended Data Fig. 6. Reconstruction of the E3BP CBD.**
Schematic representation of the image analysis pipeline for the E3BP CD.

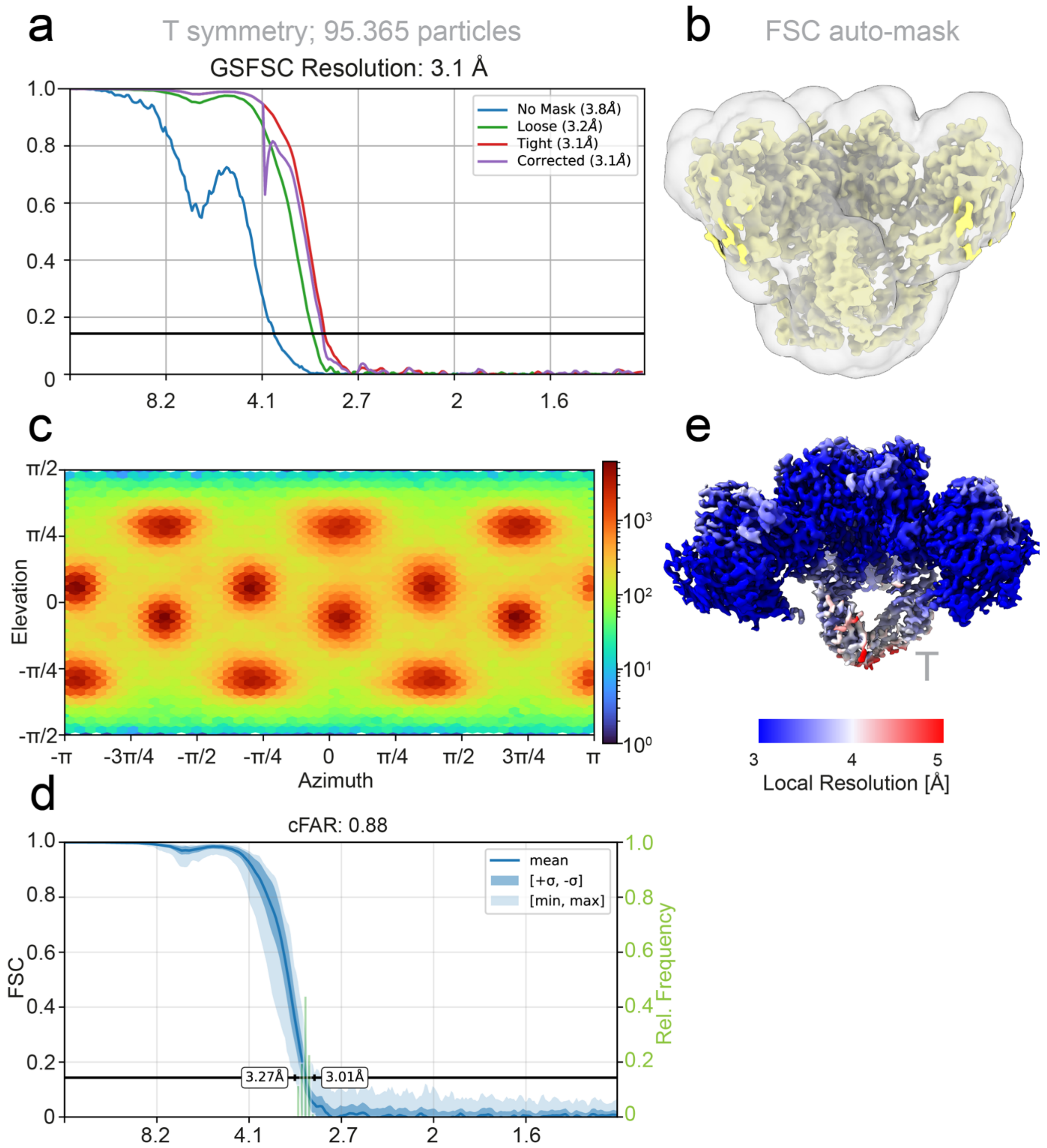


**Extended Data Fig. 7. Quality metrics of the E3BP CBD reconstruction.**
**a,** The E3BP of the PDHc was reconstructed at 3.1 Å (FSC = 0.143, T symmetry) using 95,365 particles. **b,** The mask used for structure refinement is displayed. **c-d,** The overall view coverage and the directional FSC (3D FSC) [2] are shown. **e,** The calculated local resolution estimation shows a distribution between 3 and 5 Å (contour level σ 0.2).

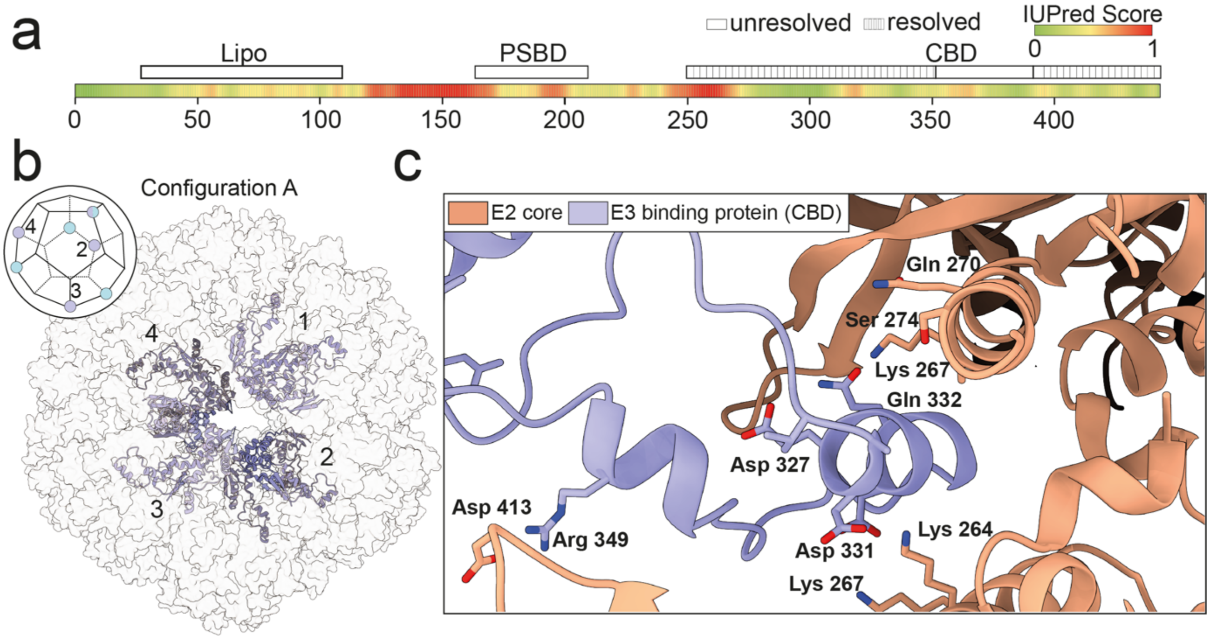


**Extended Data Fig. 8. Architecture of the E3BP scaffold.**

**a,** Each E3BP contains three distinct domains connected by flexible linker regions (IUPred score for local disorder given[3]). **b,** The E3BP complex can adopt two symmetrically exclusive tetrahedral configurations within the E2 CD. Approximately 70% of trimers adopt configuration A (purple), while 30% adopt configuration B (cyan), displaying a mixed structural population within the dodecahedral scaffold. The atomic model of the E3BP (Configuration A), consisting of 4 trimeric E3BP building blocks, is shown inside the dodecahedral E2 CD. **c,** The E3BP interacts with the E2 catalytic domain (orange) at the intertrimeric interface, where the local resolution enables visualization of the atomic-level contact surface. The E2–CBD interaction interface is shown, displaying the hydrogen-bond and electrostatic network stabilizing the transient association. Key residues contributing to the interaction include Lys 264, Lys 267, Asp 327, Asp 331, and Arg 349 on the E2 core, and Asp 413, Gln 270, and Ser 274 on the CBD, forming a charge-stabilized contact surface that anchors the flexible E3-binding domain at the core periphery.

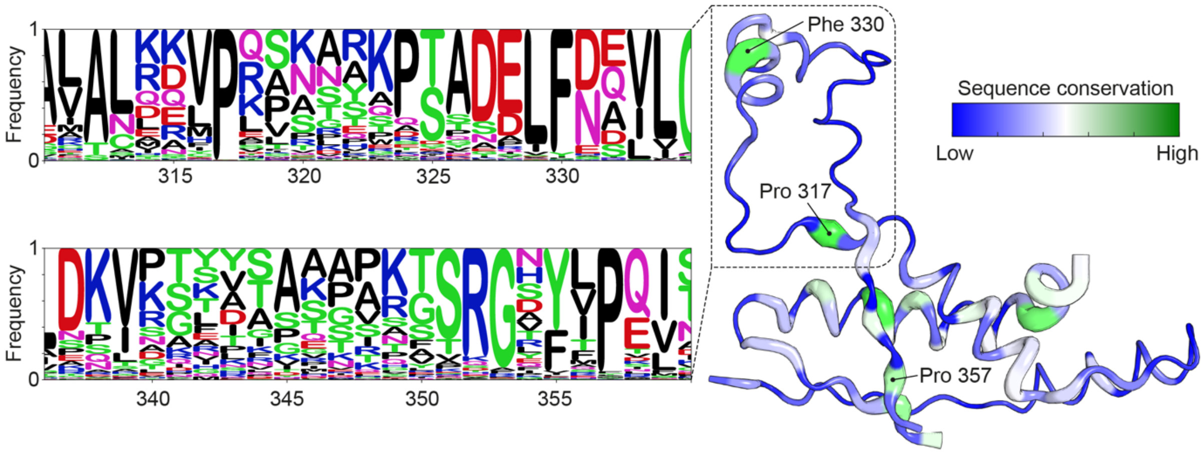


#### Extended Data Fig. 9. Structural conservation of the E3BP.

To identify the conserved regions responsible for mediating E2-E3BP association, the sequence conservation of fungal proteins was examined using the MSA implementation of AF2 via ConservedFold [4]. The analysis reveals a highly conserved structural core responsible for forming the E3BP trimer, along with a short helical segment (residues 326–334) that contributes directly to the E2–E3BP interaction interface. Conservation is highlighted in the sequence logo plot of the E3BP MSA (left) and mapped to the atomic model.

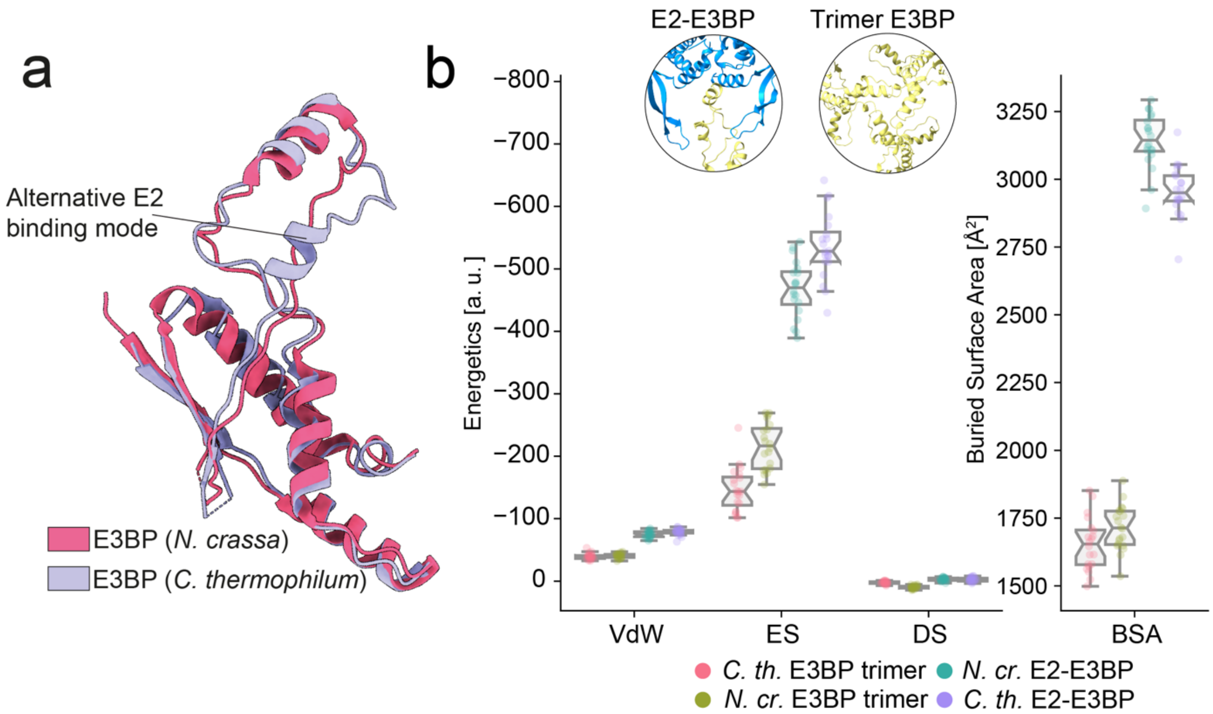


**Extended Data Fig. 10. Structural polymorphism in the fungal E3BP.**
**a,** The structural alignment of the E3BP CBDs from *T. thermophila* (purple) and *N. crassa* (pink) [5] reveals overall high structural conservation, with local deviations located at the E2 interfacing domain. **b,** Comparative energetic evaluation of E3BP–E2 and E3BP trimer interfaces from *T. thermophila* and *N. crassa* was conducted using the HADDOCK refinement server [6]. Energetic contributions are separated into van der Waals, electrostatic, and desolvation energies, shown in arbitrary units, alongside the corresponding buried surface area of the interface. Electrostatics, Desolvation, Van der Waals, and buried surface area were plotted using the Haddock Plotter package [7].

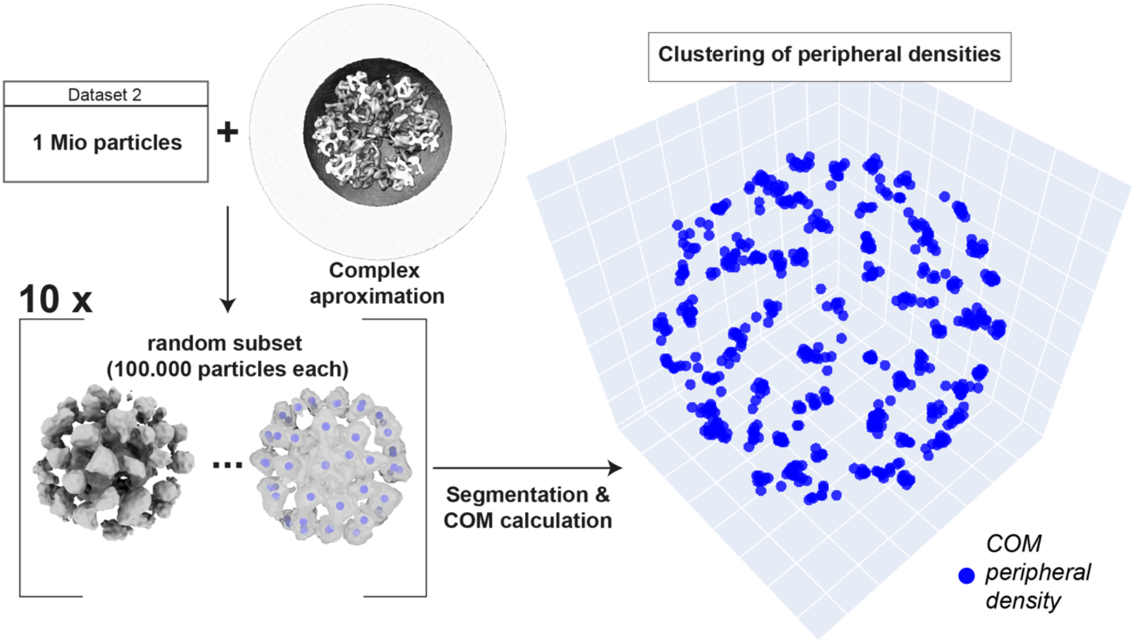


**Extended Data Fig. 11. Asymmetric reconstructions of the PDHc.**
A total of 1 million particles from Dataset 2 were used to approximate the overall complex architecture. To assess the reproducibility of peripheral density localization, ten random subsets of 100,000 particles each were independently refined. Volumes were segmented, and the COMs of peripheral densities were calculated. The resulting distribution reveals consistent clustering of peripheral densities around the E2/E3BP core, indicating reproducible positioning of peripheral subunits such as E1 and E3 across independent reconstructions.

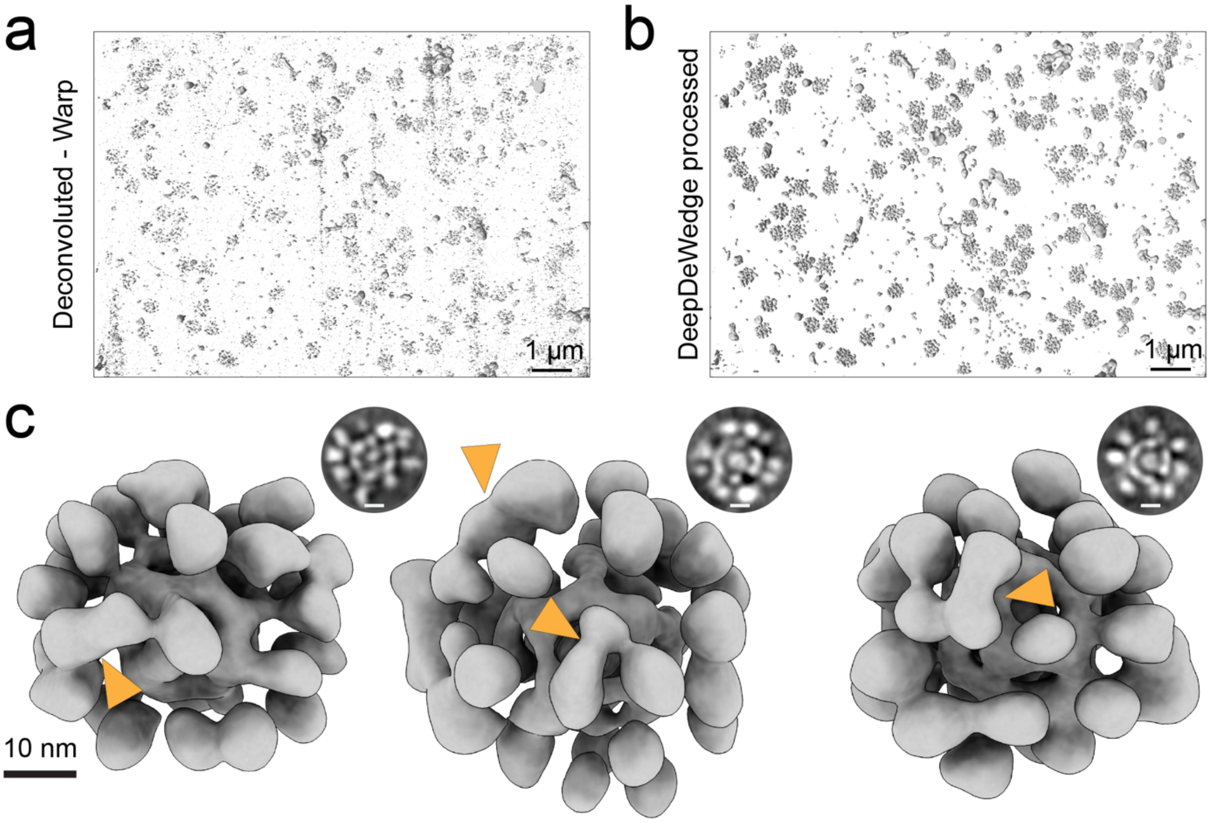


**Extended Data Figure 12: Gallery: single molecule PDHc.**
**a,** Example tomogram after deconvolution with the tomography package Warp. **b,** The same tomogram after denoising and dewedging using the U-net–based DeepDeWedge neural network. **c,** Representative denoised subtomograms are shown with their respective X-Y slice. Regions where subunits are in close proximity, potentially indicating local interaction sites relevant to reaction cycle completion, are highlighted in orange

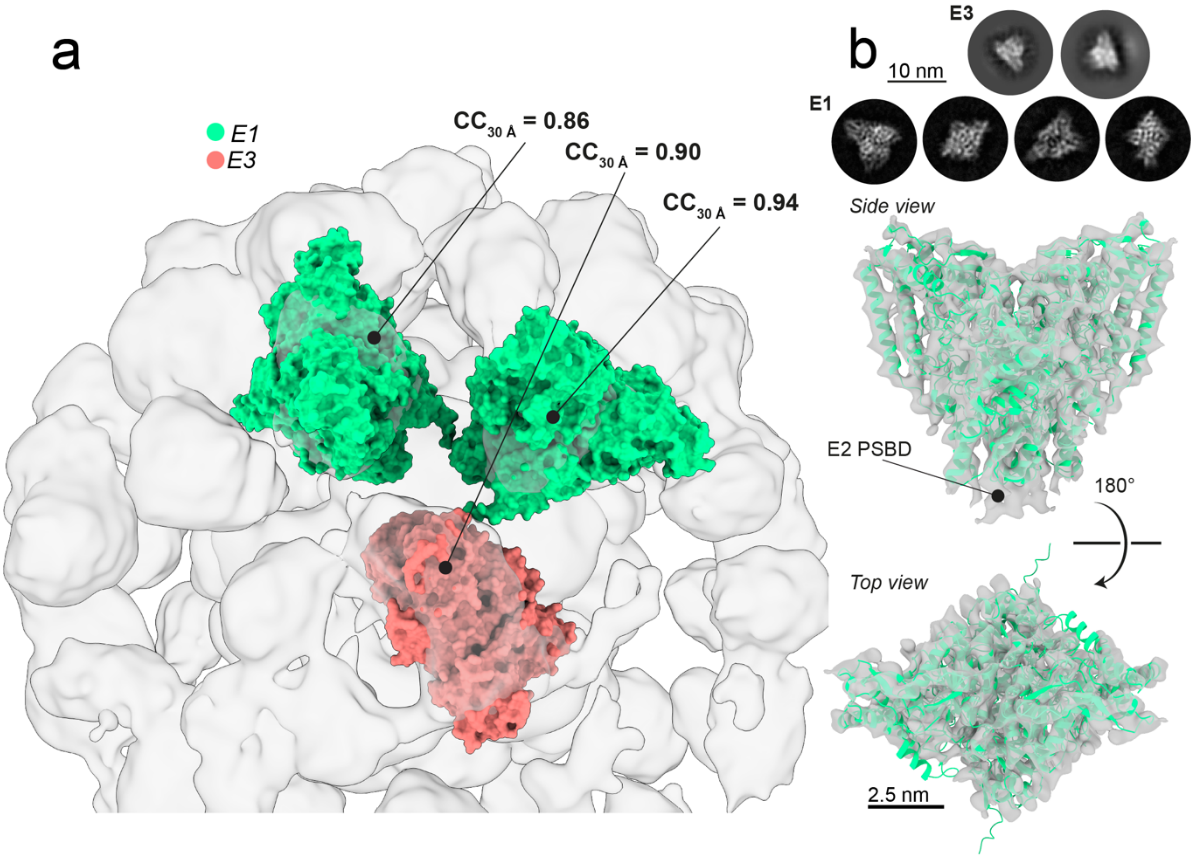


**Extended Data Figure 13: The peripheral subunits of the PDHc.**
**a,** Systematic correlation-based fitting of E1 (green) and E3 (red) models into the peripheral cryo-EM densities revealed discrete binding positions with high correlation coefficients ($CC_{30\,Å}$ = 0.86–0.94). The highest-scoring positions for each subcomplex were subsequently used for placement of the peripheral subunits within the integrative all-atom model of the complete PDHc. **b,** 2D-class averages for the E1 and E3 complexes of the PDHc are shown. The E1 domain of the PDHc was reconstructed from the endogenous intact PDHc. Rigid-body fitting of the AF2-predicted E1 model (green) into the experimental Coulomb potential map (grey) revealed a consistent domain arrangement, closely matching both the predicted architecture and known structural orthologs. A distinct density corresponding to the E2 PSBD was identified near the C2 symmetry axis.

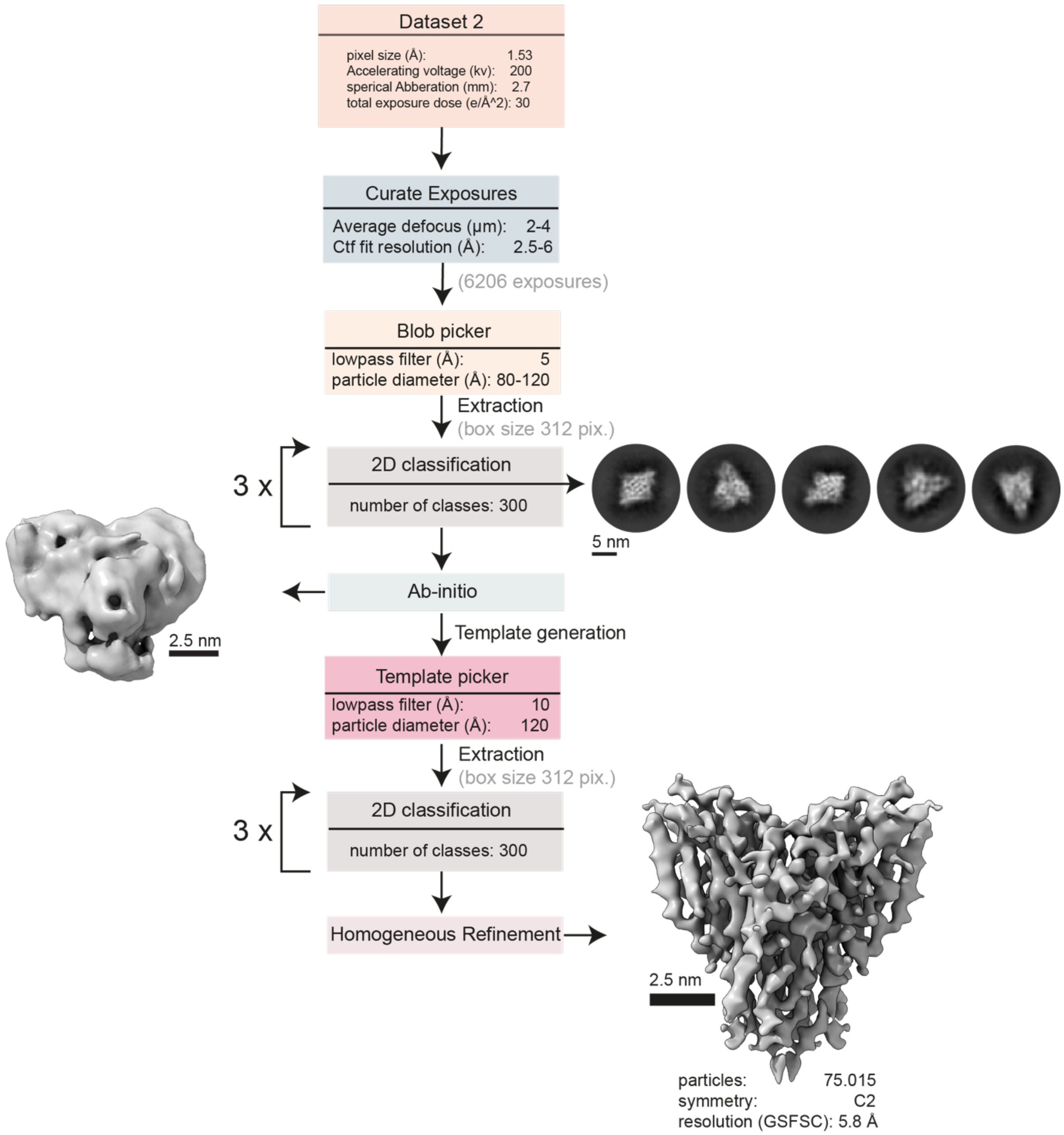


**Extended Data Figure 14: Reconstruction of the E1.**
Schematic representation of the image analysis pipeline for the E1.

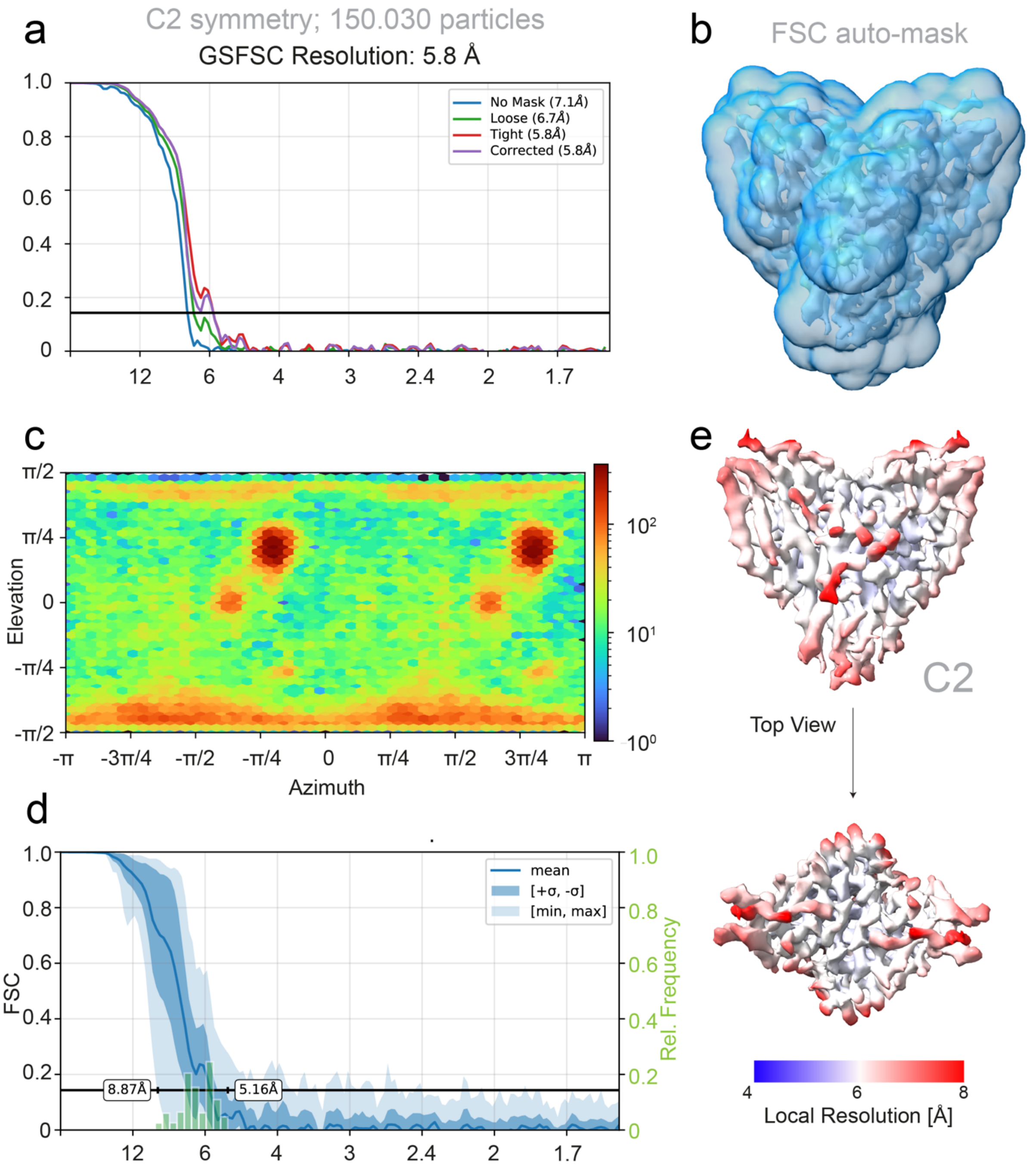


**Extended Data Figure 15: Quality metrics of the E1 reconstruction.**
**a**, The E1 of the PDHc was reconstructed at 5.8 Å (FSC = 0.143, C2 symmetry) using 150035 particles. **b**, The mask used for structure refinement is displayed. **c-d**, The overall view coverage and the directional FSC (3D FSC) [2] are shown. **e,** The calculated local resolution estimation shows a distribution between 5 and 8 Å (contour level σ 0.2).

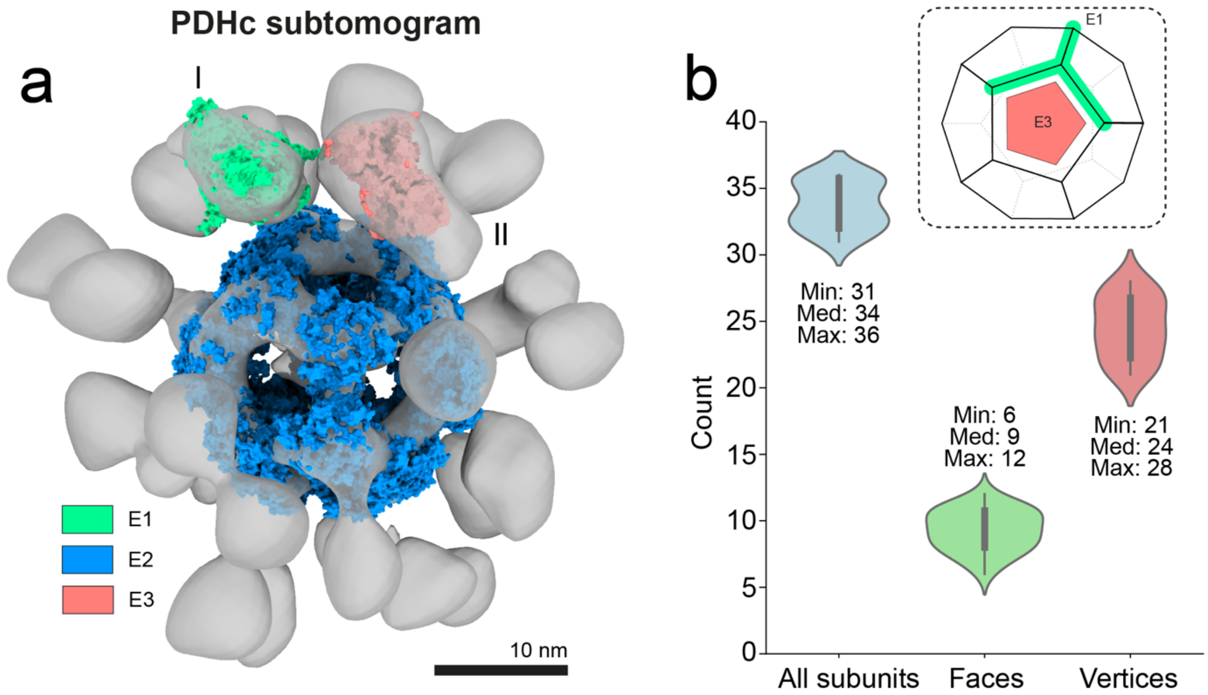


**Extended Data Figure 16: Stoichiometry of the native PDHc.**
**a,** AI-based denoising enabled the *in vitro* analysis of PDHc directly from subtomograms of the enriched complex. The extracted subtomograms display clear signatures of the dodecahedral core, surrounded by peripheral densities related to the E1 (green) and E3 (orange) subunits. **b,** Peripheral densities were manually assigned to the faces and vertices of the dodecahedral scaffold based on their spatial orientation relative to the central core. This analysis yielded approximate stoichiometries for the endogenous complex, indicating 31-36 peripheral densities per particle. On average, 24 densities were localized to face positions and 9 to vertex positions, corresponding closely to the expected distribution of E1 and E3 subunits, respectively.

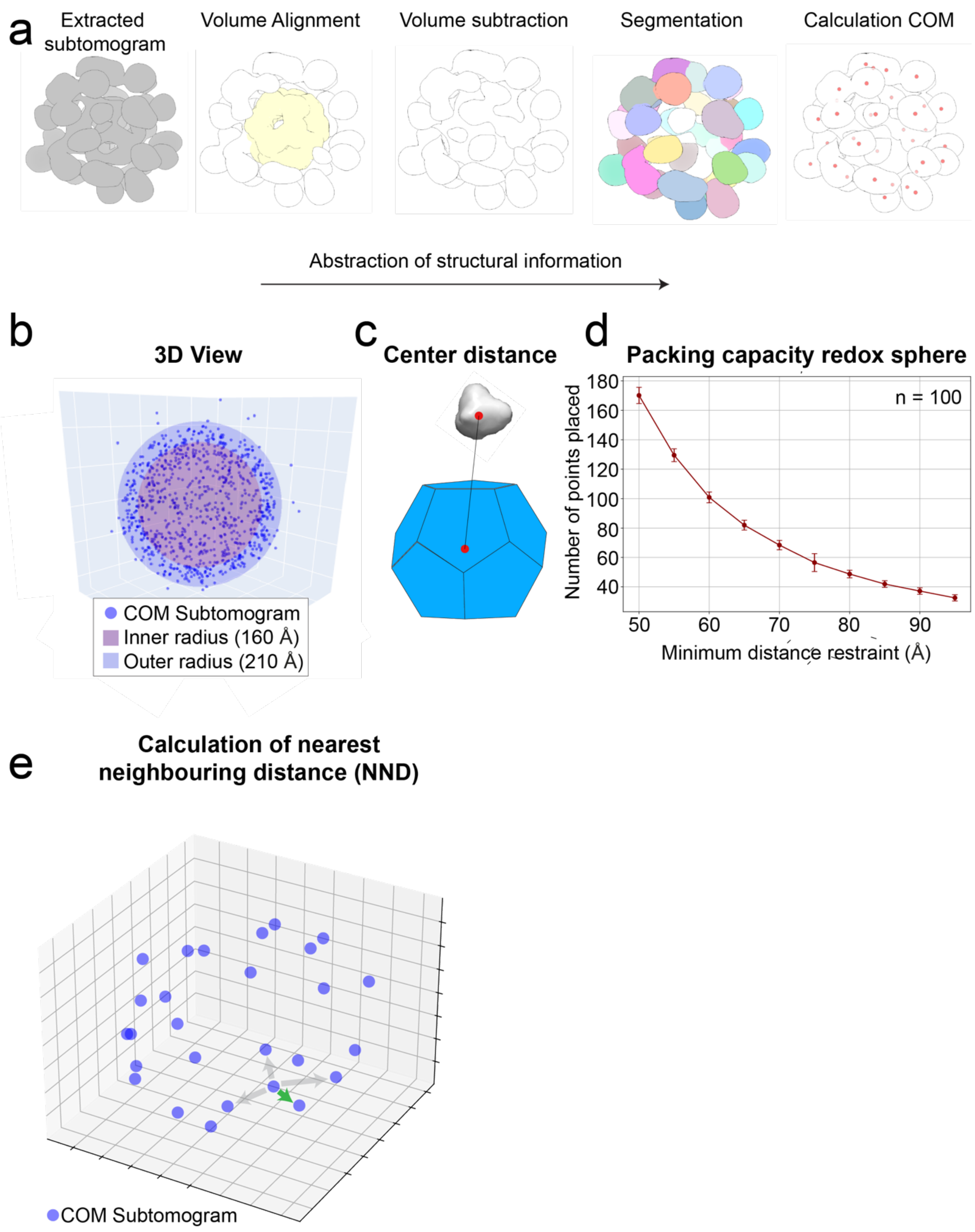


**Extended Data Figure 17: Analysis of PDHc sub-tomograms.**
**a,** Sub-tomograms are extracted from the denoised full tomograms, aligned based on their icosahedral core signature, which is subsequently removed. The peripheral densities are then segmented, and the COM is calculated. **b,** 90% of extracted points are contained within a hollow sphere with the dimensions of ($r_{inner}$ = 160 Å, $r_{outer}$ = 210 Å), as can be calculated by their distance from the symmetry center (**c**). **d,** To probe the maximum packing of the redox sphere, different exclusion criteria were sampled (n=100) for the maximum number of placeable points. **e,** To understand PDHc inter-subunit spacing, the NND between 2 COM points was calculated.

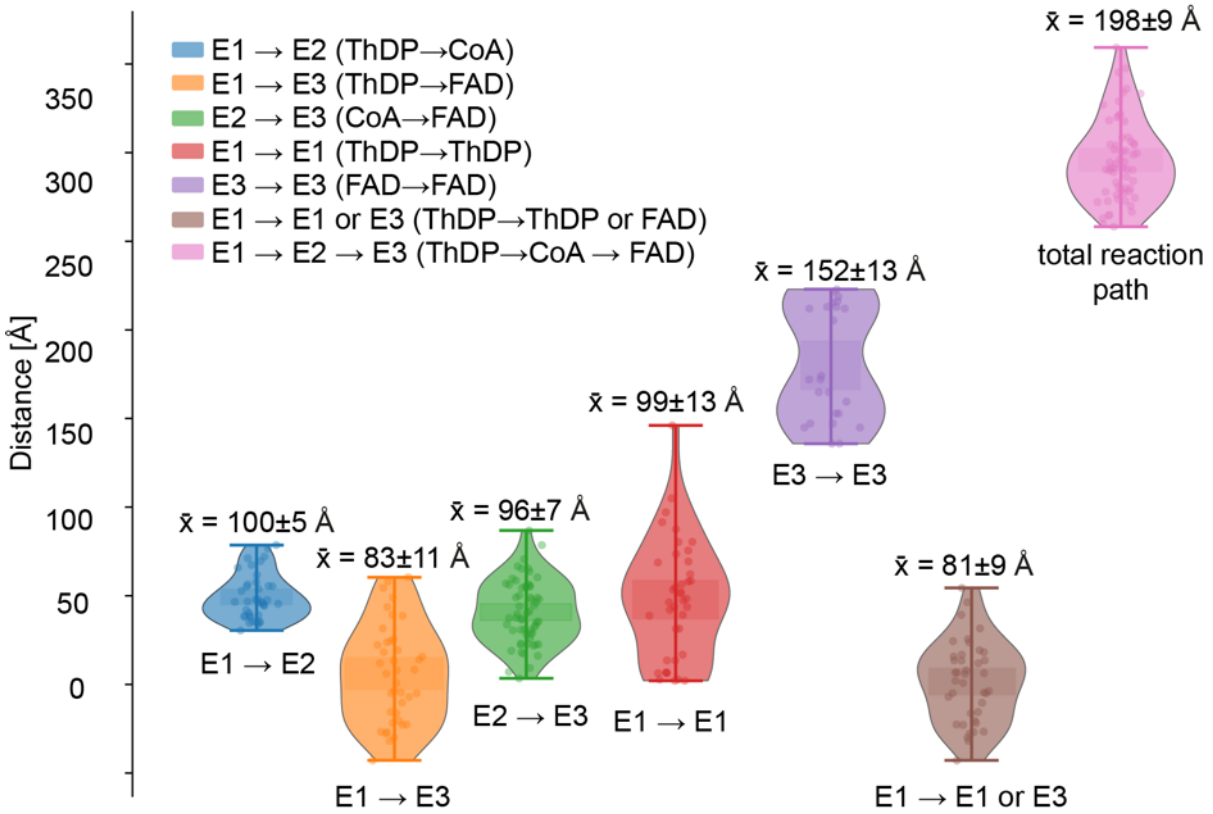


**Extended Data Figure 18: Structural Mapping of the PDHc active sites.**
Violin plots show the distributions of distances between catalytic centers along the enzymatic reaction pathway. Each color corresponds to a specific transfer step between subunits: E1→E2 (ThDP→CoA, blue), E1→E3 (ThDP→FAD, orange), E2→E3 (CoA→FAD, green), E1→E1 (ThDP→ThDP, red), E3→E3 (FAD→FAD, purple), E1→E1 or E3 (ThDP→ThDP or FAD, brown), and the overall E1→E2→E3 transfer path (ThDP→CoA→FAD, pink). Mean reaction distance values ($\bar{x}$) and standard deviations for each distribution are given. The total reaction path (pink) represents the cumulative distance between the three closest cofactor positions (ThDP→CoA→FAD) involved in substrate channeling.

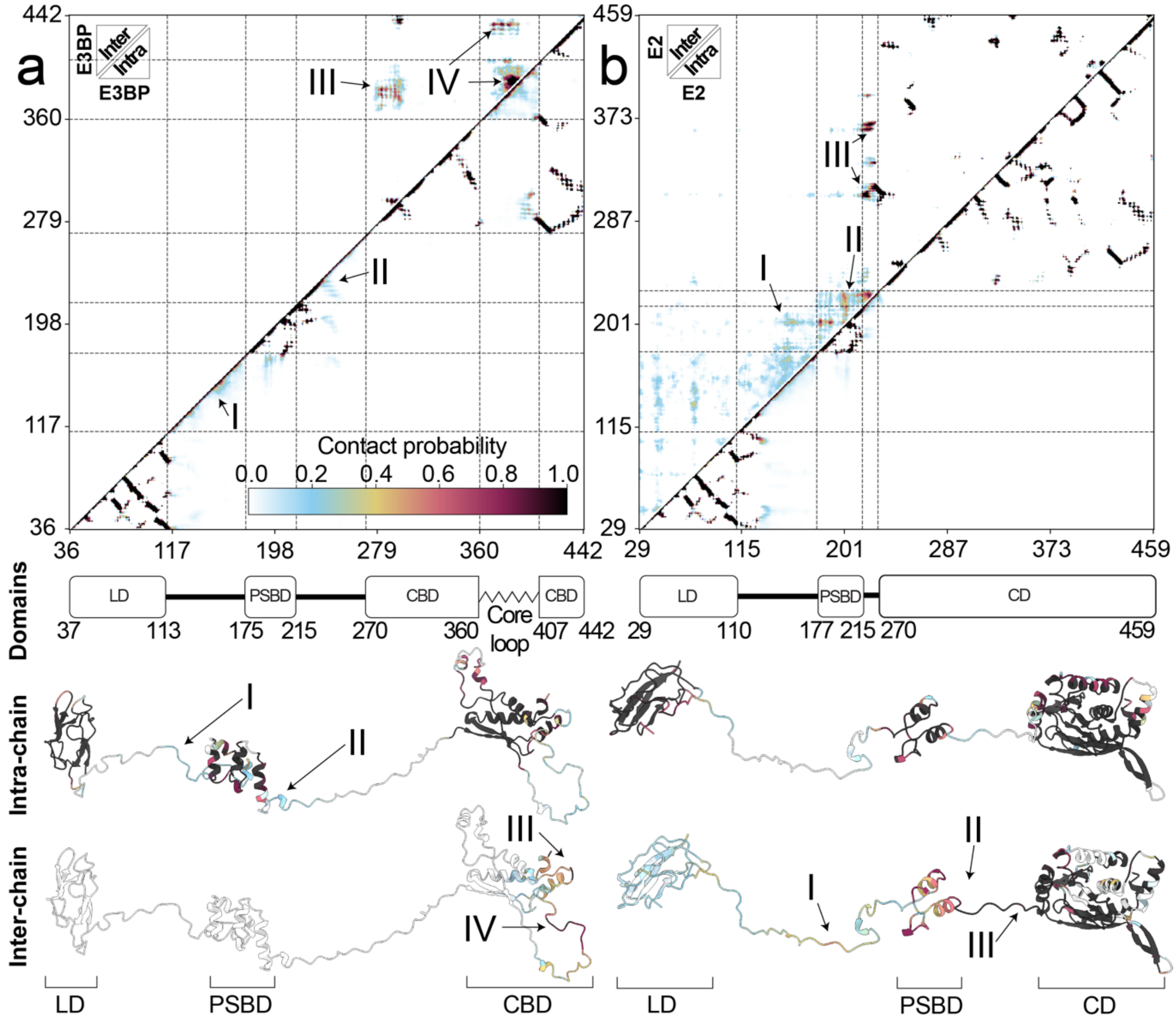


**Extended Data Figure 19: Principal contacts of the flexible regions embedded in PDHc.** Principal residue-residue contacts were computed from the MEMMI simulation of the full PDHc, displaying the resulting contact matrix of the final ensemble. Intra- and inter-chain contact maps were computed for E3BP and E2, with corresponding domain architecture and structural mapping shown below each panel. Contact frequencies are color-coded from low (light blue) to high (black), and major interaction regions (**I-IV**) are annotated. **a,** E3BP flexible regions engage in extensive intramolecular contacts, revealing an E3-binding mode that extends beyond the canonical PSBD (**I–II**). The CBD contains a folded region and a disordered core loop that protrudes into the interior of the dodecahedral core assembly. This region includes a highly conserved DIIDL motif (residues 377-382), which predominantly interacts both with the trimeric E3BP interface (**III**) and with identical motifs on neighboring chains (**IV**). **b,** E2 flexible regions contribute to the structural organization of the complex, stabilizing the positioning of their respective PSBDs (**I**) and forming additional contacts with the lipoyl domain (LD). The E2 PSBDs form inter-chain contacts with neighboring PSBDs (**II**) and additionally interact with other flexible parts linking the PSBD to the CD (**III**), collectively reinforcing higher-order assembly and communication across E2.

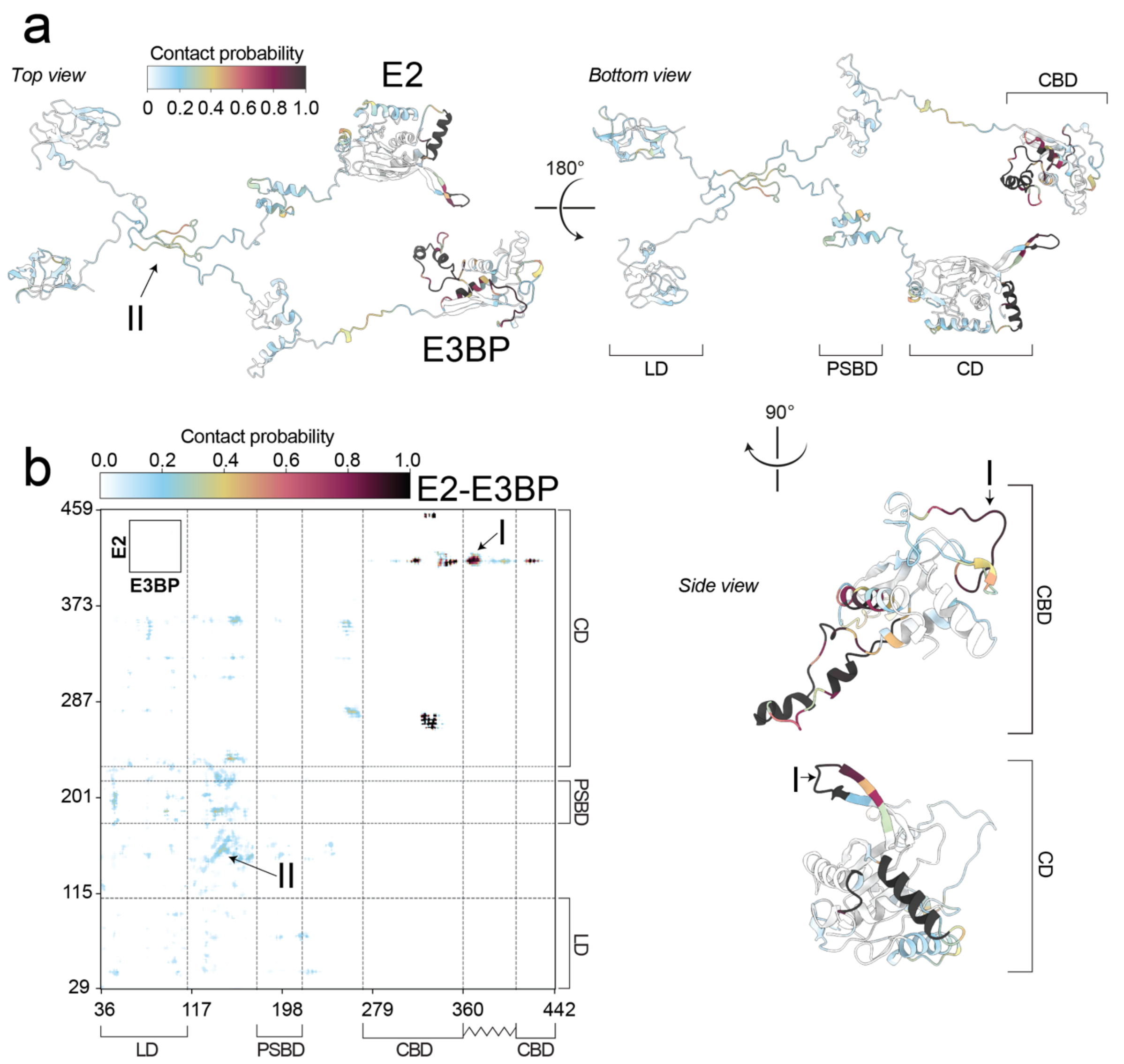


**Extended Data Figure 20: Stabilization of the E3BP assembly.**
Residue-residue interaction patterns of the PDHc MEMMI simulation are displayed. Structural mapping (**a**) and contact matrix (**b**) are shown, highlighting a distinct interaction pattern involving flexible linker regions of the inner core (I) and the nested shell (II). Contact frequencies are color-coded from low (light blue) to high (black), with major interaction regions labeled and E2/E3BP domains annotated.

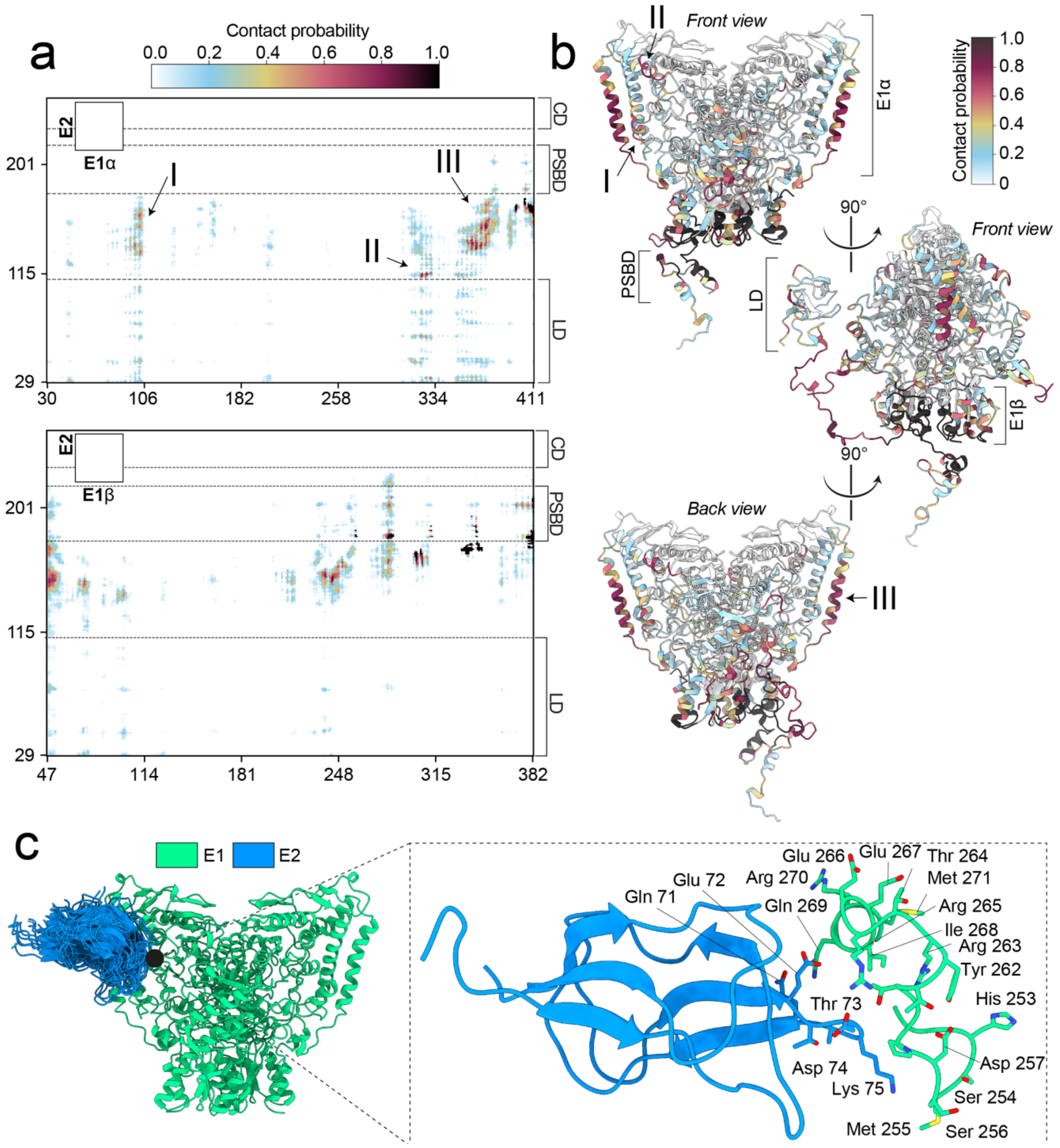


**Extended Data Figure 21: Guiding structural principles of the E2-E1 encounter complex.** To validate the ability to sample relative PPIs related to the initial rate-limiting step of Pyr conversion, contact probabilities from the final MEMMI ensemble were analyzed for the binary E1α/E1β-E2 interaction. **a,** Residue-residue interaction patterns of the PDHc MEMMI simulation are displayed. Contact frequencies are color-coded from low (light blue) to high (black), with major interaction regions labeled and E2 domains annotated. **b,** Structural mapping reveals distinct contacts close to the E1 active site that are sampled by the LD and its organizing flexible regions (**I-III**). **c,** PDHc ensemble structures were examined for conformations in which the lipoylated moiety is positioned near the active-site residues, mimicking feasible insertion complexes. The top 20 scoring models, scored on the distance between Lys 75 and His 253, show a cluster around the E1 active-site pocket, adjacent to the capping loop and catalytic residues (insert).

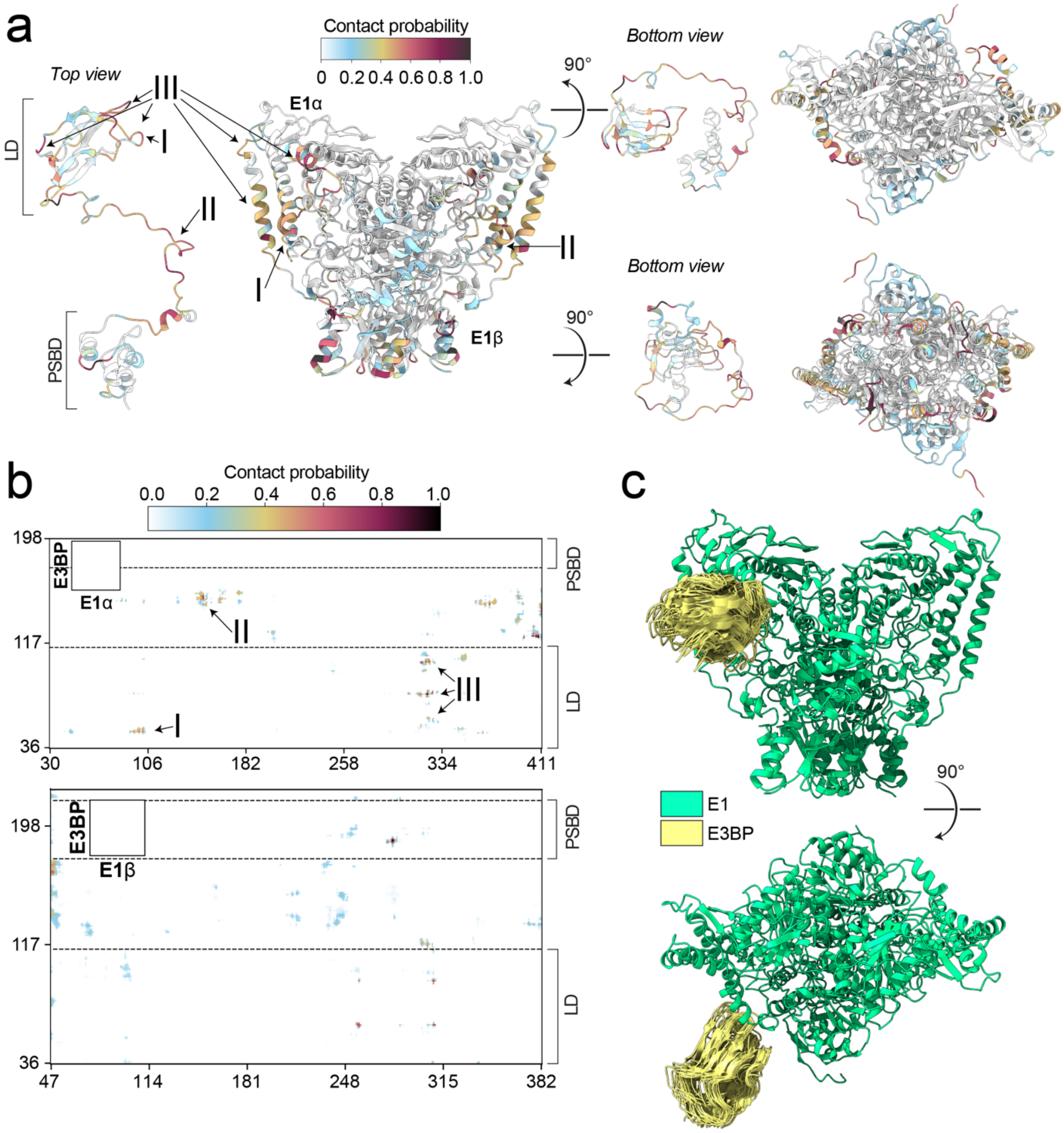


**Extended Data Figure 22: Binary contact map of the E1-E3BP interaction.**
Residue-residue interaction matrix (**a**) and the structural mapping (**b**) of the PDHc MEMMI simulation for the E1-E3BP interaction are displayed. E3BP establishes contacts via both the LD and the flexible LD–PSBD linkers, interacting with E1 residues located in proximity to the active site (**I–III**). Contact frequencies are color-coded from low (light blue) to high (black), with major interaction regions labeled and E2/E3BP domains annotated. **c,** The ensemble was analyzed for E3BP-LD – E1 interactions, highlighting the 20 top-scoring models based on the distance between Lys 75 and His 253. The resulting cluster is located near the E1 active-site pocket.

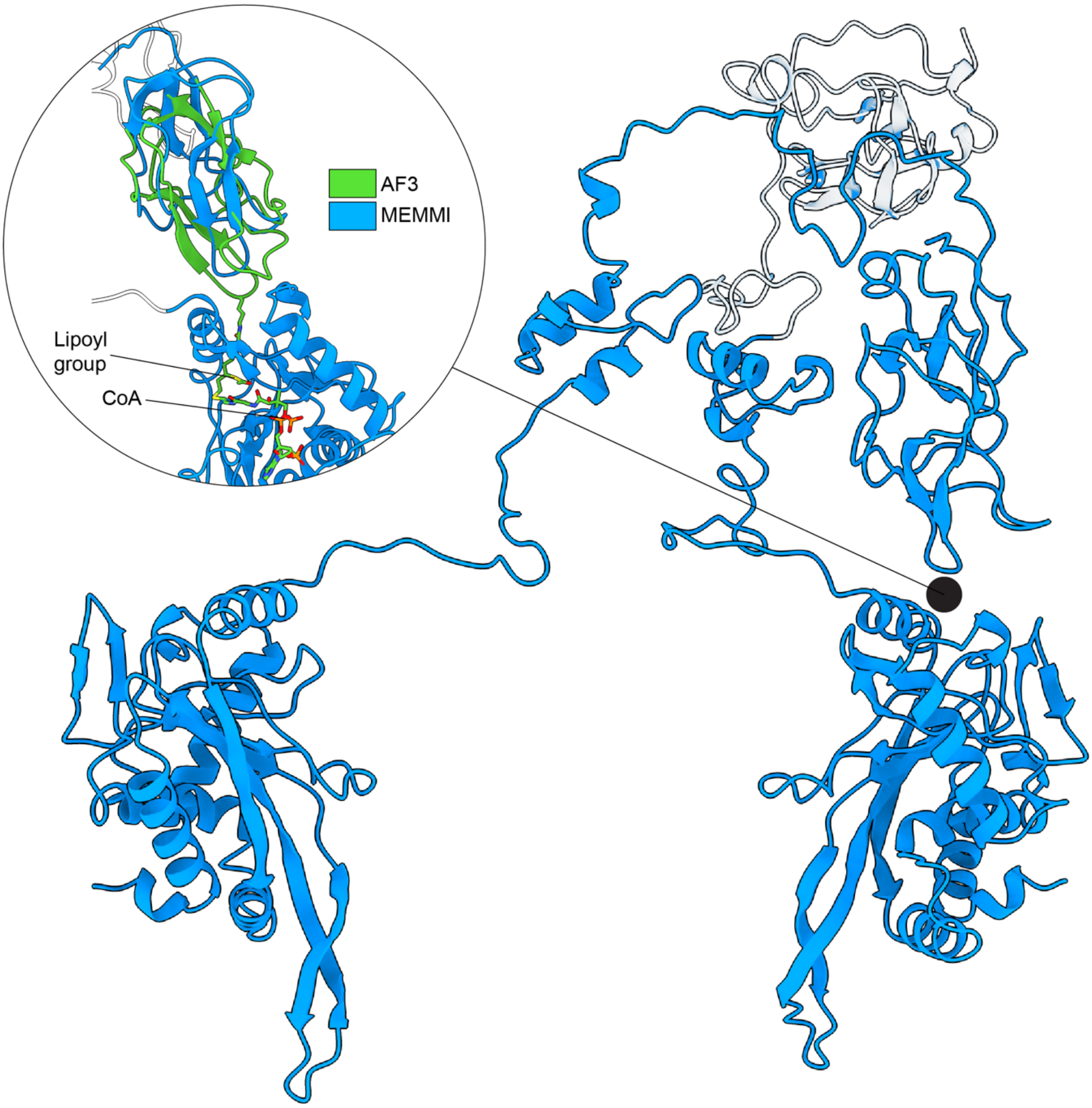


**Extended Data Figure 23: E2 transacetylation complex.**
The ensemble was analyzed for the E2 LD-CD transacetylation complex, highlighting the top-scoring model based on the distance between Lys 75 and Ser 354. The MD-derived model (blue) is closely aligned with the AF3 prediction (green).

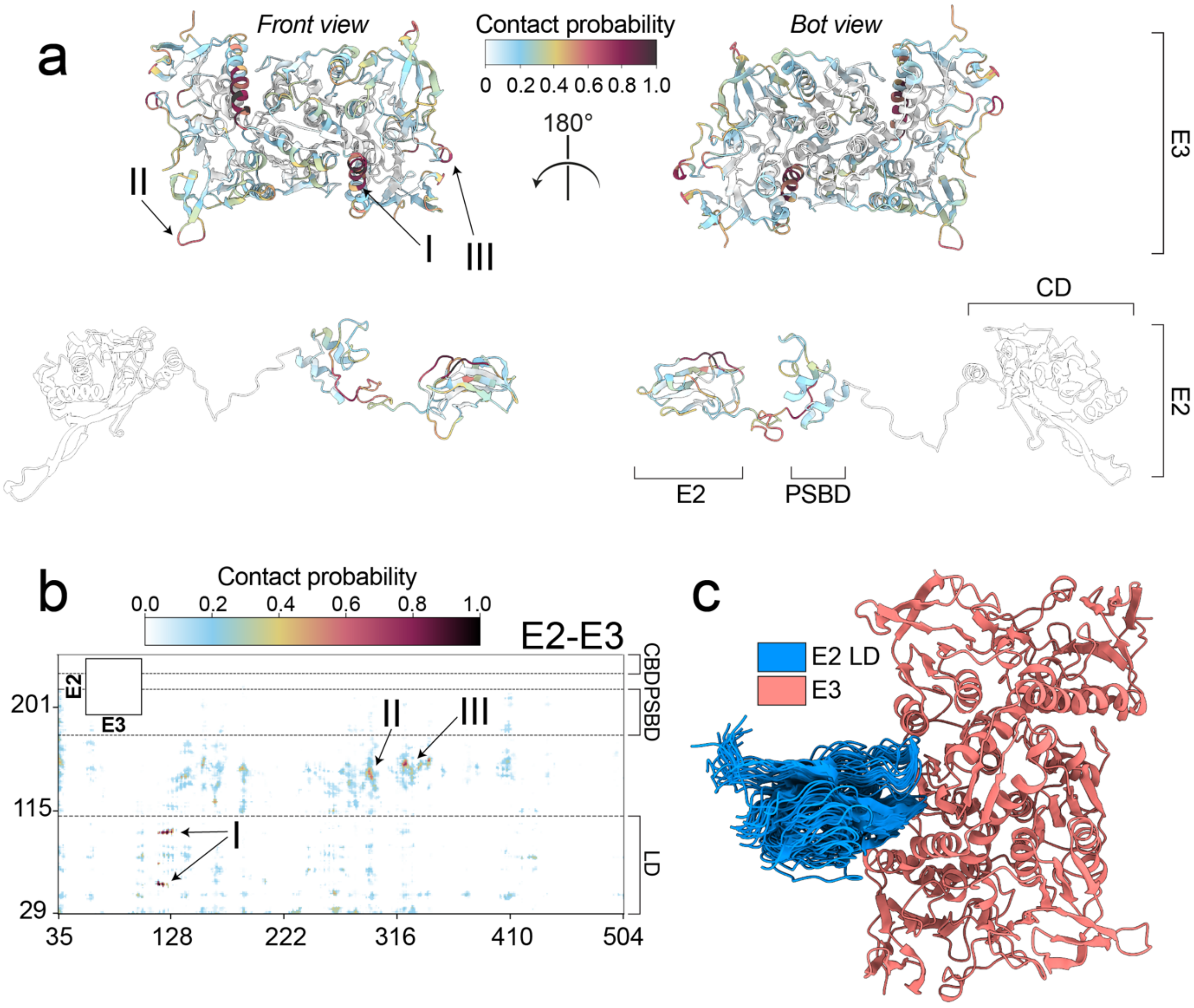


**Extended Data Figure 24: Structural principles of LD regeneration.**
LD reoxidation requires the formation of a transient LD-E3 complex and the insertion of the prosthetic lipoyl arm. **a,** Structural mapping of the interacting regions and contact matrix reveals a distinct pattern of residue contacts. **b,** Contact frequencies are color-coded from low (light blue) to high (black), with major interaction regions highlighted and the E2 domains annotated. **c,** 20 E2 LD domains were chosen for visualization based on the distance between Lys 75 (lipoyl-bearing) and Cys 79, displaying a cluster around the E3 active-site insertion pocket.

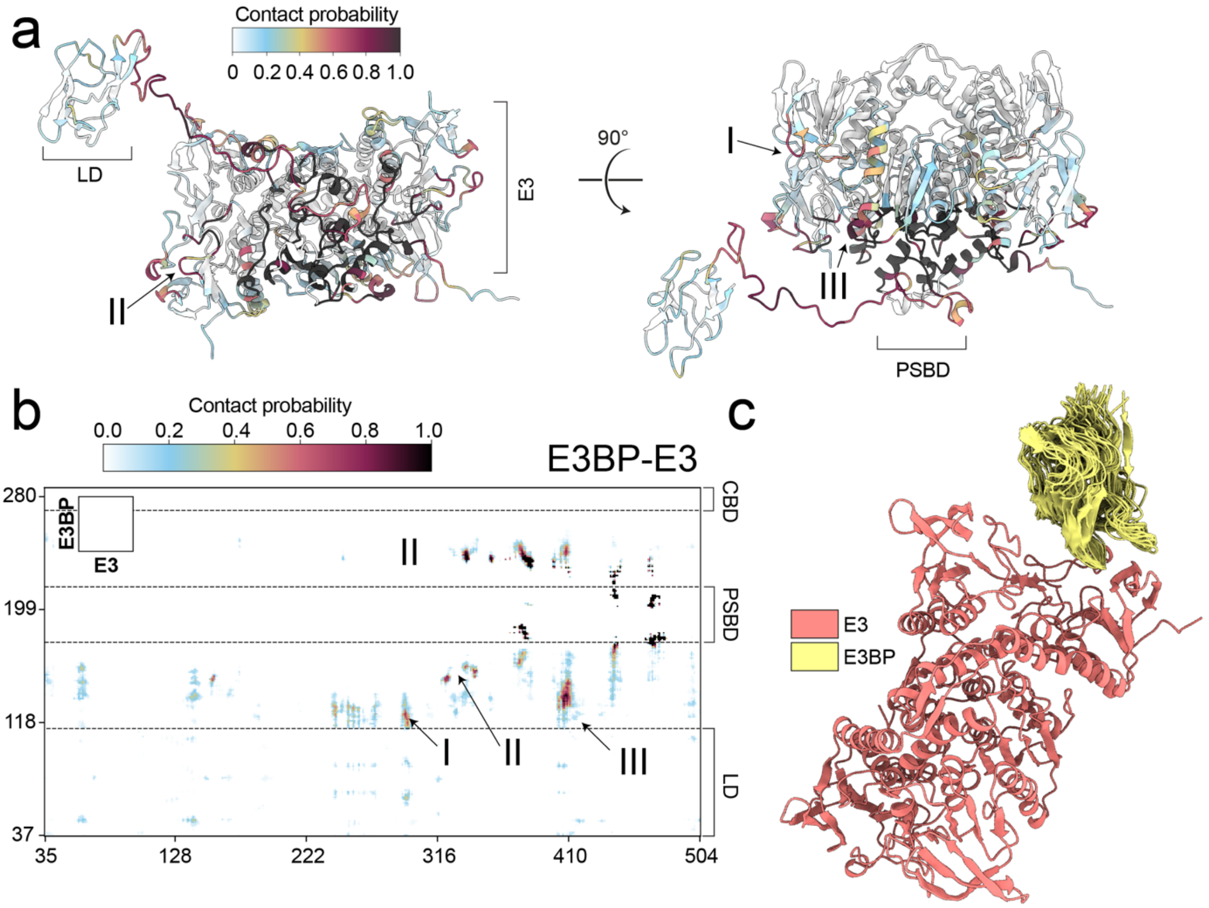


**Extended Data Figure 25: Principal contact map of the E3-E3BP interaction.** Residue-residue interaction patterns of the E3 and E3BP are displayed. Structural mapping (**a**) and contact matrix (**b**) are shown. Contact frequencies are color-coded from low (light blue) to high (black), with major interaction regions labeled. **c,** 20 E3BP LD domains were chosen for visualization based on the distance between Lys 78 (lipoyl bearing) and Cys 79.

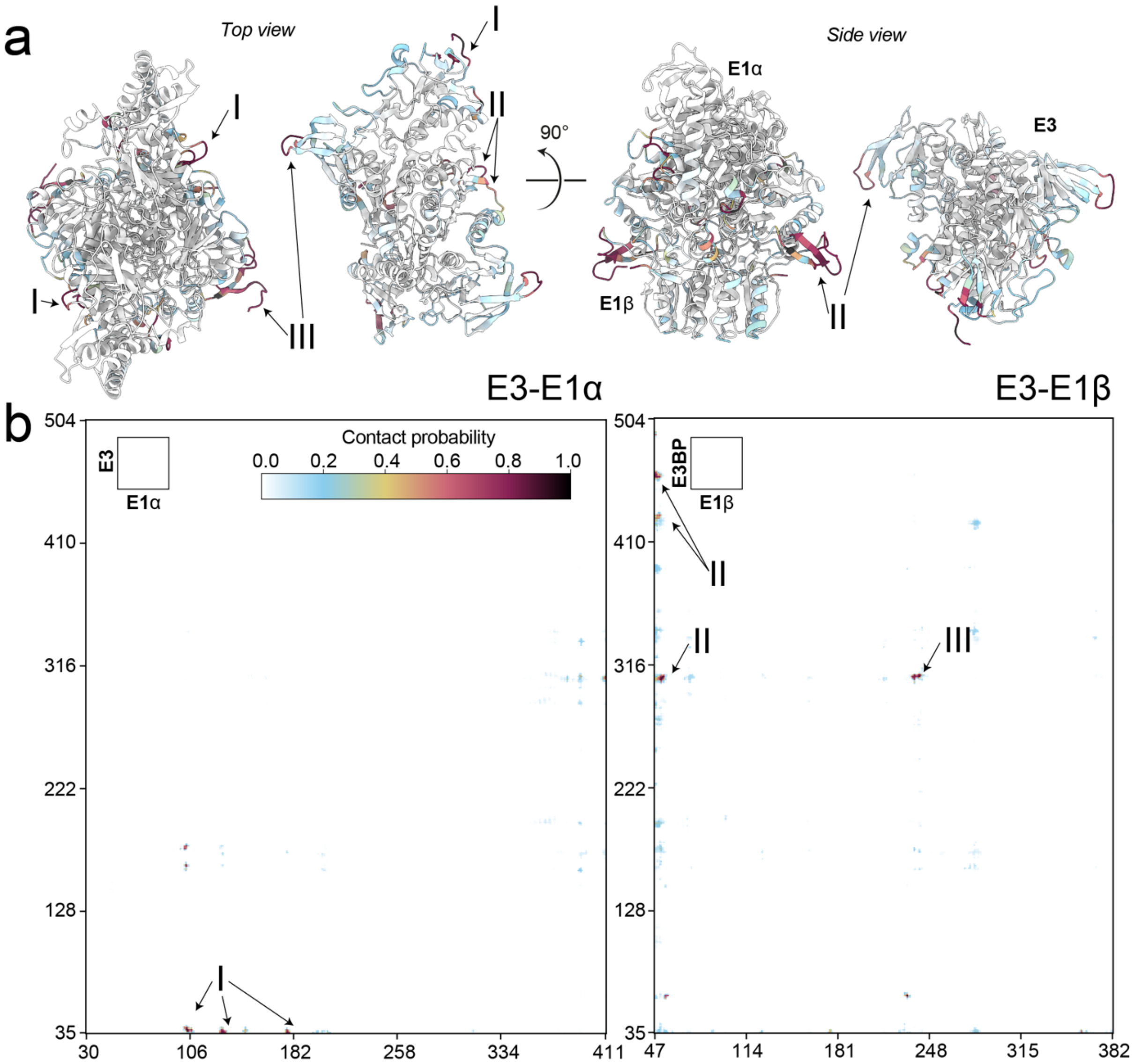


**Extended Data Figure 26: Principal contact map of the E1-E3 interaction.**
Residue-residue interaction patterns of the E1 and E3 are displayed. Structural mapping (**a**) and contact matrix (**b**) are shown. Contact frequencies are color-coded from low (light blue) to high (black), with major interaction regions labeled.

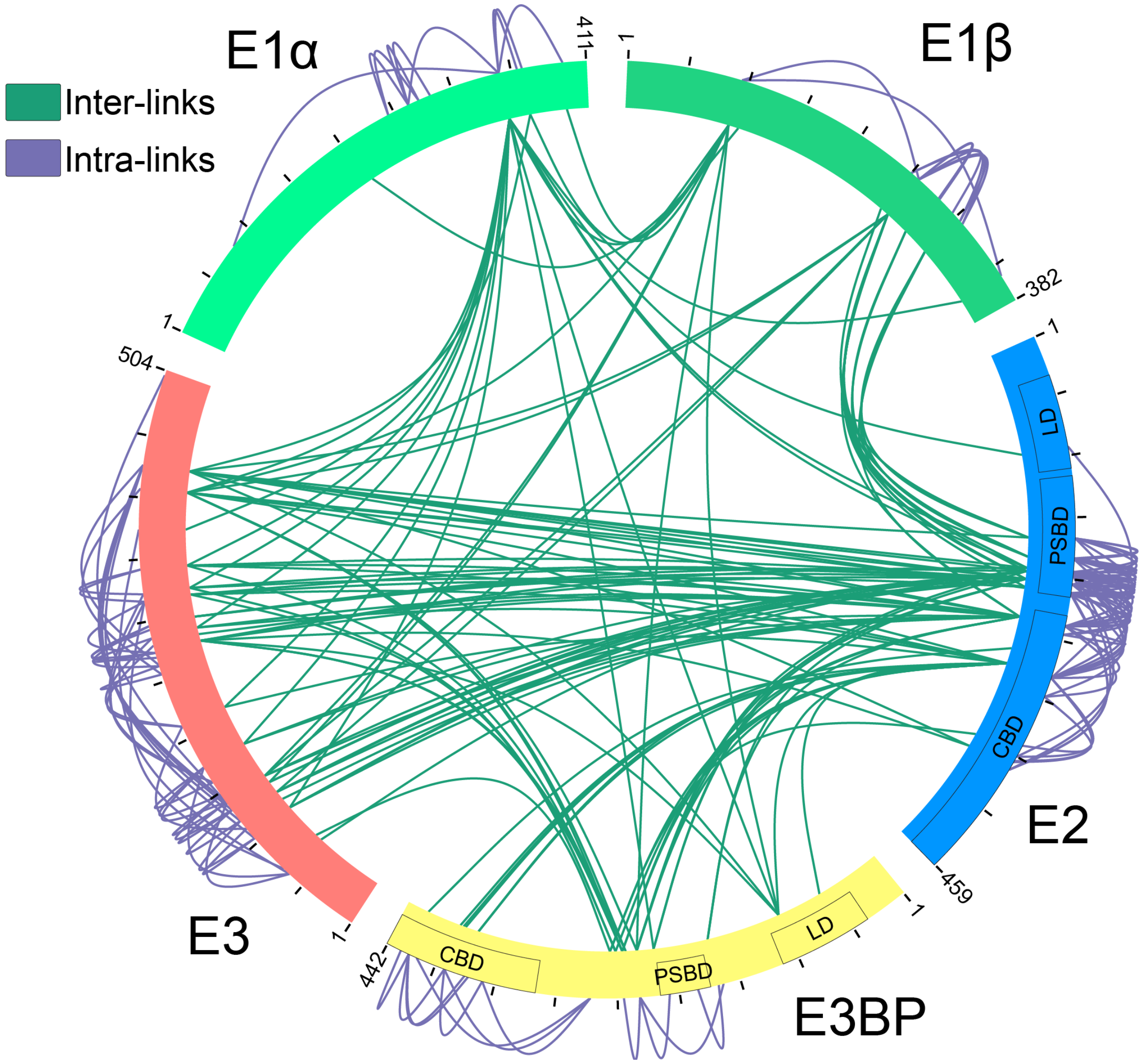


**Extended Data Figure 27: Cross-linking MS of the PDHc.**
The circular plot of the protein crosslinks among subunits of the *T. thermophila* PDHc (E1α (light green), E1β (dark green), E2 (blue), E3 (orange), and E3BP (yellow)) reveals interdomain connectivity across the catalytic network of the link reaction. Domain architecture of the E2 and E3BP is shown. Inter-links (green) and intra-links (purple) are shown. Crosslinks were visualized using Xview[8].

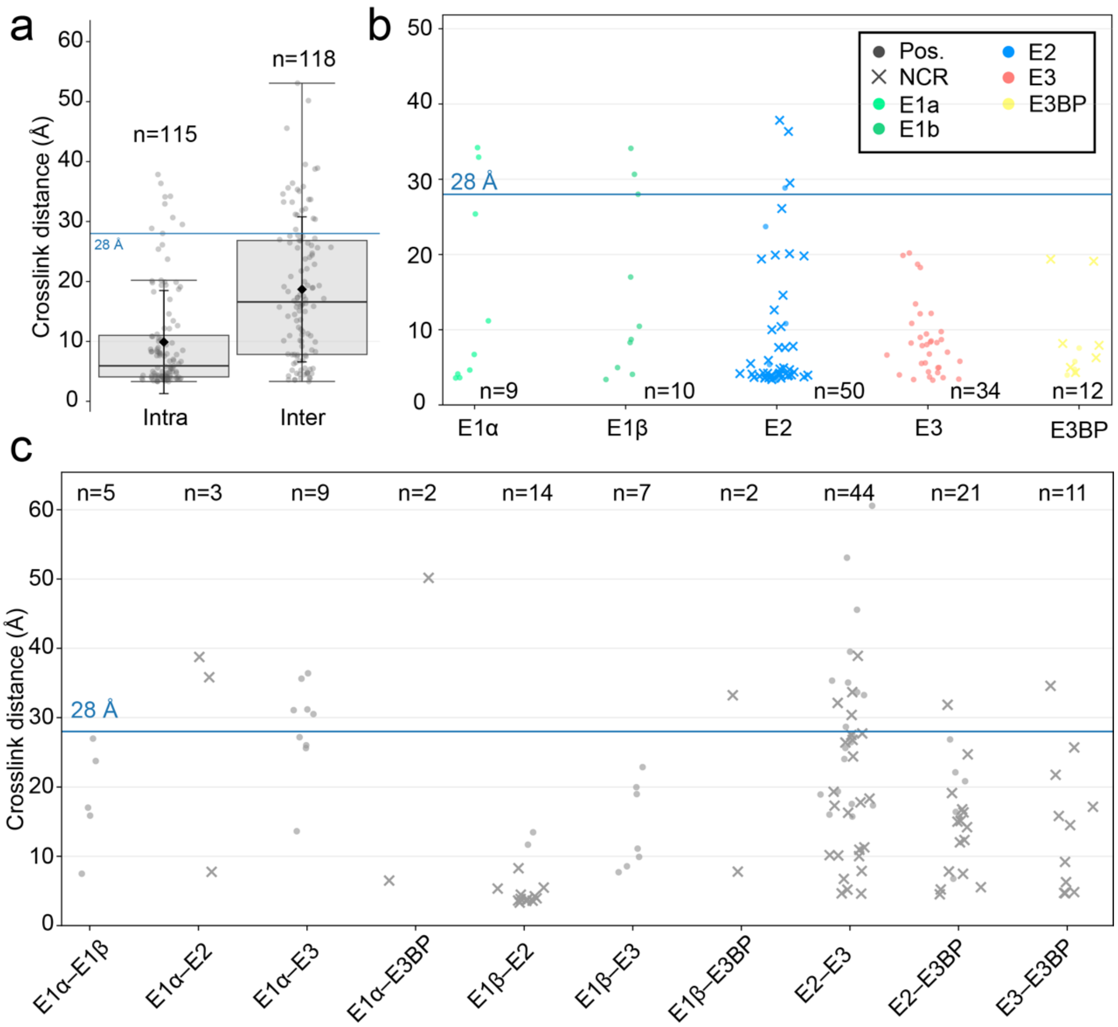


**Extended Data Figure 28: Evaluation of crosslinking distances.**
**a.** Overall satisfaction of the intra- and intercrosslinks is displayed. **b,** Intra-subunit crosslink distances are shown per subunit. Crosslinks mapping to the non-core-residing region (NCR) of the E2 CD and E3BP CB assembly are indicated (x). **c,** Inter-subunit cross-link distances grouped by interacting subunit pairs are shown.

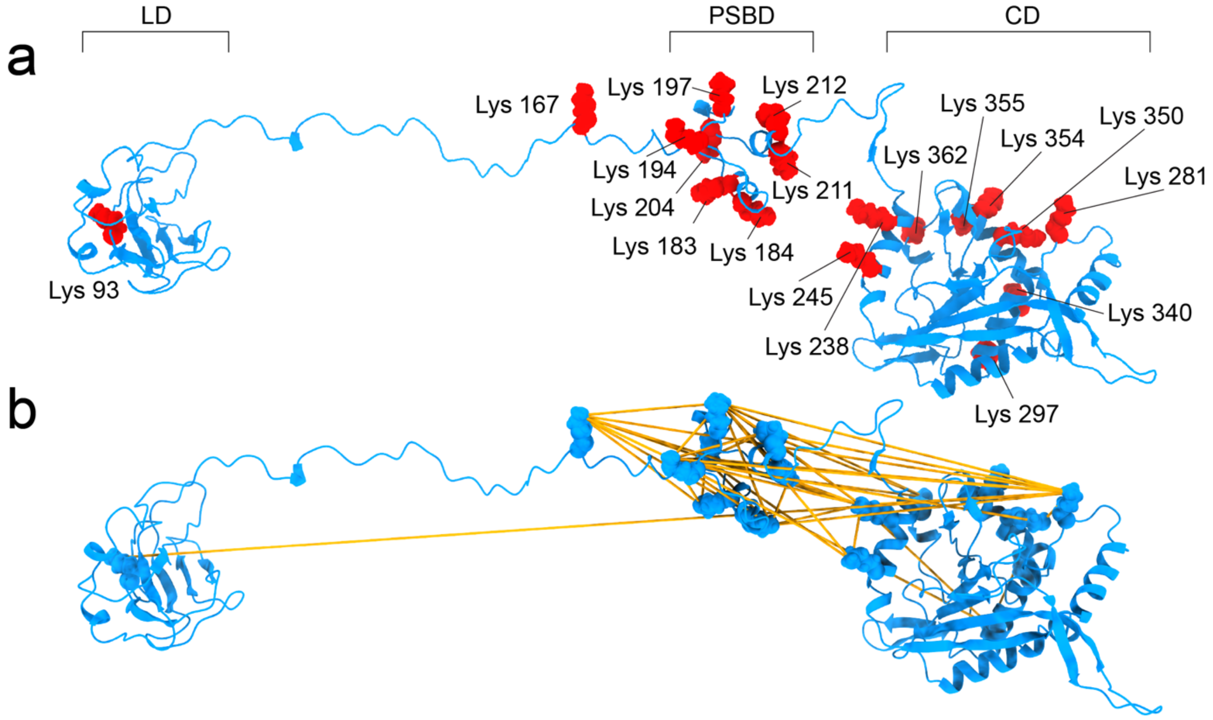


**Extended Data Figure 29: molecular interactions of the E2.**
Crosslinks mapped onto the integrative PDHc model reveal extensive connectivity among E2 domains (**a&b**).

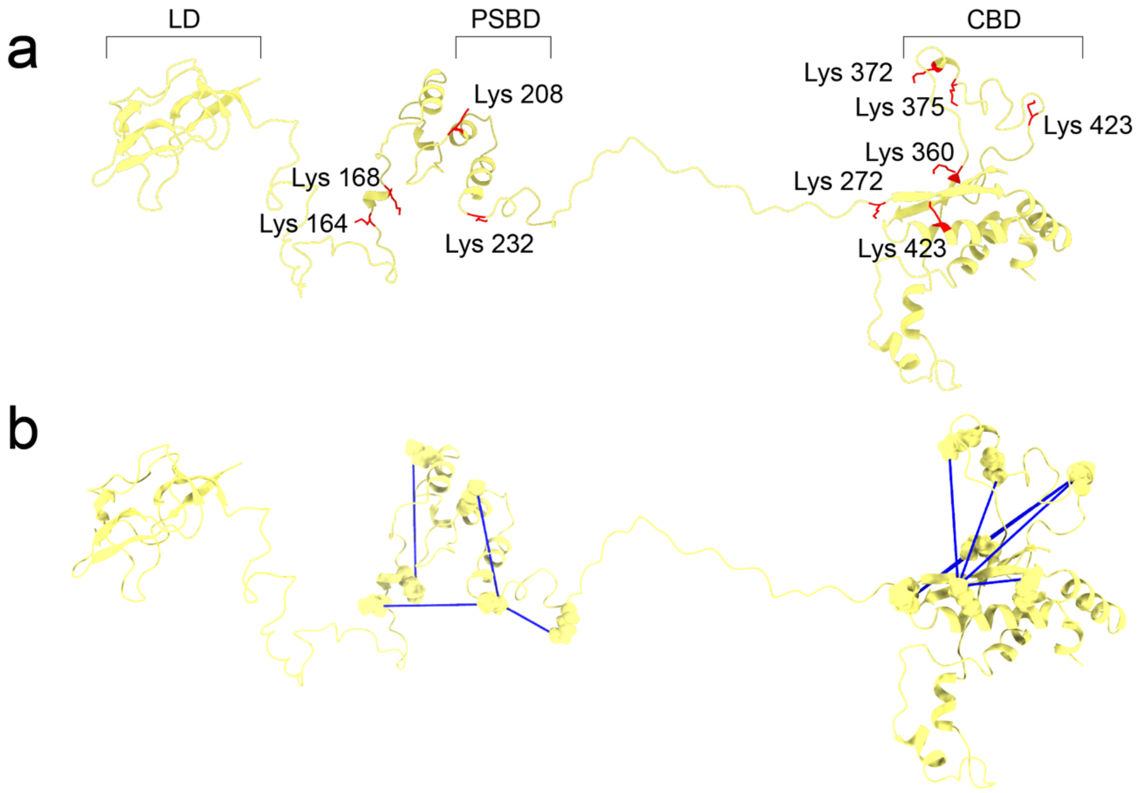


**Extended Data Figure 30: molecular interactions of the E3BP.**
The intra-crosslinks were mapped onto the model of the E3BP, revealing inter-domain connectivity within the multidomain protein (**a&b**).

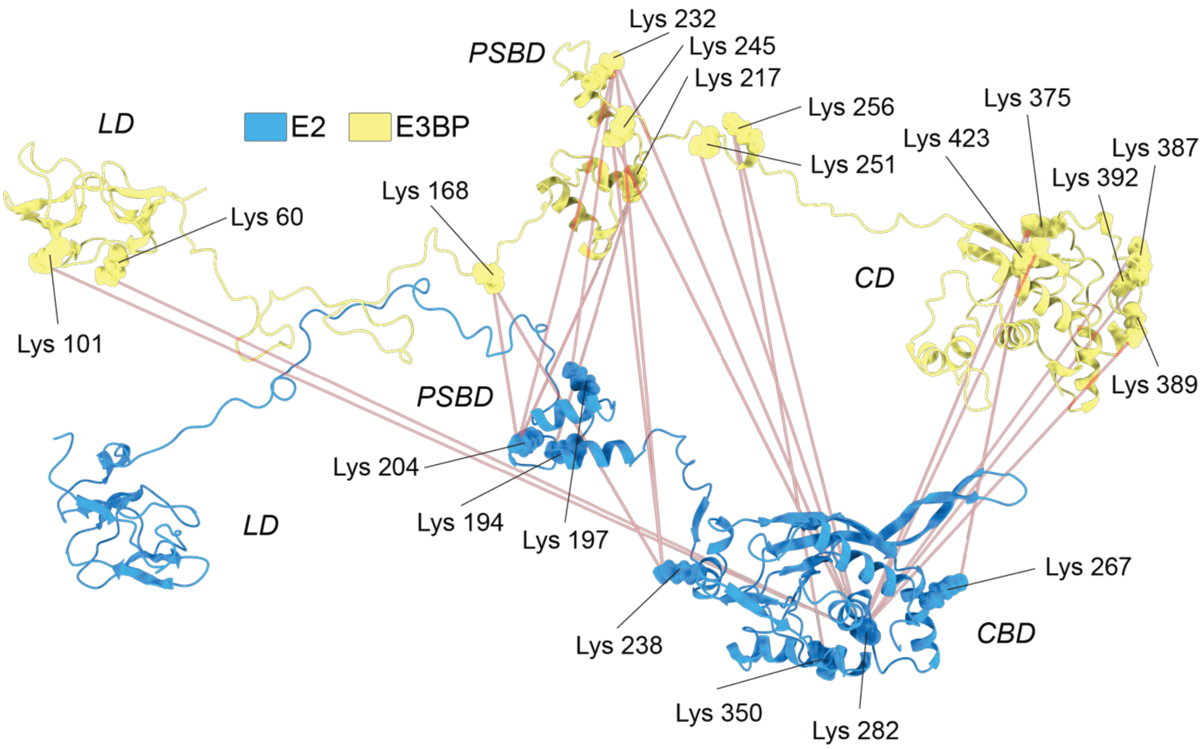


**Extended Data Figure 31: Binary interaction of the E2-E3BP.**
Inter-chain crosslink distances are shown, with each point corresponding to the distance between an individual lysine pair in the model.

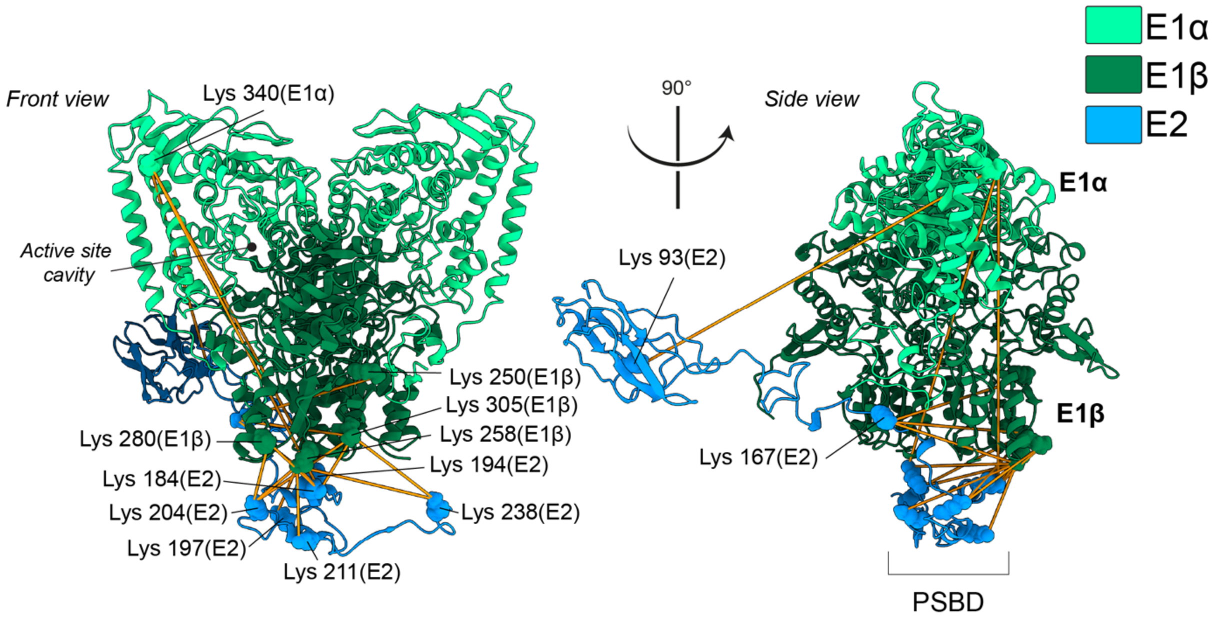


**Extended Data Figure 32: Binary interaction of the E1-E2.**
Inter-chain crosslink distances are plotted, with each point representing an individual lysine pair distance within the model.

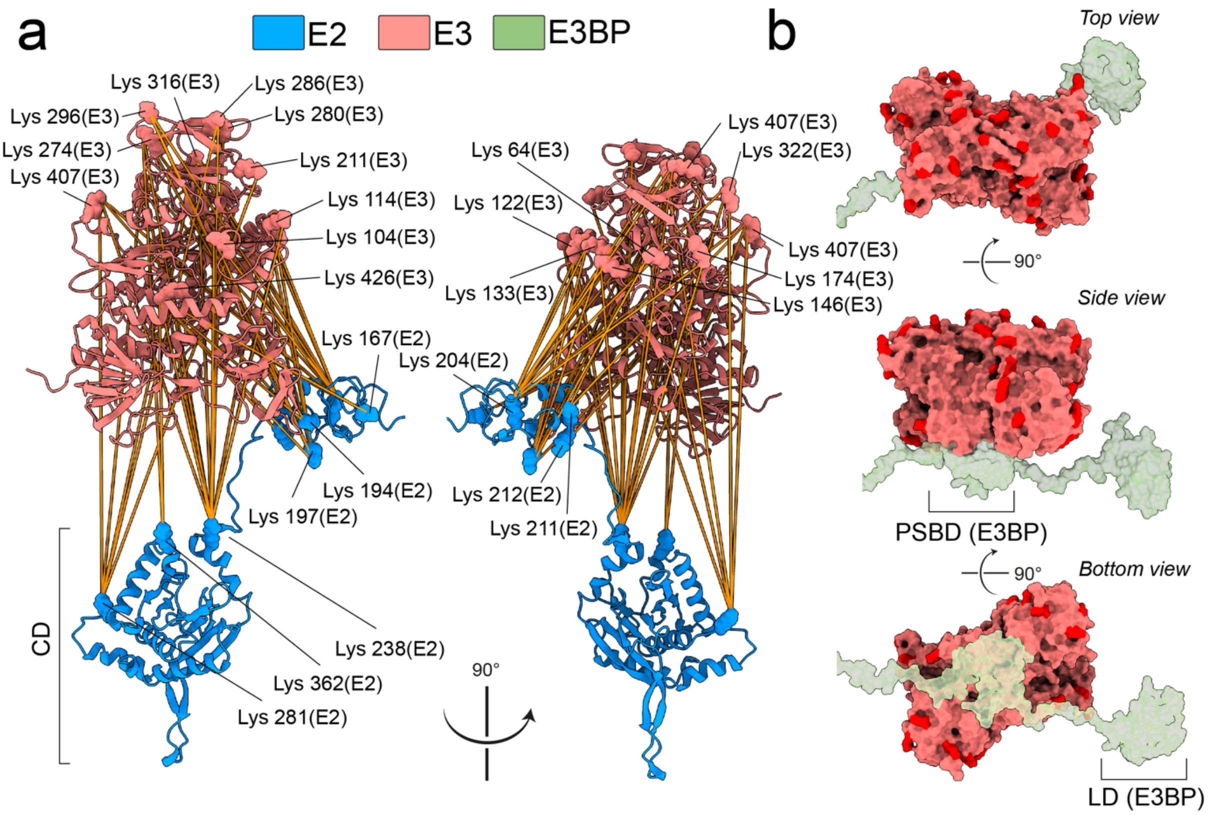


**Extended Data Figure 33: Binary interaction of the E2-E3.**
**a**, Crosslinks mapped onto the integrative PDHc model reveal extensive connectivity among E2-E3 domains. **b**, No E2–E3 crosslinks were observed at the E3BP–E3 interface, consistent with E3BP occluding this site within the assembled complex.

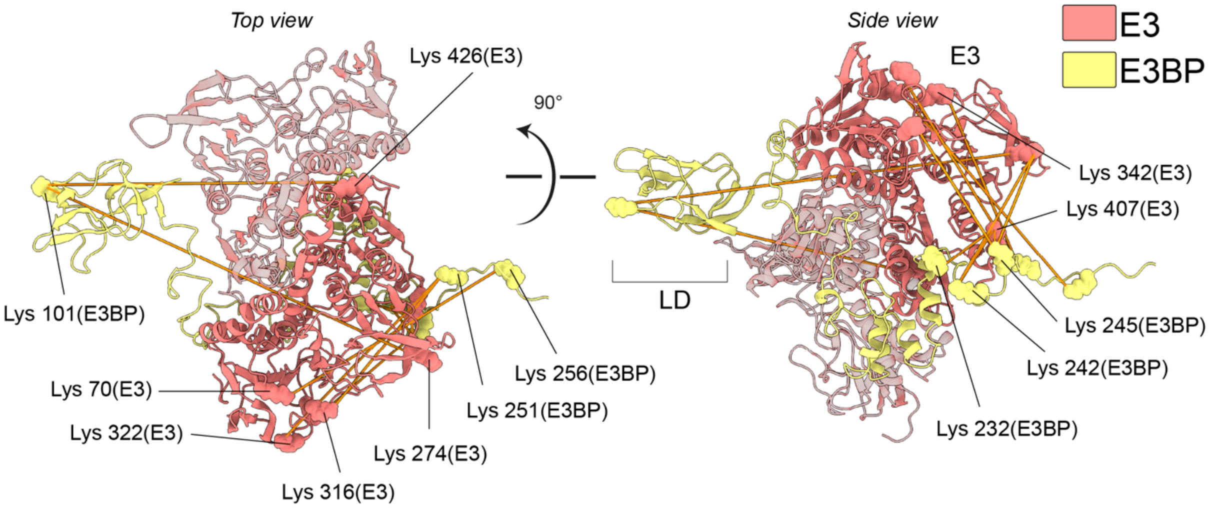


**Extended Data Figure 34: Binary interaction of the E3-E3BP.**
Crosslinks between E3 (orange) and E3BP (yellow) were mapped onto their structural models to visualize interactions. The respective residues and their interactions are marked.

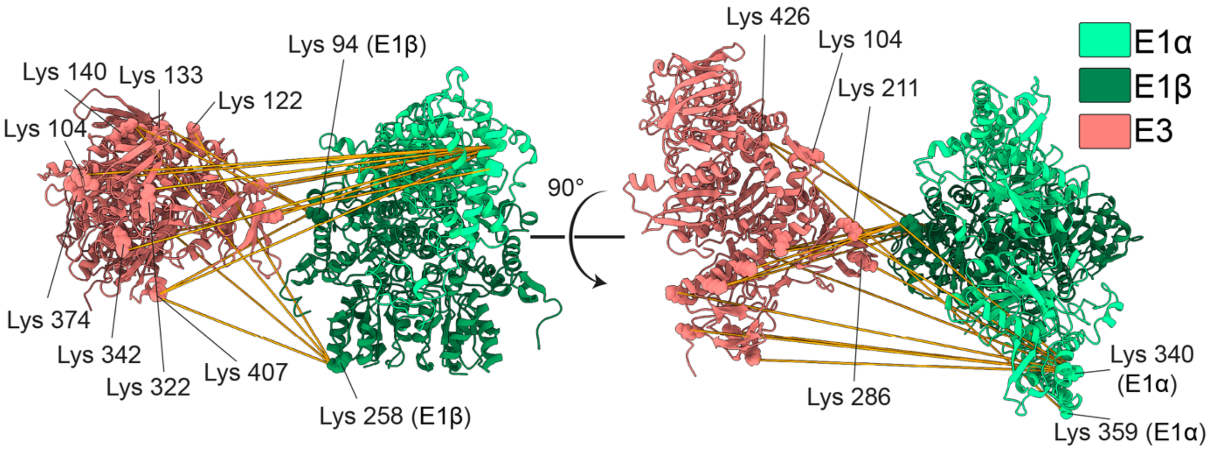


**Extended Data Figure 35: Binary interaction of the E1-E3.**
Inter-crosslinks between the E1 and E3 domains of the PDHc are shown, highlighting the individual lysine pairs. Despite the absence of a direct catalytic interaction, E1 and E3 share several interaction sites due to tight packing within the redox shell.

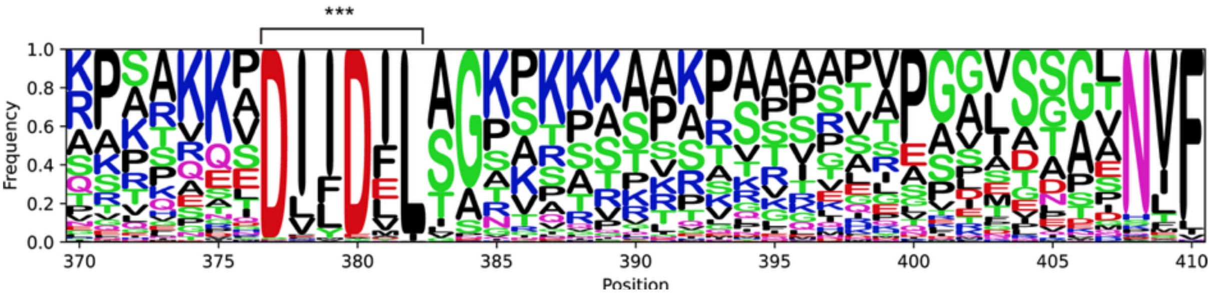


**Extended Data Figure 36: Conservation of the E3BP.**
To identify the conserved regions responsible for mediating E2-E3BP association, the sequence conservation of fungal proteins was examined using the MSA implementation of AF2 via ConservedFold[4]. The unstructured region between residues 360 and 407 features a conserved DIIDIL motif that is highly retained across the fungal E3BP sequences.

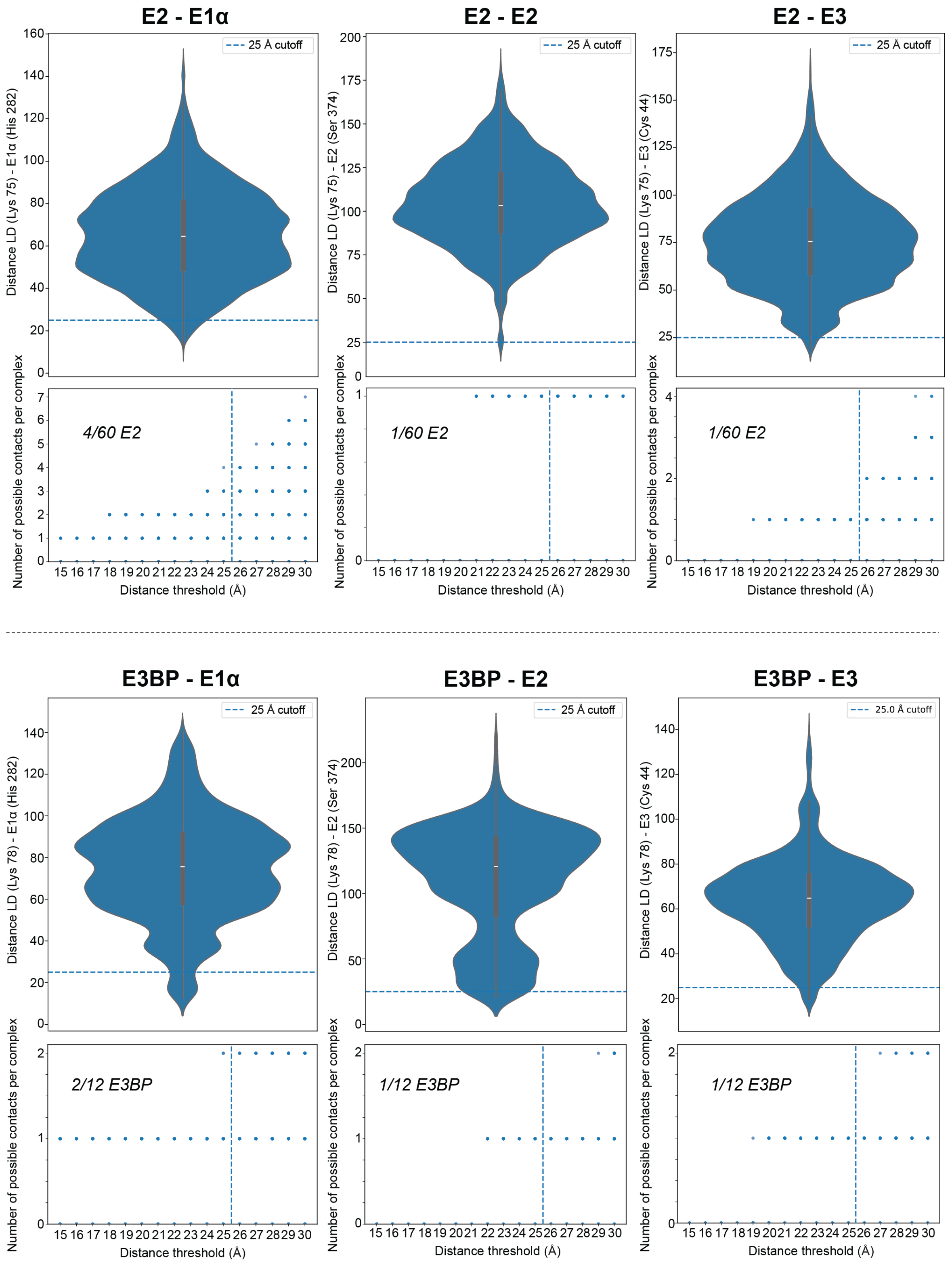


**Extended Data Figure 37: Simulated proximity of LD to the active sites of E1, E2, and E3.** For each indicated binary interaction of the LD (E2&E3BP), the upper panel shows the distribution of distances between the lipoamide-bearing lysine and the selected catalytic-site residue across the simulated ensemble, per protein chain and ensemble structure (Cα–Cα distances). The dashed horizontal line marks the 25 Å contact cutoff (chosen to account for the bridging lipoamide group and to account for flexibility). The lower panel benchmarks the number of satisfied contacts observed per complex in each simulation frame, for different distance thresholds.

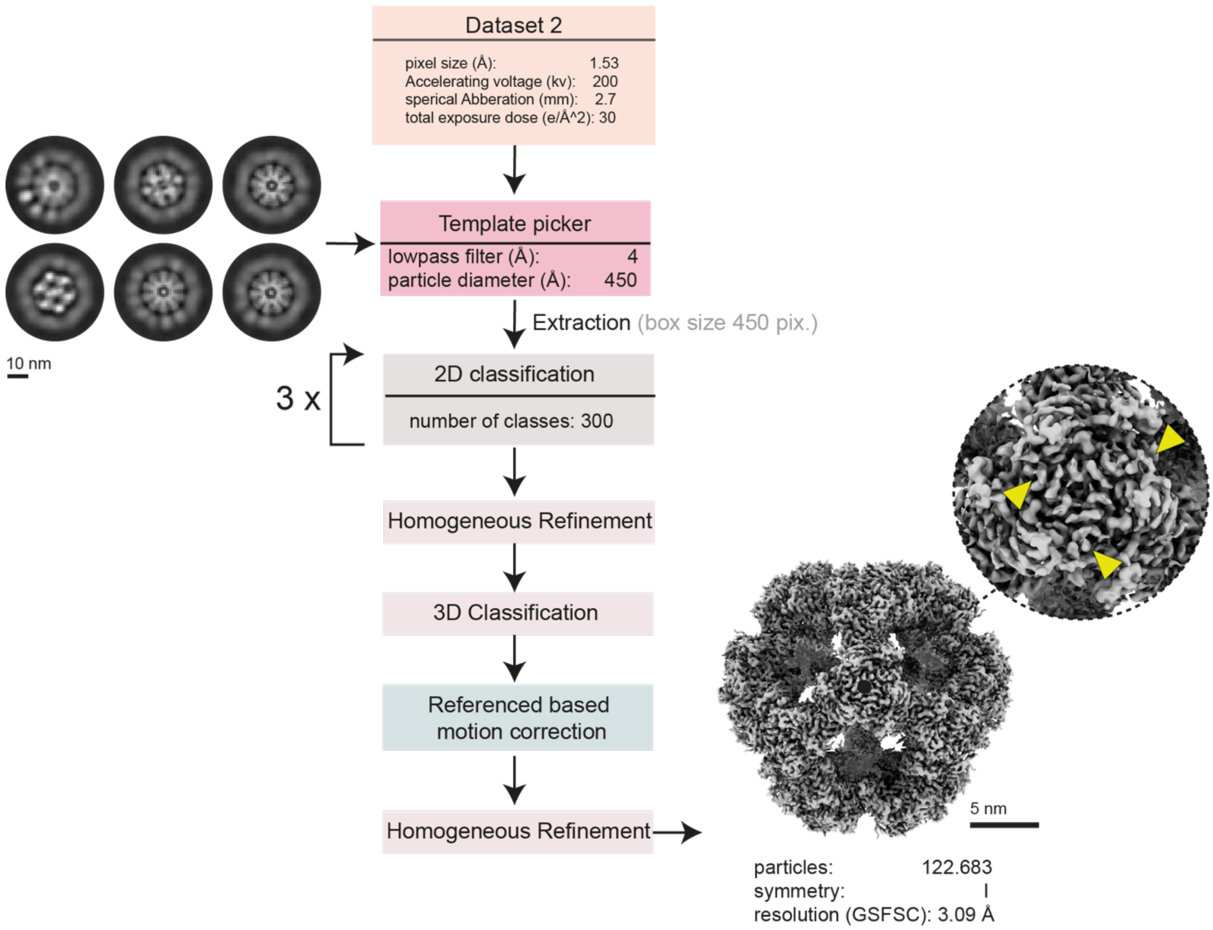


**Extended Data Figure 38: Reconstruction of substrate-bound E2 core.**
Schematic representation of the image analysis pipeline for substrate-bound E2 CD.

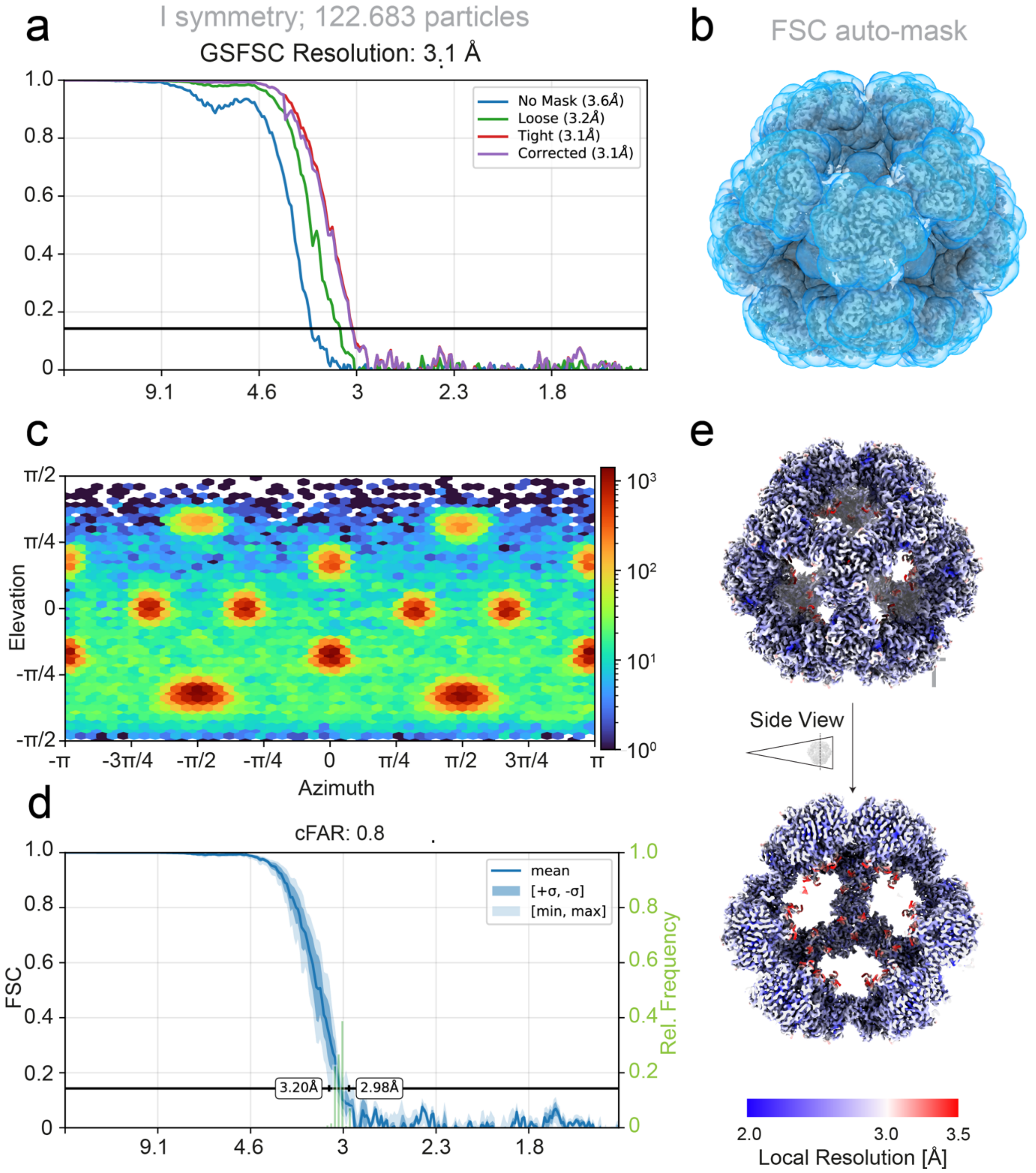


**Extended Data Figure 39: Quality metrics of the substrate-bound E2 core.**
**a,** The substrate-bound E2 CD of the PDHc was reconstructed at 3.1 Å (FSC = 0.143, I symmetry) using 122683 particles. **b**, The mask used for structure refinement is displayed. **c-d**, The overall view coverage and the directional FSC (3D FSC) [2] are shown. **e,** The calculated local resolution estimation shows a distribution between 2 and 3.5 Å (contour level σ 0.2).

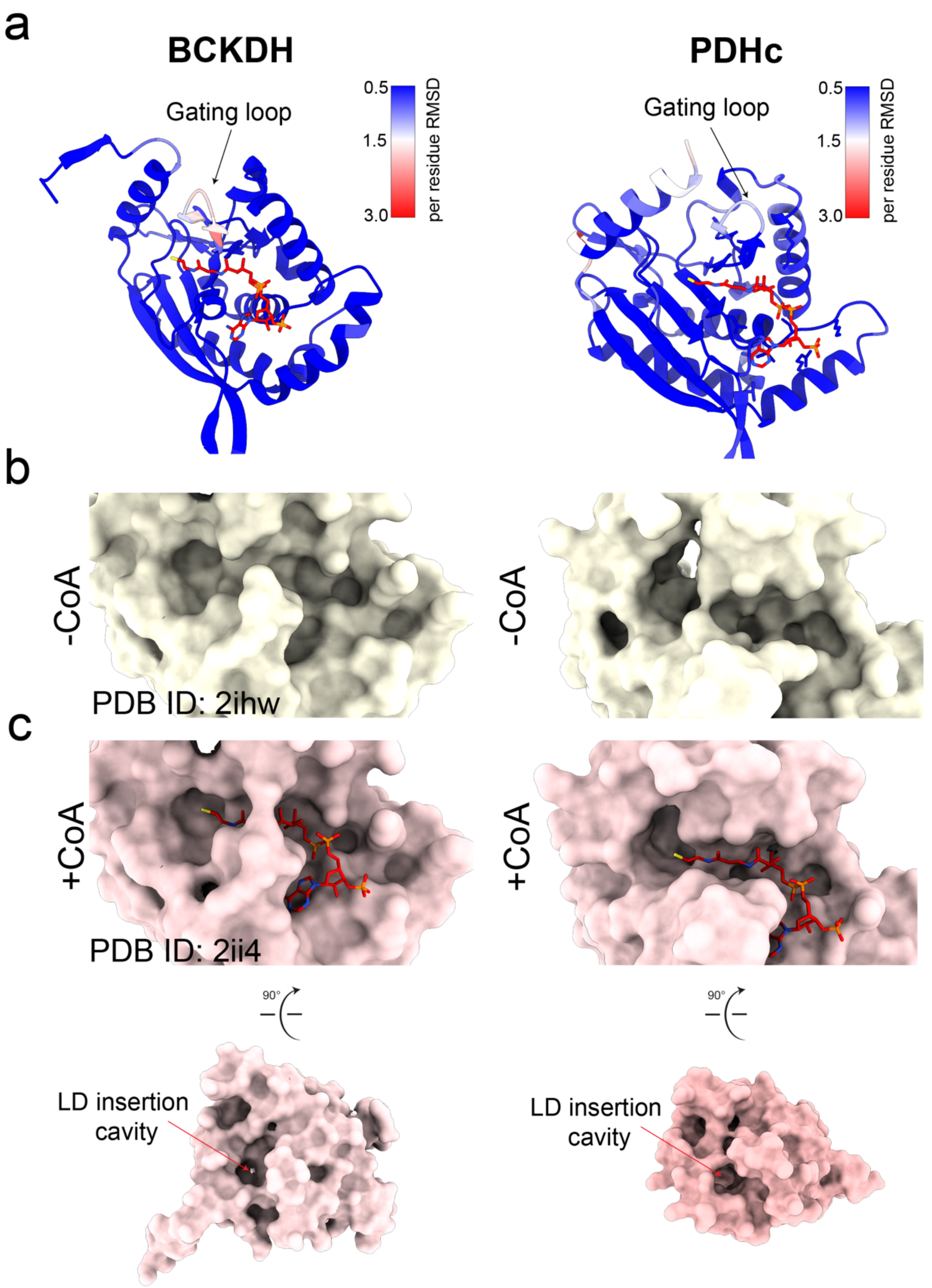


**Extended Data Figure 40: Comparison of substrate-induced conformational changes in the BCKDH complex and the PDHc.**

The monomeric E2 CD of the BCKDH (left) and the PDHc (right) show overall high structural conservation (**a-c**), particularly in the positioning and binding configuration of CoA[9]. To examine conformational rearrangements associated with substrate binding and transacetylation, the ground-state and CoA-bound structures were compared (BCKDH: PDB IDs 2IHW and 2II4). In BCKDH, CoA binding induces a 2 Å displacement of the active-site gating loop (**a**), which has been proposed to facilitate opening of a transient pore for LD insertion. Surface representations of the apo and CoA-bound conformations illustrate slight expansion of the LD insertion cavity in BCKDH, whereas the corresponding region in PDHc remains largely unaltered upon CoA binding (**b&c**).

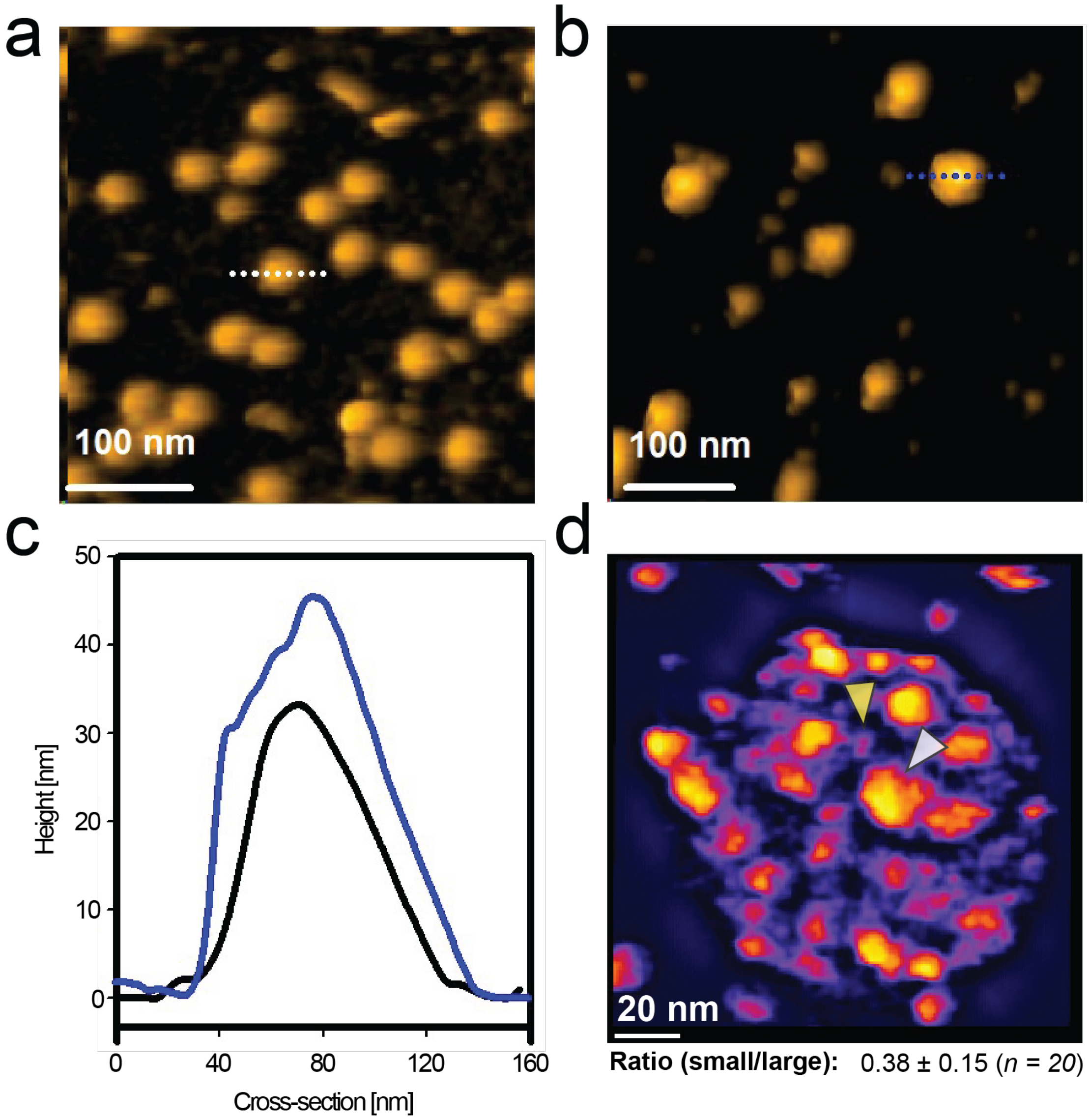


**Extended Data Figure 41: HS-AFM resolves the single-molecule architecture of PDHc particles in the ground state and saturated state.** **a, b,** HS-AFM images of representative PDHc particles adsorbed on mica in ground state (in **a**) and in saturated state (in **b**). **c,** Representative particle signatures and cross-sectional height profiles in the ground state (GS) and under substrate-saturating conditions (Satur.) show an increase in apparent height upon substrate addition. **d,** Representative HS-AFM image of a single PDHc particle showing distinct peripheral features with different structural signatures. The ratio of smaller (LD-related, yellow arrow) to larger features (E1/E3, white arrow) agrees with the expected stoichiometric relationship between the 72 LDs and 32 peripheral subunits of endogenous PDHc (~0.4).

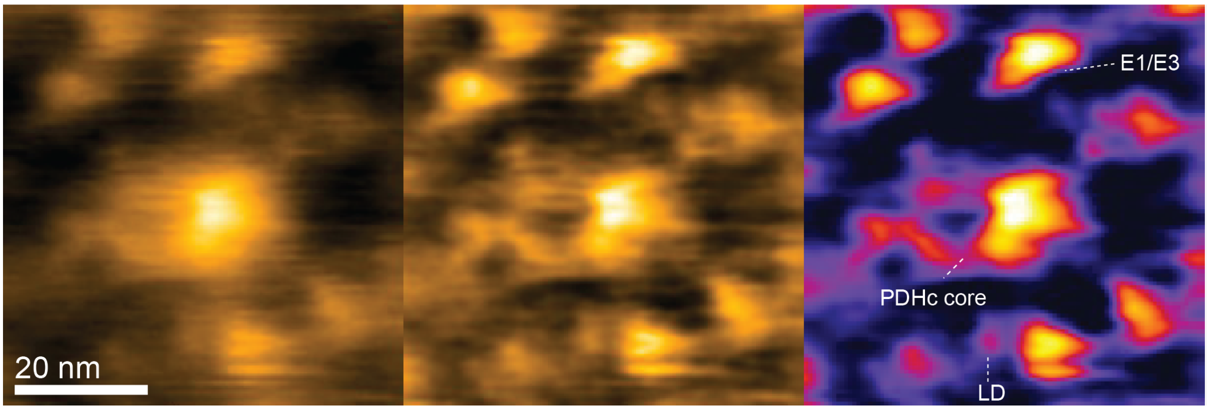


**Extended Data Figure 42: Force-dependent AFM imaging resolves core and peripheral regions of PDHc.** Unfiltered (left) and band-pass filtered (middle, right) AFM images show distinction between PDHc core- and peripheral domains. The central feature corresponds to the PDHc E2/E3BP core, whereas outer signatures can be related to peripheral E1/E3 assemblies and LD-associated signals.

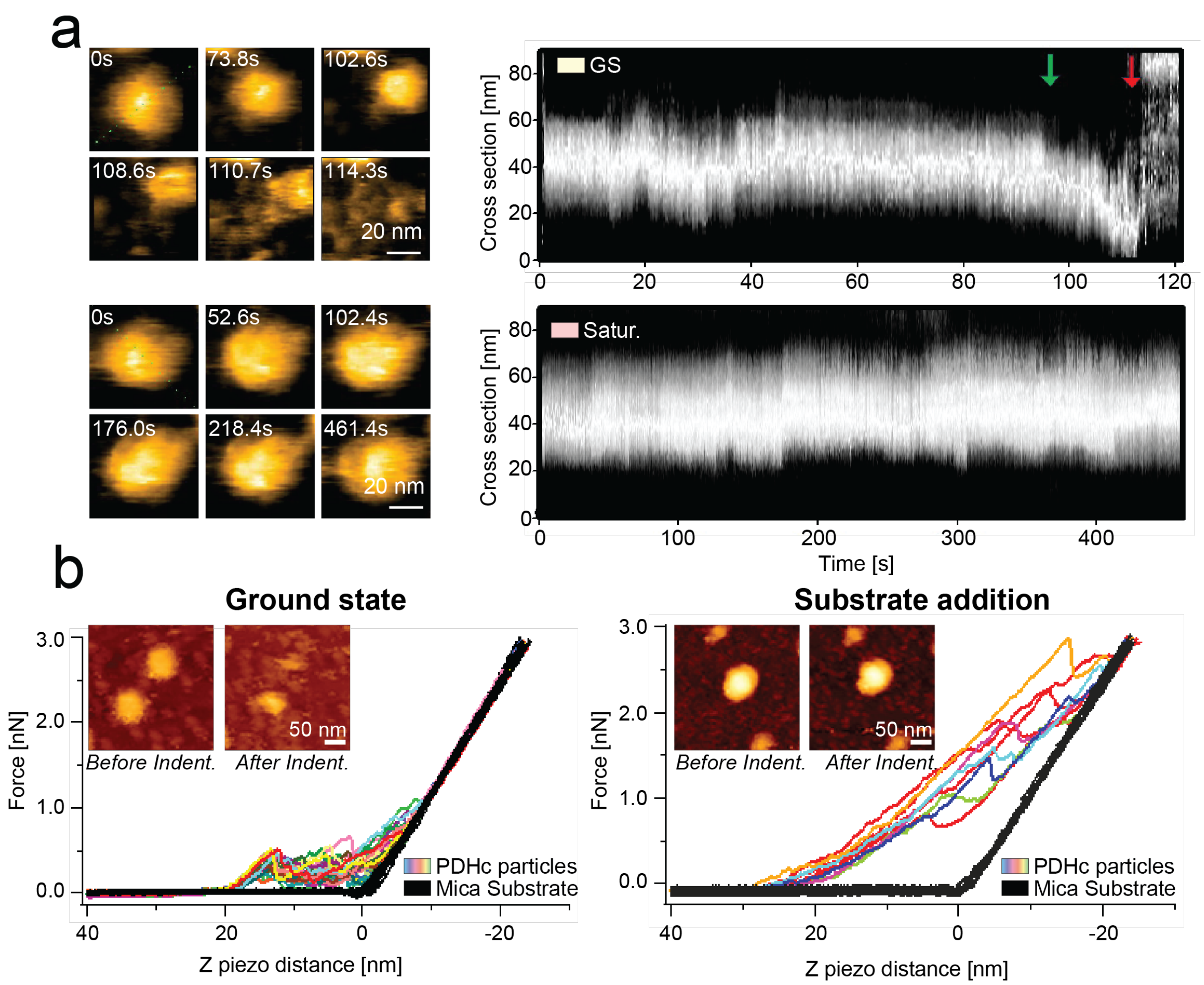


**Extended Data Figure 43: Substrate-dependent mechanical stabilization of PDHc**. **a,** Mechanical fatigue experiments support enhanced particle stability under substrate-saturating conditions. Representative time series and corresponding kymographs show greater structural persistence after substrate addition compared to the GS. **b,** Nanoindentation measurements of assembled PDHc in the GS and under substrate-saturating conditions reveal greater resistance to indentation after substrate addition, consistent with increased stiffness and mechanical stability. Indentation on mica is shown as a control. Insets show representative particles before and after indentation.

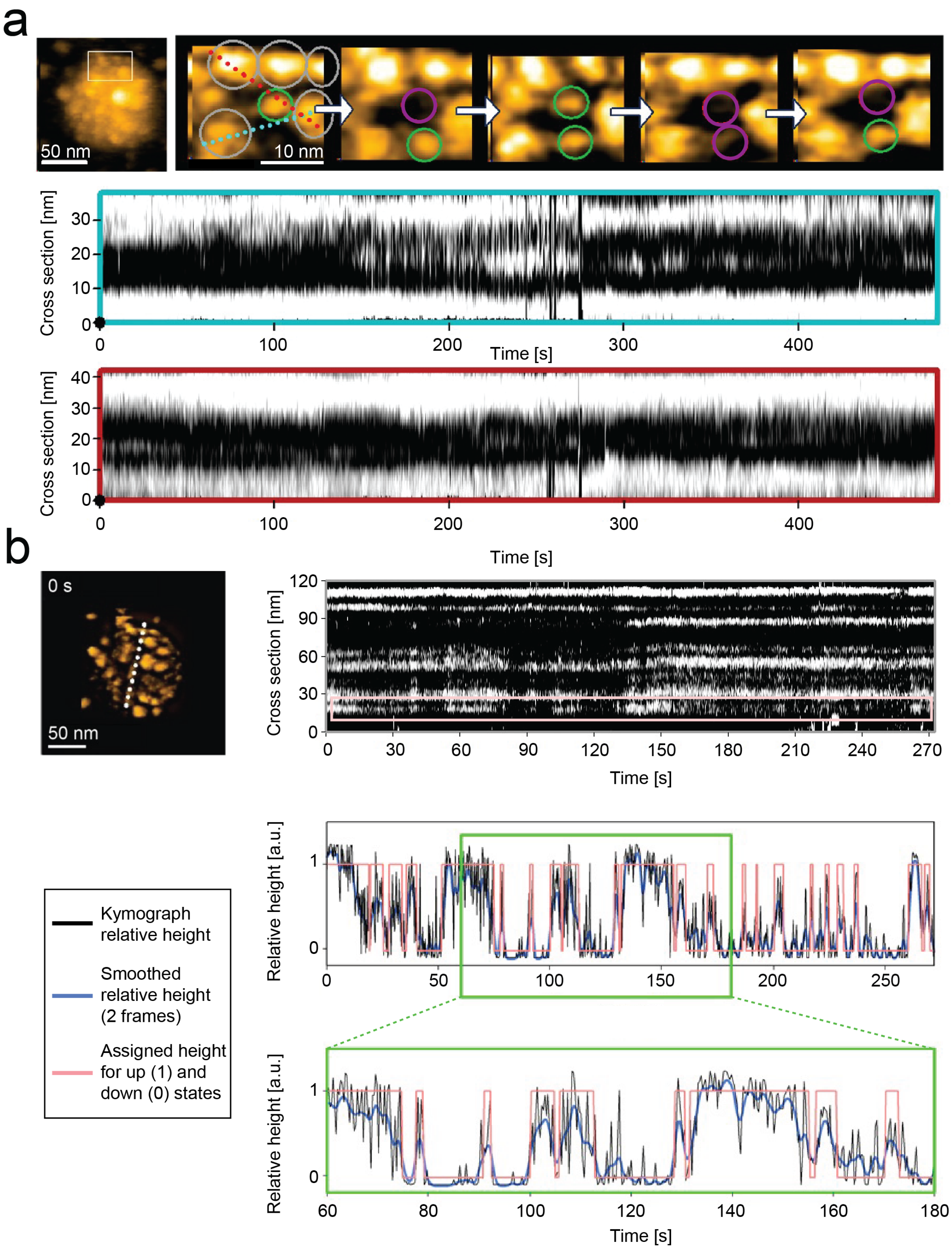


**Extended Data Figure 44: Visualization of peripheral domain mobility.**
**a,** Representative HS-AFM image of a single PDHc particle and enlarged series of image examples showing dynamic rearrangements of peripheral LD-related signatures. Kymographs along the indicated scan lines reveal temporally resolved transitions between distinct peripheral and core-bound states. Grey circles indicate the stable larger domain, followed by smaller domains in the down (purple circle), or up state (green circle). **b,** Single frame HS-AFM image using band pass filter (top left). (Top right) Kymographs showing the height fluctuations along the white dashed line of the HS-AFM image reveal localized domain rearrangements over time. (right middle)

Representative relative height profiles over time of one smaller domain marked by a magenta rectangle in the above kymograph; (right bottom) a zoomed-in relative height graph showing different curve fittings to extract the relative state of the LD domains. Black line representing direct cross-section from the kymograph; blue line shows smooth cross-section over two frames; the red line represents the assigned up (1) and down (0) states of the domain of interest.

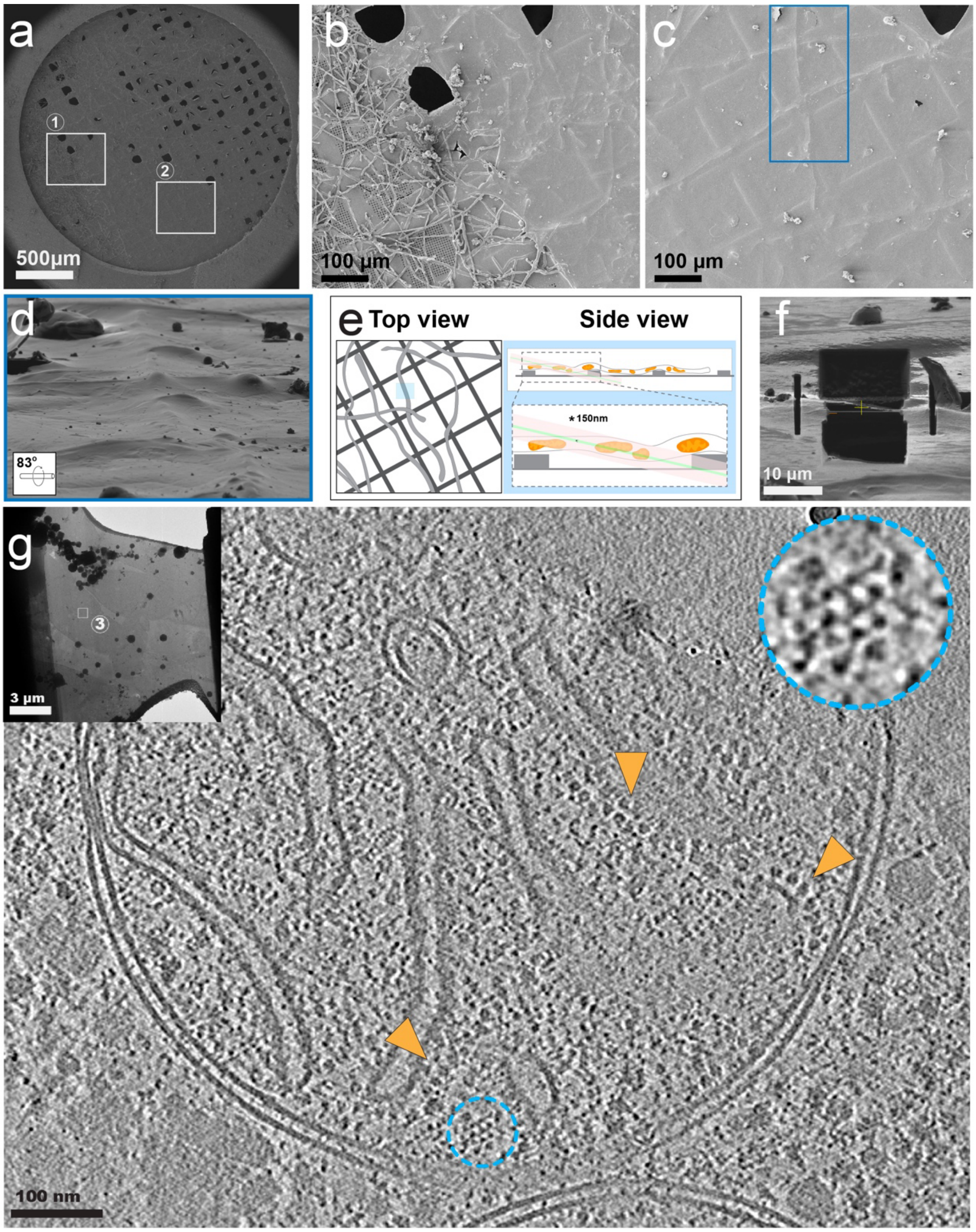


**Extended Data Figure 45: Cellular lamella from *T. thermophila*.**

**a,** Hyphae of the thermophilic fungus *T. thermophila* were cultured and vitrified directly on the TEM grid to enable imaging of the intact cellular environment. An exemplary SEM image of the prepared grid containing the fungal specimen is shown, highlighting regions of interest selected for targeted FIB milling (**b&c**). **d,** Regions of interest where hyphae are uniformly embedded in vitreous ice were selected for lamella preparation. **e,** Prior characterization demonstrated that the fungal hyphae are densely populated with mitochondria [10], making it possible to perform unspecific milling with a high likelihood of targeting mitochondria. **f,** Selected sites were FIB-milled to an approx. thickness of 100-200 nm, producing electron-transparent lamellae, which were then transferred to the TEM for cryo-ET tilt-series acquisition. **g,** The reconstructed, denoised tomogram (slice shown) displays a structurally preserved mitochondrion in its cellular environment. The location of an exemplary PDHc complex is highlighted (cyan), displaying the clear icosahedral symmetry of the core as well as surrounding peripheral subunits. Complex V of the respiratory chain is marked in orange.

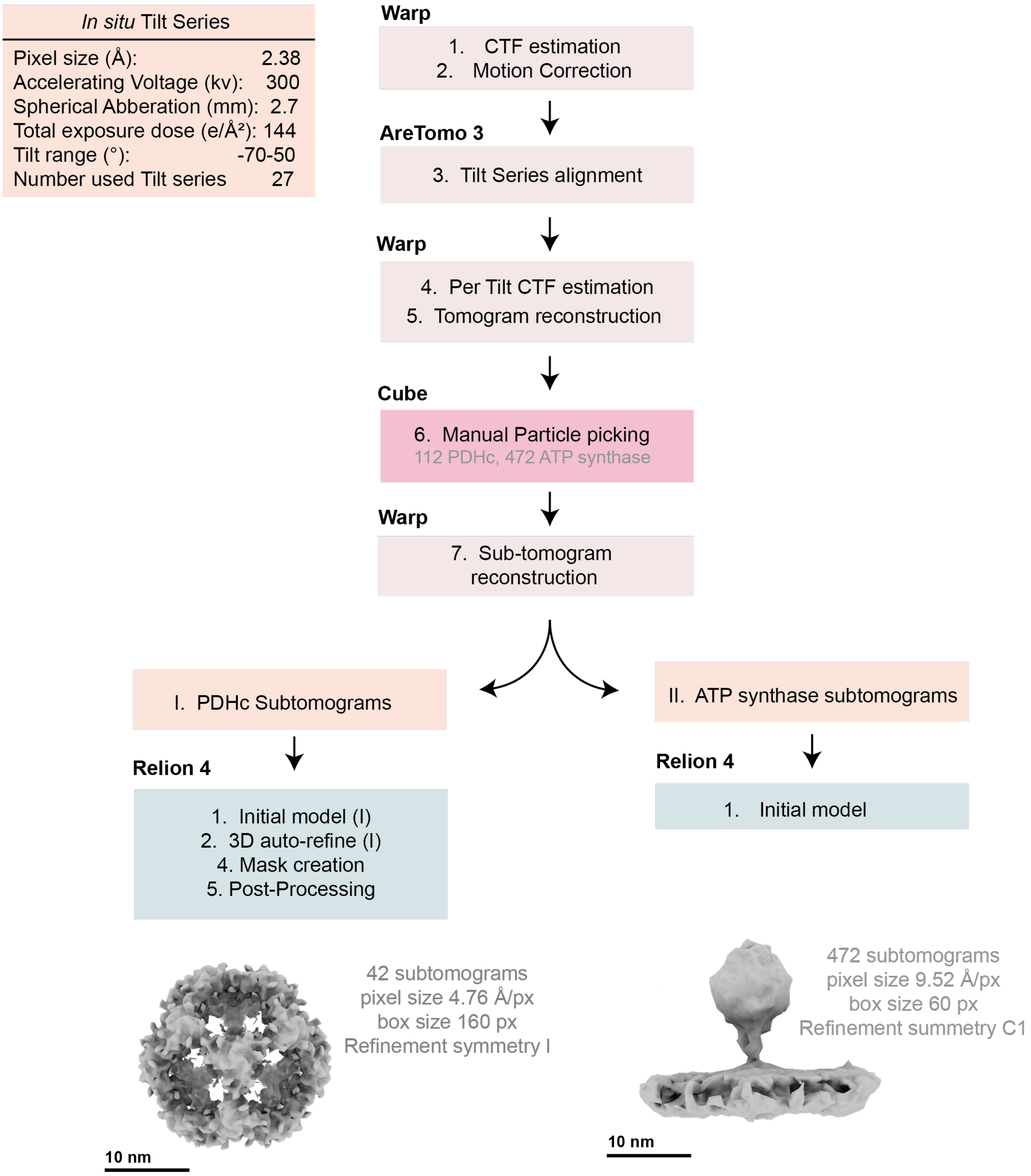


**Extended Data Figure 46: Schematic representation of the processing pipeline used for the reconstruction of the *in situ* PDHc and ATP synthase.**

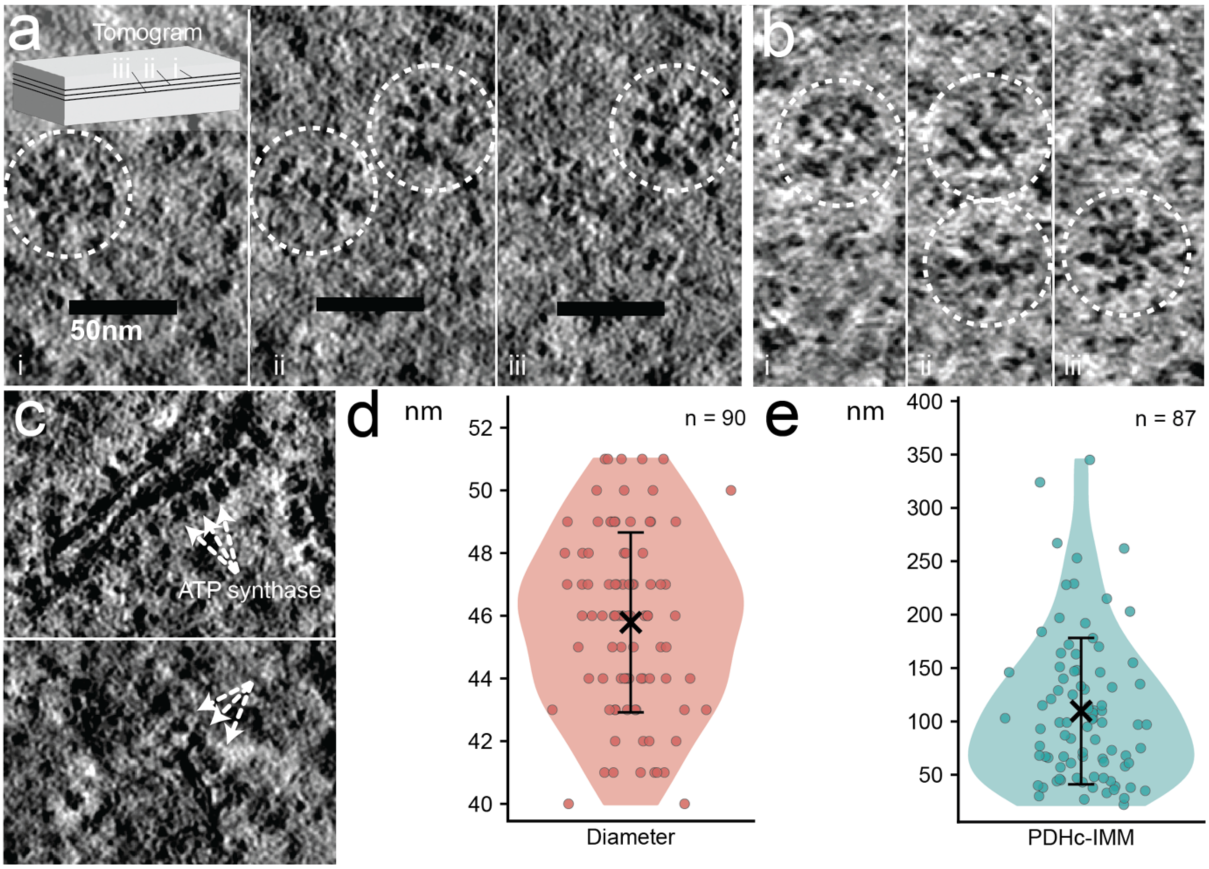


**Extended Data Figure 47: *In situ* analysis of the PDHc.**
**a,** Consecutive tomogram slices of mitochondrial lamellae showing two PDHc molecules in close proximity (i-iii). **b,** Pairs of contacting PDHc molecules were identified on multiple occasions throughout the tomogram dataset. **c,** Cristae bearing ATP synthase are shown. **d,** PDHc diameters were measured directly from *in situ* tomograms manually[11]. **e,** The distances from each PDHc to the nearest inner mitochondrial membrane (IMM) were measured manually.

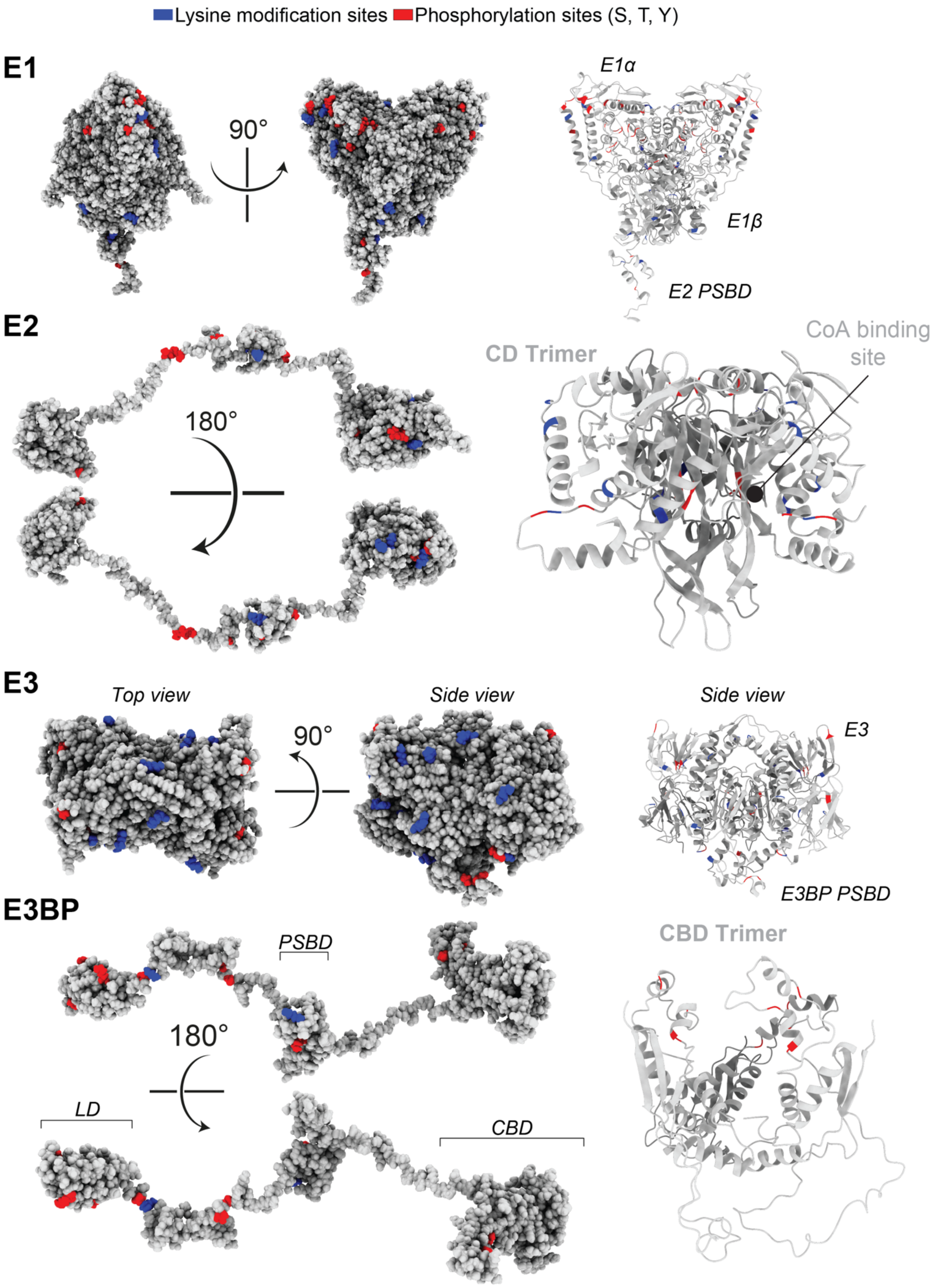


**Extended Data Figure 48: Structural mapping of post-translational modifications on PDHc components.** Surface and cartoon representations of E1, E2, E3, and E3BP with experimentally reported PTM sites from yeast and human proteins mapped onto the integrative model. Phosphorylation sites (Ser/Thr/Tyr) are shown in red and lysine-directed modifications (acetylation, ubiquitination, succinylation, and sumoylation) in blue.

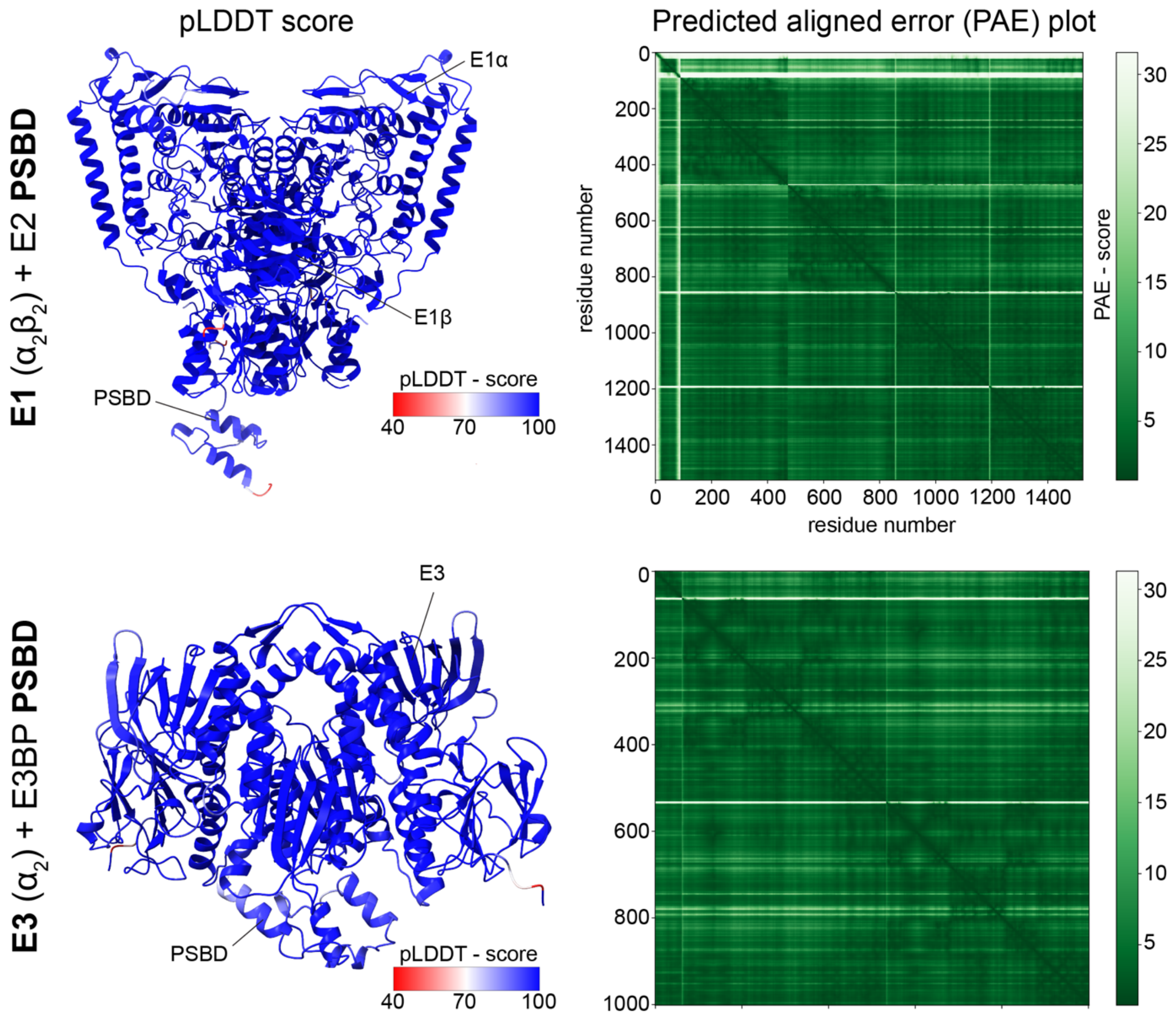


**Extended Data Figure 49: Prediction of Peripheral Subunits.**
AF2 predictions of the PSBD-bound E1 and E3 complexes with their corresponding pLDDT scores and PAE plots are shown.

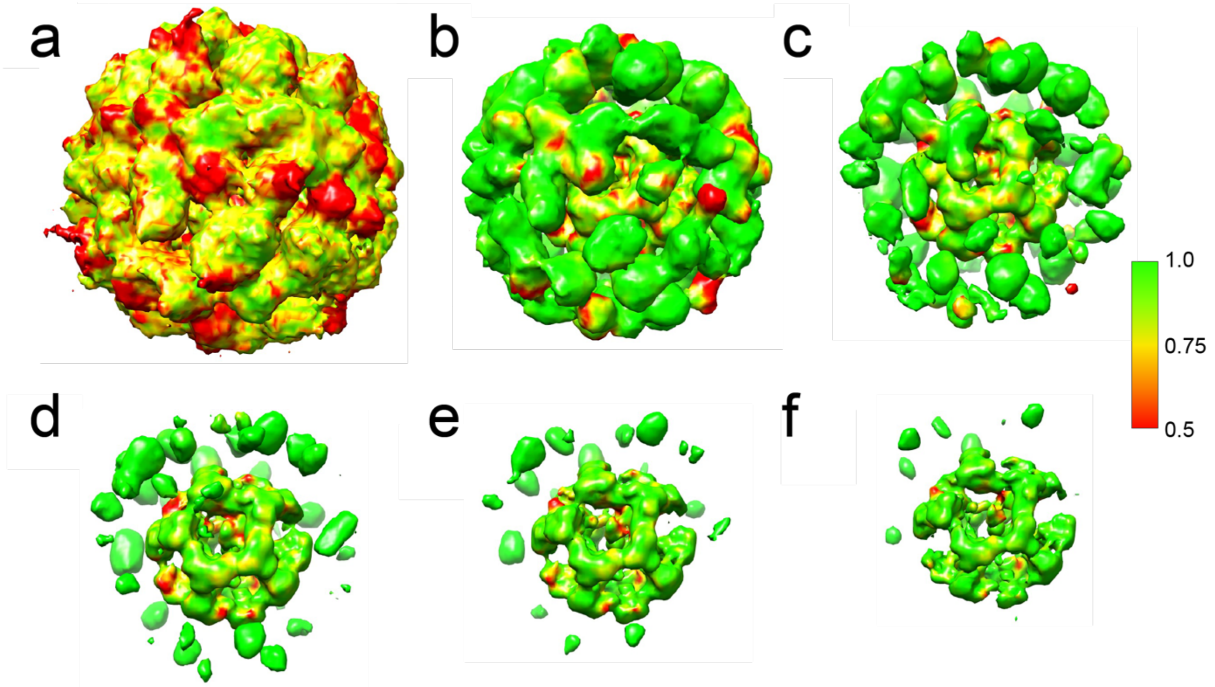


**Extended Data Figure 50: Local cross-correlation between experimental and MEMMI-based simulation.**

Local correlation between the 6 Å trimmed experimental asymmetric reconstruction of the PDHc of the ensemble-based map, projected on the former map, as a function of increasing strength of density (decreasing electron density thresholds): (**a**) 0σ, (**b**) 1σ, (**c**) 2σ, (**d**) 3σ, (**e**) 4σ, (**f**) 5σ. The global cross-correlation corresponds to 0.752.

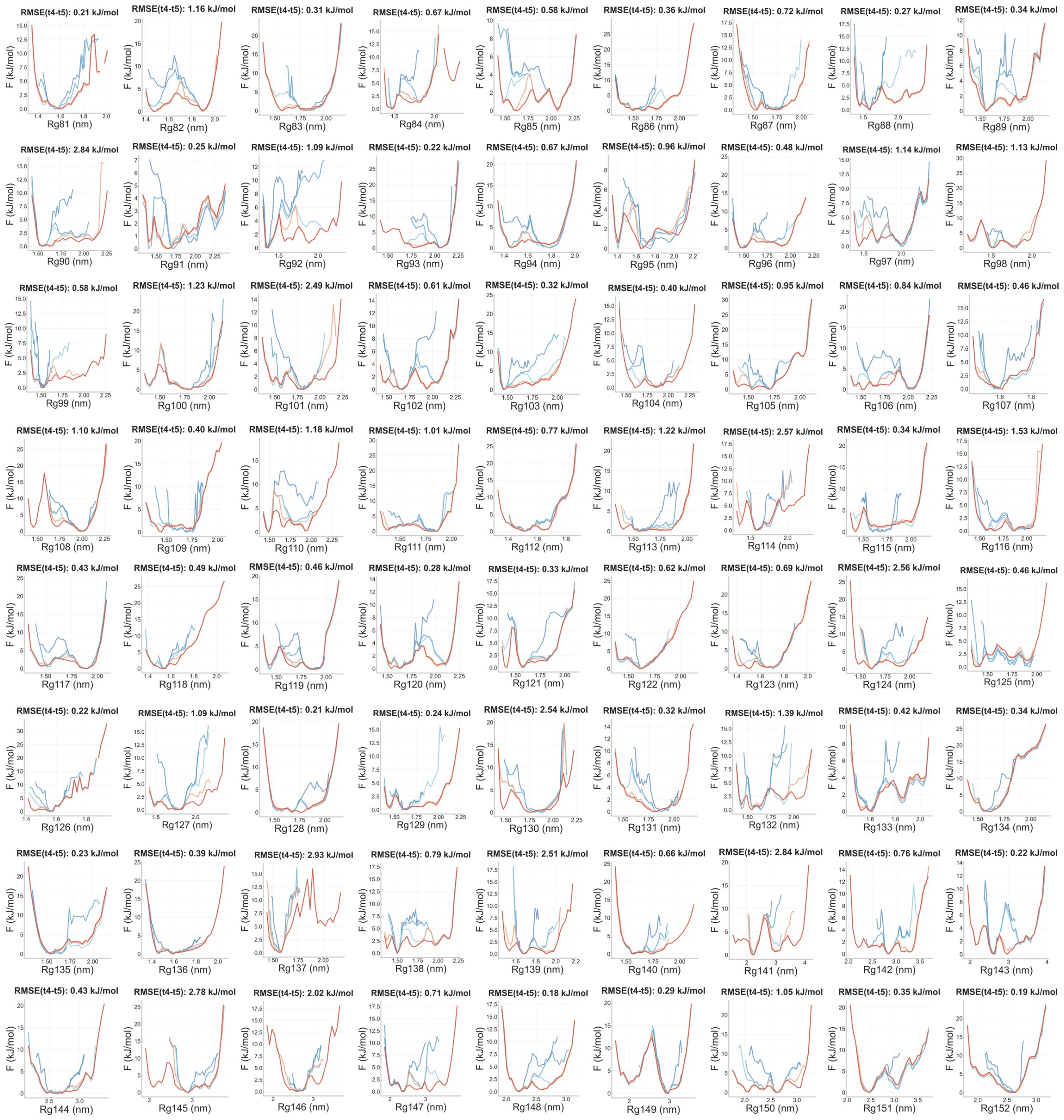


**Extended Data Figure 51: Assessment of the convergence of the MEMMI simulations.**

Free energy profiles for all biased CVs. Dark purple, light purple, cyan, orange, and red correspond to FES projections using data of 20%, 40%, 60%, 80%, 100% coverage of the concatenated MEMMI trajectory. The Root Mean Square Error quantifies the convergence of the FES as a function of time and corresponds to the difference between the FES projections at 80% and 100%.

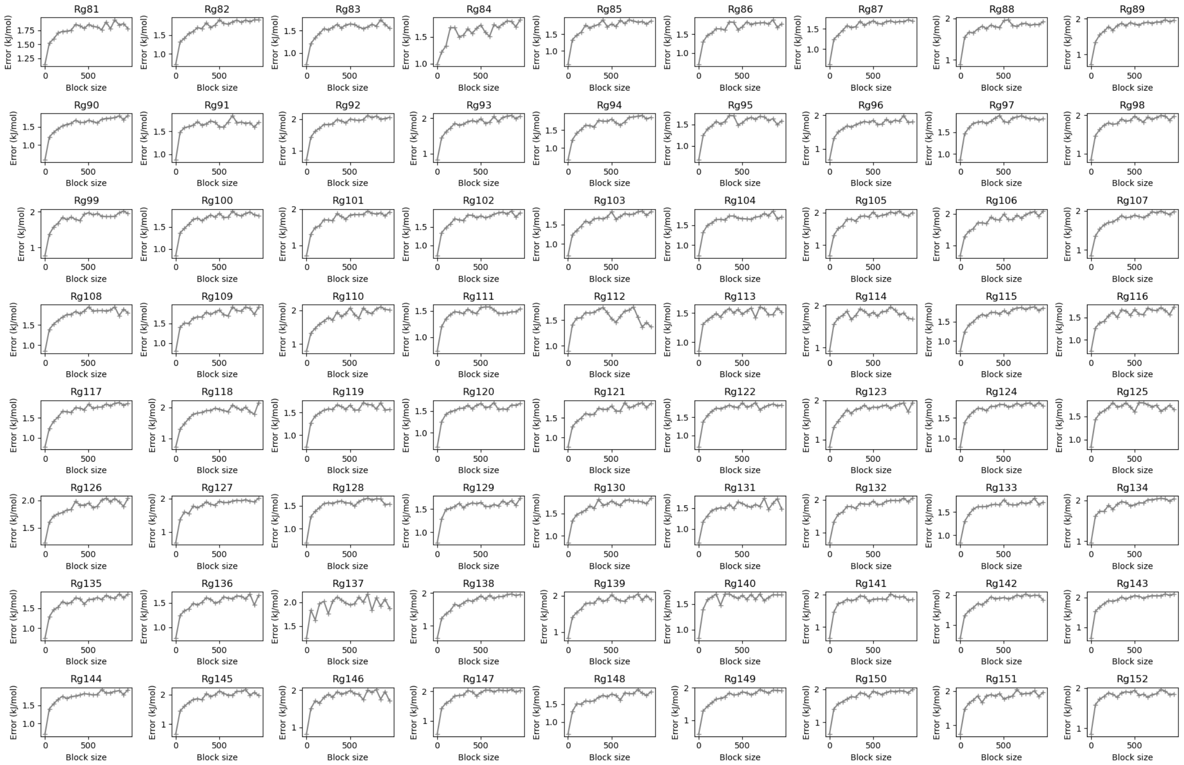


**Extended Data Figure 52: Assessment of the convergence of the MEMMI simulations via block analysis for each CV trace.**

Each plot illustrates the average error along the free-energy profile of each CVs as a function of the block length.

### Extended Data Tables

#### Extended Data Table 1: Comparison of published PDH E2 structures.

PDB ID, biological source, expression system, endogenous expression if applicable, the achieved resolution, and experimental methodology are listed.

| ID | Origin | Expression | Endog. | Res.[Å] | Method | Cofactors |
|---|---|---|---|---|---|---|
| 6CT0 | *H. sapiens* | *E. coli* | no | 3.1 | SPA | / |
| 6ZLO | *N. crassa* | *E. coli* | no | 2.9 | SPA | / |
| 7R5M | *N. crassa* | *E. coli* | no | 3.3 | SPA | / |
| 8OHS | *N.crassa* | *N. crassa* | no | 4.1 | SPA | / |
| 8PIU | *H. sapiens* | *E. coli* | no | 2.9 | SPA | / |
| 3B8K | *H. sapiens* | *E. coli* | no | 8.8 | SPA | / |
| 1B5S | *G.stearothermophilus* | *G. stearothermophilus* | no | 4.4 | X-Ray | / |
| 1DPB | *A. vinelandii* | *A. vinelandii* | no | 2.5 | X-Ray | / |
| 7B9K | *E. coli* | *E. coli* | yes | 3.2 | SPA | LA2 |
| 8OSY | *E. coli* | *E. coli* | no | 1.9 | X-Ray | / |
| 7UOM | *B. taurus* | *B. taurus* | yes | 3.8 | SPA | / |
| 6H55 | *H. sapiens* | *E. coli* | no | 6.0 | SPA | / |
| 6ZZI | *C.glutamicum* | *E. coli* | no | 1.9 | X-Ray | / |
| 4N72 | *E. coli* | *E. coli* | no | 2.3 | X-Ray | / |
| 6ZZL | *C. glutamicum* | *E. coli* | no | 2.2 | X-Ray | PO4,GOL |
| 1EAB | *E. coli* | *E. coli* | no | 2.6 | X-Ray | **CoA**, LPM |
| 1DPC | *A. vinelandii* | *A. vinelandii* | no | 2.6 | X-Ray | / |
| 1DPD | *A. vinelandii* | *A. vinelandii* | no | 2.7 | X-Ray | / |
| 7UOL | *B. taurus* | *B. taurus* | yes | 3.5 | SPA | / |
| 6ZZM | *C. mustelae* | *E. coli* | no | 2.5 | X-Ray | **CoA** |
| 7R5M | *N. crassa* | *E. coli* | no | 3.3 | SPA | / |
| 7BGJ | *T. thermophila .* | *T.thermophila* | yes | 6.9 | SPA | / |
| 7OTT | *T.thermophila .* | *T. thermophila* | yes | 3.8 | SPA | / |
| 6ZZL | *C.glutamicum .* | *E. coli* | no | 2.2 | X-Ray | PO4,GOL |
| 6ZZJ | *C.glutamicum .* | *E. coli* | no | 1.4 | X-Ray | CAO, EPE |
| 8OSY | *E. coli* | *E. coli* | no | 1.9 | X-Ray | / |
| **PDBID** | ***Th. thermophila .*** | ***T.thermophila*** | **Yes** | **2.8** | **SPA** | **CoA** |

**Extended Data Table 2: Reconstruction and refinement statistics of the SPA reconstructions.**

| | E2 CD GS (PDB-29OF ) (EMD-57265) | E2 CD Saturated (PDB-29OE ) (EMD-57264) | E3BP CBD (PDB- 29OG) (EMD-57266) | E1 (PDB-29OD) (EMD-57263) |
|---|---|---|---|---|
| **Data collection and processing** | | | | |
| Voltage (kV) | 300 | 200 | 300 | 300 |
| Microscope model | TFS Krios | TFS Glacios | TFS Krios | TFS Krios |
| Camera Model | Gatan K3 | Falcon 4i | Gatan K3 | Gatan K3 |
| Electron exposure (e-/Å^2) | 30 | 30 | 30 | 30 |
| Defocus range (μm) | - 3 to - 1 | - 3 to - 0.8 | - 3 to - 1 | - 3 to - 1 |
| Pixel size (Å) | 0.68 | 1.53 | 0.68 | 0.68 |
| Images (number aquired) | 19049 | 35668 | 19049 | 19049 |
| Aquisition software | TFS EPU | TFS EPU | TFS EPU | TFS EPU |
| Symmetry imposed | I | I | T | C2 |
| Final particle images (number) | 421139 | 122683 | 95365 | 75015 |
| Map resolution (Å) | 2.8 | 3.1 | 3.1 | 5.8 |
| FSC treshold | 0.143 | 0.143 | 0.143 | 0.143 |
| Map B-factor | 167.0 | 160.4 | 120.2 | 704.3 |
| **Refinement** | | | | |
| Initial model used | 0 (not modified) | | | |
| Map sharpening B factor (Å^2) | 0 (not modified) | | | |
| Model composition | | | | |
| Chains | 1 | 1 | 1 | 2 |
| Atoms | 1765 | 1831 | 991 | 5530 |
| Residues | 232 | 235 | 126 | 718 |
| Ligands | 0 | 1 | 0 | 0 |
| Water | 0 | 0 | 0 | 0 |
| **Bonds (RMSD)** | | | | |
| Length (Å) (# > 4 σ) | 0 | 0.007 | 0.006 | 0.003 |
| Angles (°) (# > 4 σ) | 0 | 1.061 | 1.076 | 0.655 |
| **MolProbity score** | 1.18 | 1.36 | 1.86 | 2.03 |
| **Clash score** | 3.90 | 6.48 | 18.70 | 13.78 |
| **Ramachandran plot (%)** | | | | |
| Outliers | 0 | 0 | 0 | 0 |
| Allowed | 0.43 | 1.29 | 2.46 | 5.60 |
| Favoured | 99.57 | 98.71 | 97.54 | 94.40 |
| **Rama-Z (Ramachandran plot Z-Score, RMSD)** | | | | |
| whole (N=2060) | 1.69 (0.54) | 1.62 (0.54) | -1.11 (0.77) | -0.74 (0.31) |
| helix (N=748) | 3.08 (0.54) | 2.15 (0.49) | -1.60 (0.58) | 0.13 (0.30) |
| sheet (N=368) | 1.06 (0.85) | 0.65 (0.88) | 0.40 (1.24) | -1.64 (0.72) |
| loop (N=944) | -0.45 (0.54) | 0.47 (0.62) | 0.67 (1.14) | -0.76 (0.33) |
| **Rotamer outliers (%)** | 0 | 0 | 0 | 0 |
| **Cβ outlier (%)** | 0 | 0 | 0 | 0 |
| **Peptide plane (%)** | | | | |
| Cis proline/general | 9.1/0.0 | 9.1/0.0 | 0.0/0.0 | 0.0/0.0 |
| Twisted proline/general | 0.0/0.0 | 0.0/0.0 | 0.0/0.0 | 0.0/0.0 |
| **CaBLAM outliers (%)** | 0.44 | 0.43 | 0.85 | 3.38 |
| **Model vs. Data** | | | | |

| | | | | |
|---|---|---|---|---|
| CC (mask) | 0.83 | 0.86 | 0.73 | 0.62 |
| CC (box) | 0.35 | 0.34 | 0.43 | 0.41 |
| CC (volume) | 0.75 | 0.84 | 0.73 | 0.60 |
| CC (peaks) | 0.13 | 0.03 | 0 | 0.06 |
| CC (main chain) | 0.81 | 0.83 | 0.74 | 0.61 |
| CC (side chain) | 0.77 | 0.79 | 0.70 | 0.56 |

**Extended Data Table 3: Compositional statistics of the integrative PDHc model.**
The overall amino acid composition of the integrative all-atom model was analyzed.

| AA Composition | | |
|---|---|---|
| Amino Acid | Count | Percentage |
| A | 8306 | 11.75 |
| C | 444 | 0.63 |
| D | 3204 | 4.53 |
| E | 5896 | 8.34 |
| F | 2160 | 3.06 |
| G | 6136 | 8.68 |
| H | 948 | 1.34 |
| I | 4048 | 5.73 |
| K | 5342 | 7.56 |
| L | 5232 | 7.40 |
| M | 1576 | 2.23 |
| N | 2256 | 3.19 |
| P | 4112 | 5.82 |
| Q | 2072 | 2.93 |
| R | 2480 | 3.51 |
| S | 3496 | 4.95 |
| T | 4540 | 6.42 |
| V | 5884 | 8.32 |
| W | 500 | 0.71 |
| Y | 2052 | 2.90 |
| Number of residues | **70684** | |
| Atoms (non protonated) | 534044 | |
| Atoms (protonated) | 1073192 | |
| neg. charged AA | 9100 | |
| pos. charged AA | 8770 | |
| Theoretical pI | 5.5 | |

**Extended Data Table 4: Mapping of PTMs for the E1α.**
PTMs were based on conservation with yeast and human orthologs and validated for the fungal sequence.

| Subunit | Source | Mapped res. | Modification |
|---|---|---|---|
| E1α | human | K73 | AC,Ub |
| E1α | human | K168 | Ub |
| E1α | human | Y217 | P |
| E1α | human | T221 | P |
| E1α | human | S222 | P |
| E1α | human | Y232 | P |
| E1α | human | K234 | AC,Suc,Ub |
| E1α | human | Y279 | P |
| E1α | human | S283 | P |
| E1α | human | S285 | P |
| E1α | human | T291 | P |
| E1α | human | K311 | AC,Ub |
| E1α | human | Y320 | P |
| E1α | human | T322 | P |
| E1α | human | S331 | P |
| E1α | human | K326 | AC,Suc,Ub |
| E1α | yeast | S47 | P |
| E1α | yeast | Y101 | P |
| E1α | yeast | T166 | P |
| E1α | yeast | T250 | P |
| E1α | yeast | S255 | P |
| E1α | yeast | Y308 | P |
| E1α | yeast | S312 | P |
| E1α | yeast | S314 | P |
| E1α | yeast | S331 | P |
| E1α | yeast | Y398 | P |

**Extended Data Table 5: Mapping of PTMs for the E1β.**
PTMs were based on conservation with yeast and human orthologs and validated for the fungal sequence.

| Subunit | Source | Mapped res. | Modification |
|---|---|---|---|
| E1β | human | K75 | AC,Ub,Sum |
| E1β | human | Y86 | P |
| E1β | human | Y90 | P |
| E1β | human | K91 | AC,Ub |
| E1β | human | K207 | Ub |
| E1β | human | K211 | Ub |
| E1β | human | K250 | Ac |
| E1β | human | K359 | Ub |
| E1β | human | K359 | AC |
| E1β | human | K332 | AC,Ub |

**Extended Data Table 6: Mapping of PTMs for the E2.**
PTMs were based on conservation with yeast and human orthologs and validated for the fungal sequence.

| Subunit | Source | Mapped res. | Modification |
|---|---|---|---|
| E2 | human | S43 | P |
| E2 | human | K183 | AC,Ub |
| E2 | human | K197 | AC,Ub |
| E2 | human | K281 | AC,Ub |
| E2 | human | S283 | P |
| E2 | human | K290 | Ub |
| E2 | human | K355 | Ac,SucUb |
| E2 | human | K360 | AC |
| E2 | human | K448 | AC |
| E2 | yeast | S153 | P |
| E2 | yeast | T154 | P |
| E2 | yeast | S155 | P |
| E2 | yeast | S164 | P |
| E2 | yeast | S179 | P |
| E2 | yeast | S215 | P |
| E2 | yeast | S247 | P |
| E2 | yeast | T257 | P |
| E2 | yeast | S258 | P |
| E2 | yeast | Y280 | P |

##### Extended Data Table 7: Mapping of PTMs for the E3.

PTMs were based on conservation with yeast and human orthologs and validated for the fungal sequence.

| Subunit | Source | Mapped res. | Modification |
|---|---|---|---|
| E3 | human | K70 | AC |
| E3 | human | K106 | AC,Ub,Suc |
| E3 | human | K229 | AC,Ub,Suc |
| E3 | human | K236 | AC,Suc |
| E3 | human | K240 | AC,Ub |
| E3 | human | T247 | P |
| E3 | human | K316 | Ub |
| E3 | human | R296 | AC |
| E3 | human | K342 | AC,Ub |
| E3 | human | K372 | AC,Suc,Ub |
| E3 | human | K426 | Ac,Suc,Ub |
| E3 | human | Y498 | P |
| E3 | human | K500 | AC,Suc,Ub |
| E3 | yeast | S182 | P |
| E3 | yeast | S205 | P |
| E3 | yeast | T341 | P |
| E3 | yeast | S423 | P |
| E3 | yeast | T427 | P |

##### Extended Data Table 8: Mapping of PTMs for the E3BP.

PTMs were based on conservation with yeast and human orthologs and validated for the fungal sequence.

| Subunit | Source | Mapped res. | Modification |
|---|---|---|---|
| E3BP | human | K60 | Ub |
| E3BP | human | K101 | Ub |
| E3BP | human | K121 | AC,Sumo |
| E3BP | human | S161 | P |
| E3BP | human | K185 | AC |
| E3BP | human | K208 | Ub |
| E3BP | yeast | S46 | P |
| E3BP | yeast | T120 | P |
| E3BP | yeast | T159 | P |
| E3BP | yeast | S161 | P |
| E3BP | yeast | T159 | P |
| E3BP | yeast | S161 | P |

**Supplementary Table 9: Vitrification parameters.**

| ID | Dataset | Conc. | Grid type | Temp. | BT | BF | Comment |
|---|---|---|---|---|---|---|---|
| 1 | Glacios PDHc GS | 1 g/l | Quantifoil® type R2/1 holey carbon-coated support films on copper 200 mesh | 4 °C | 4 s | - 1 | / |
| 2 | Glacios PDHc substrates | 1 g/l | Quantifoil® type R2/1 holey carbon-coated support films on copper 200 mesh | 4 °C | 4 s | - 1 | Sample was incubated with 2 mM ThDP, 4 mM Pyr, 3 mM $NAD^+$, 0.4 mM CoA for 3 min and directly vitrified |
| 3 | Krios PDHc GS SPA | 1 g/l | Quantifoil® type R2/1 holey carbon-coated support films on copper 200 mesh | 4 °C | 4 s | - 1 | / |
| 4 | Krios PDHc GS Tomo | 1 g/l | Quantifoil® type R2/1 holey carbon-coated support films on copper 200 mesh | 4 °C | 4 s | - 1 | / |
| 5 | Krios *T.th* | / | Quantifoil® type R2/1 holey Au support films on Au 100 mesh (Mycelium was grown for 3 h on the grid; 1 ml of CCM was applied before blotting) | 4 °C | | - 1 | Mycelium was grown for 3 h on the grid; 1 ml of CCM was applied before blotting |

**Supplementary Table 10: Acquisition parameters.**

| ID | Dataset | Mag. | Acc. Voltage | Dose | Pixel size | Defocus range | Acquired images/tilt series | Comment |
|---|---|---|---|---|---|---|---|---|
| 1 | Glacios PDHc GS | 92 k | 200 kV | 30 e⁻/Å² | 1.53 Å | -3 to -0.8 µm | 45.780 | Falcon 4i (Glacios 200 kV) |
| 2 | Glacios PDHc substrates | 92 k | 200 kV | 30 e⁻/Å² | 1.53 Å | -3 to -0.8 µm | 35.668 | Falcon 4i (Glacios 200 kV) |
| 3 | Krios PDHc GS SPA | 64 k | 300 kV | 30 e⁻/Å² | 1.36 Å (0.68 ss) | -3 to -1 µm | 19.049 | Gatan K3 (Krios 300 kV), BioQuantum energy filter |
| 4 | Krios PDHc GS Tomo | 42 k | 300 kV | 144e⁻/Å² (total) | 2.13 Å (1.065) | -5 to -1 µm | 181 | Gatan K3 (Krios 300 kV), BioQuantum energy filter |
| 5 | Krios *T.th* | 42 k | 300 kV | 140e⁻/Å² (total) | 2.38 Å | -5 to -1 µm | 229 | Falcon 4i (Krios 300 kV), Selectris energy filter |

#### Extended Movies

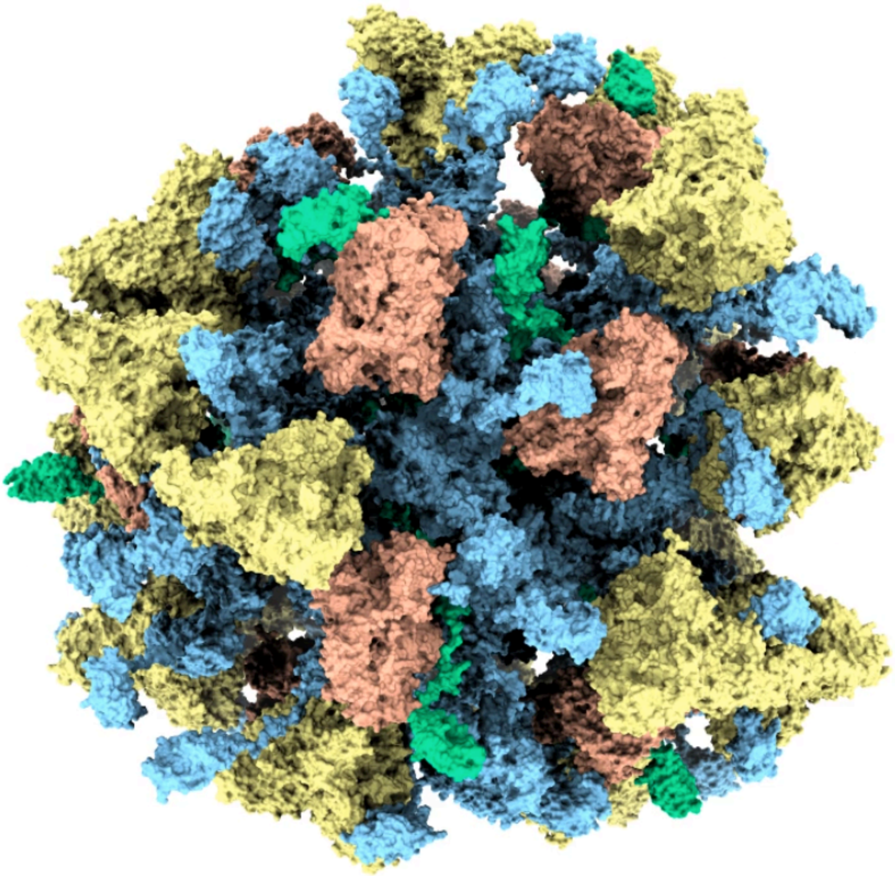

**Extended Movie 1: Simulation of the full endogenous PDHc.** The movie shows 100 selected frames from the final unbiased 500-frame MEMMI ensemble of the endogenous PDHc (E1 yellow; E2 blue; E3 orange; E3BP green). Frames were aligned on all atoms of the E2 icosahedral core. The E2 core is shown as the structural reference, while the surrounding linker regions, lipoyl domains, and peripheral E1 and E3 subunits display the conformational variability sampled by the ensemble. The movie illustrates the nested-shell organization of the PDHc and the dynamic spatial arrangement of its peripheral catalytic components around the E2 core.

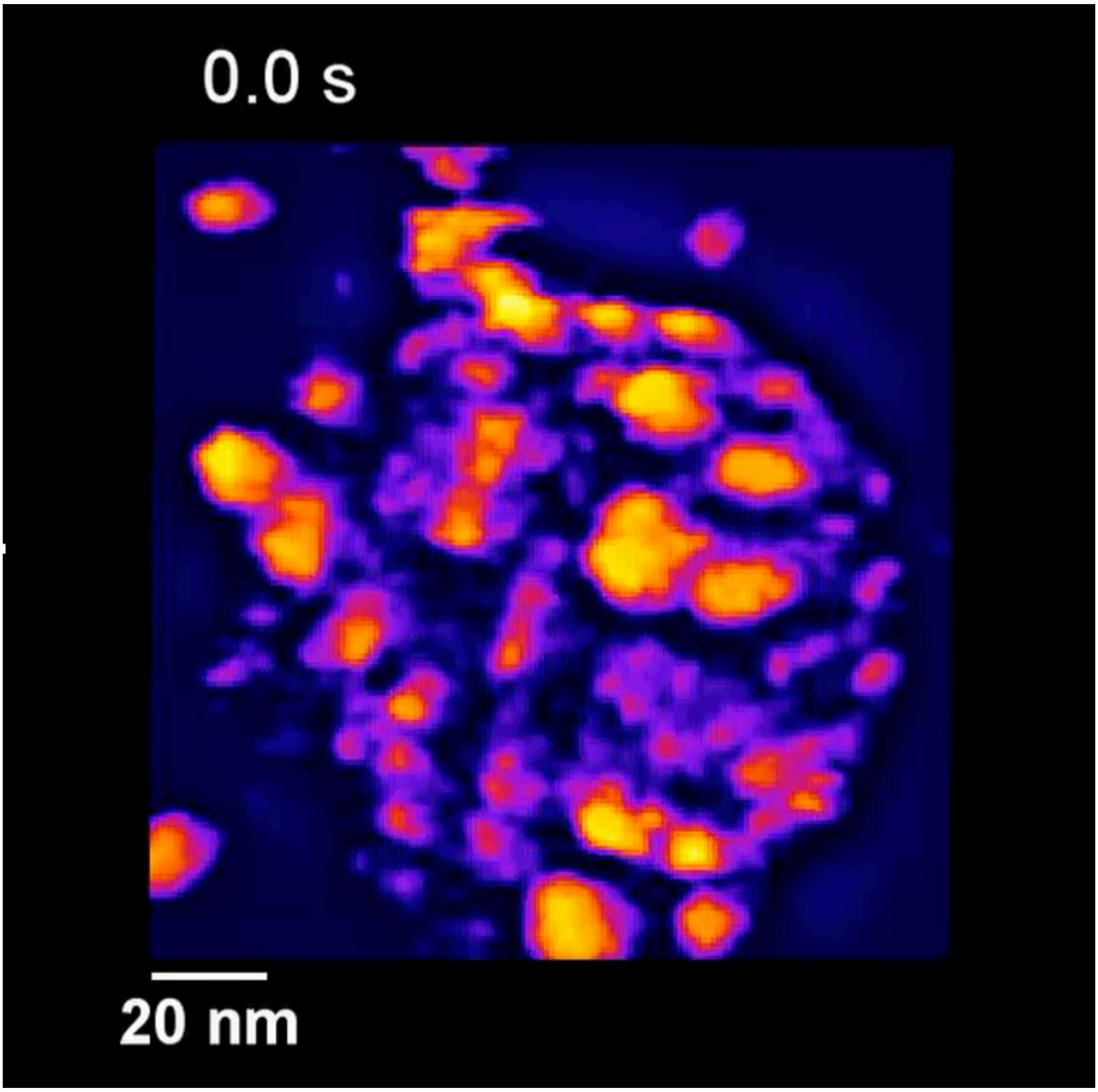


**Extended Movie 2: HS-AFM of the PDHc metabolon.** HS-AFM movie of a single surface-adsorbed endogenous PDHc. The particle appears as a large globular assembly with a heterogeneous peripheral layer surrounding a denser central region, consistent with the multidomain organization of the intact complex (imaging rate 300 ms/frame).

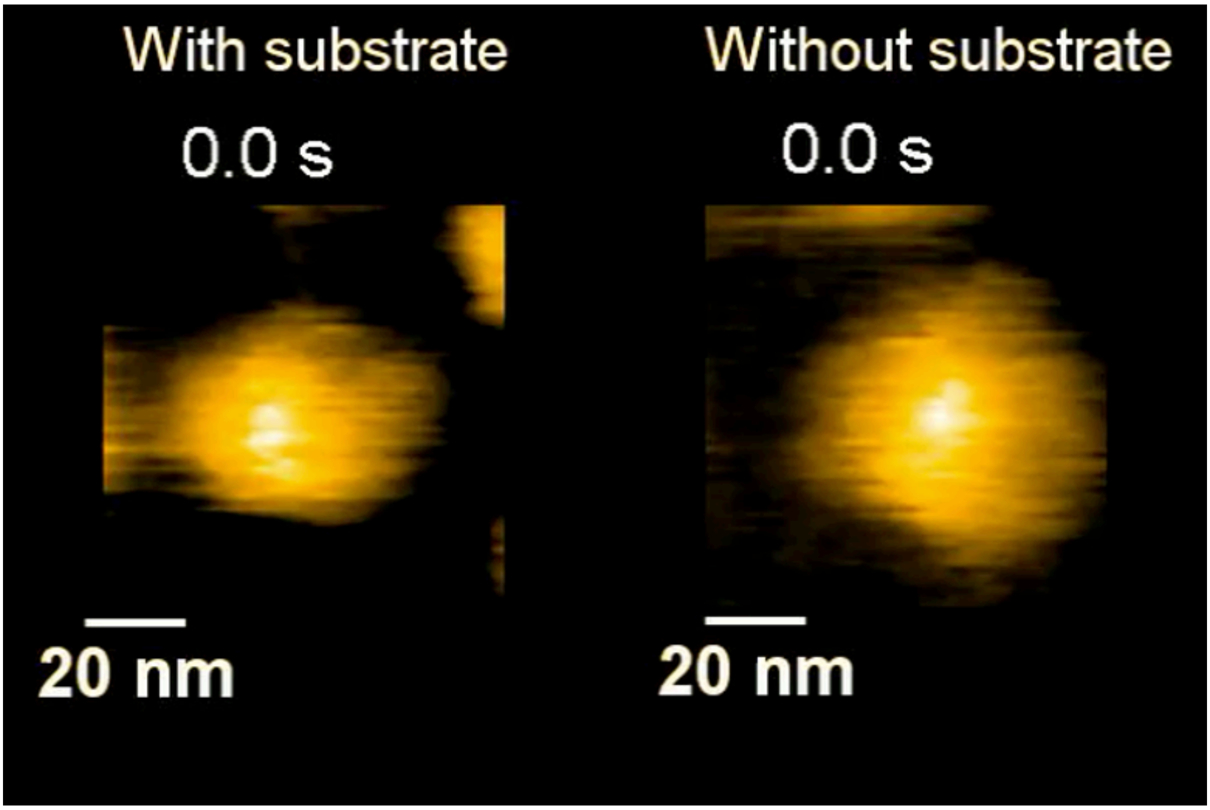


**Extended Movie 3: Mechanical Fatigue of the ground state PDHc.** Representative HS-AFM recordings of individual endogenous PDHc particles in the presence (left) and absence (right) of substrate. Comparison of the two conditions reveals differences in particle persistence and mechanical stability during repeated scanning, with substrate-bound complexes exhibiting greater structural resistance than the ground-state assembly (imaging rate 200 ms/frame (left) and 300 ms/frame (right); scale bars 20 nm).

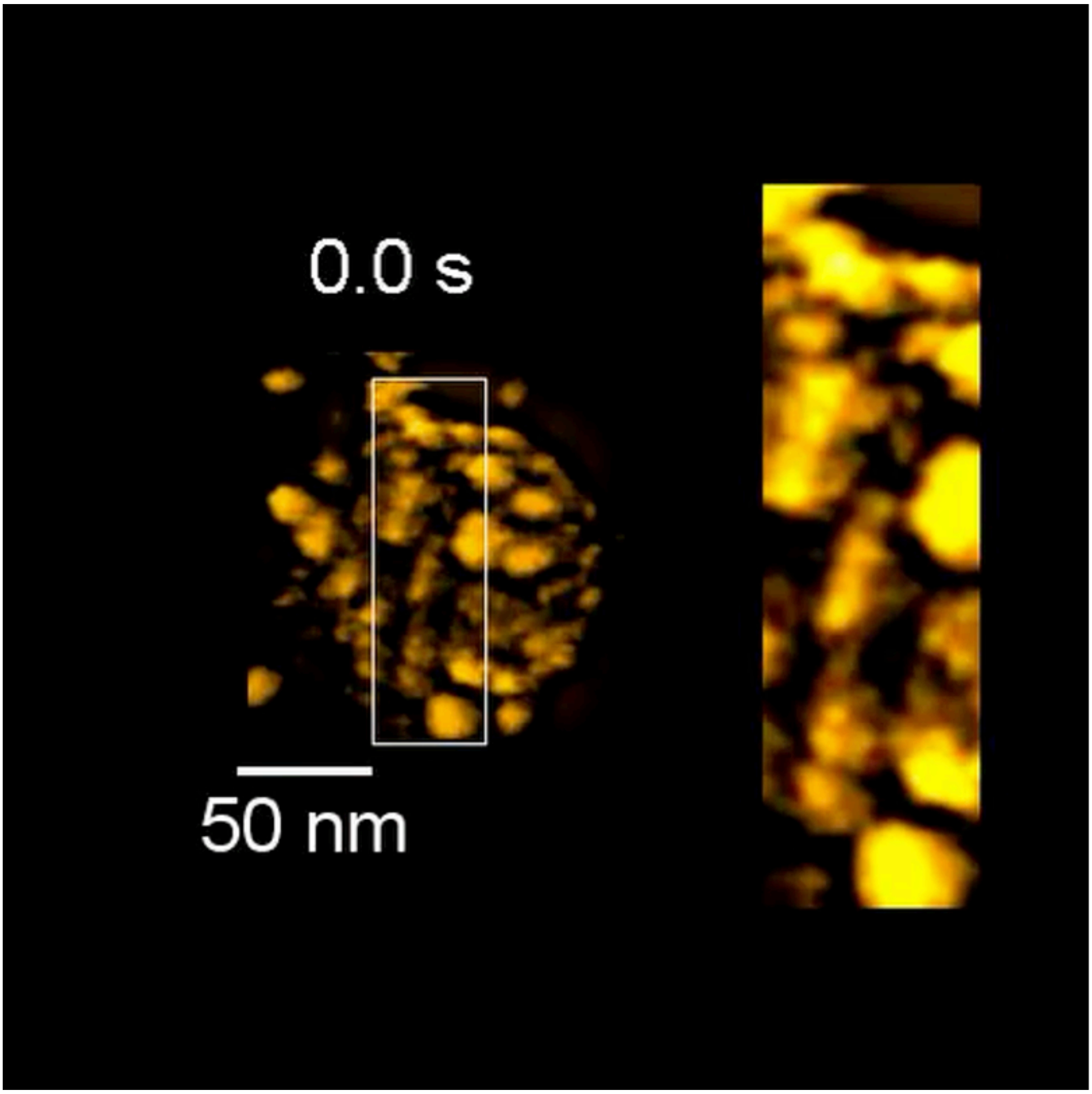


**Extended Movie 4: Subunit dynamics of PDHc visualized via HS-AFM.** HS-AFM recording of an individual endogenous PDHc particle, same as movie 2, with conventional AFM lookout color, revealing local dynamic transitions within the peripheral region of the complex. The boxed area is shown enlarged to highlight recurrent height changes of peripheral signatures. The movie captures the dynamic behavior of flexible tethered regions within the intact metabolon (imaging rate 300 ms/frame).

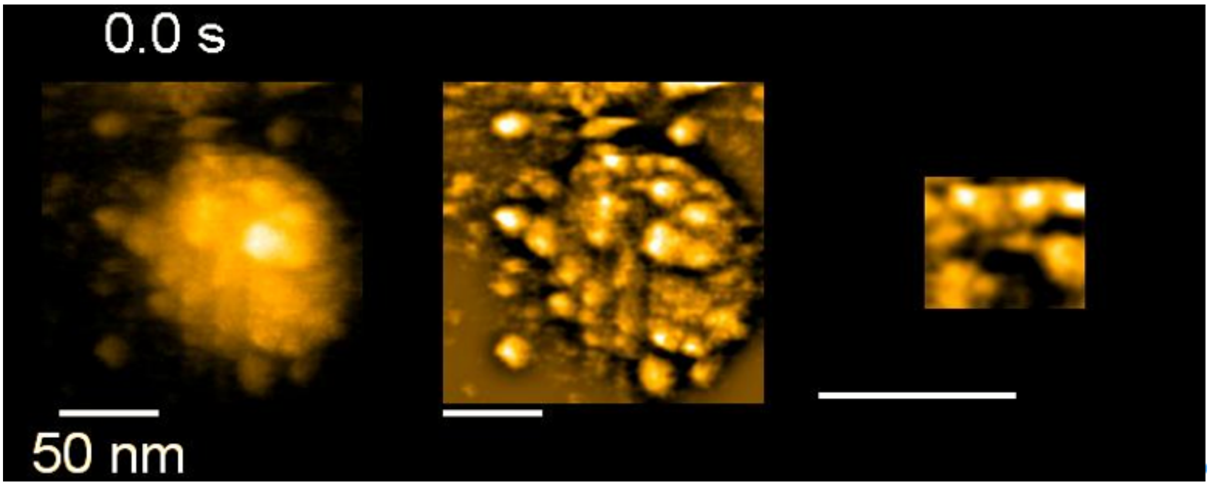


**Extended Movie 5: Subunit dynamics of PDHc visualized via HS-AFM.** HS-AFM recording of an individual endogenous PDHc particle revealing local dynamic transitions within the peripheral region of the complex. (Left) HS-AFM image, (middle) band pass filtered image, (right) zoomed in image focusing on two LDs showing axial dynamics (imaging rate 300 ms/frame, all scales 50 nm).
